# Within-Arctic horizontal gene transfer as a driver of convergent evolution in distantly related microalgae

**DOI:** 10.1101/2021.07.31.454568

**Authors:** Richard G. Dorrell, Alan Kuo, Zoltan Füssy, Elisabeth Richardson, Asaf Salamov, Nikola Zarevski, Nastasia J. Freyria, Federico M. Ibarbalz, Jerry Jenkins, Juan Jose Pierella Karlusich, Andrei Stecca Steindorff, Robyn E. Edgar, Lori Handley, Kathleen Lail, Anna Lipzen, Vincent Lombard, John McFarlane, Charlotte Nef, Anna M.G. Novák Vanclová, Yi Peng, Chris Plott, Marianne Potvin, Fabio Rocha Jimenez Vieira, Kerrie Barry, Joel B. Dacks, Colomban de Vargas, Bernard Henrissat, Eric Pelletier, Jeremy Schmutz, Patrick Wincker, Chris Bowler, Igor V. Grigoriev, Connie Lovejoy

## Abstract

The Arctic Ocean is being impacted by warming temperatures, increasing freshwater and highly variable ice conditions. The microalgal communities underpinning Arctic marine food webs, once thought to be dominated by diatoms, include a phylogenetically diverse range of small algal species, whose biology remains poorly understood. Here, we present genome sequences of a cryptomonad, a haptophyte, a chrysophyte, and a pelagophyte, isolated from the Arctic water column and ice. Comparing protein family distributions and sequence similarity across a densely-sampled set of algal genomes and transcriptomes, we note striking convergences in the biology of distantly related small Arctic algae, compared to non-Arctic relatives; although this convergence is largely exclusive of Arctic diatoms. Using high-throughput phylogenetic approaches, incorporating environmental sequence data from *Tara* Oceans, we demonstrate that this convergence was partly explained by horizontal gene transfers (HGT) between Arctic species, in over at least 30 other discrete gene families, and most notably in ice-binding domains (IBD). These Arctic-specific genes have been repeatedly transferred between Arctic algae, and are independent of equivalent HGTs in the Antarctic Southern Ocean. Our data provide insights into the specialized Arctic marine microbiome, and underlines the role of geographically-limited HGT as a driver of environmental adaptation in eukaryotic algae.

## Introduction

Four global marine biomes have been defined based on temperature, salinity and mixing regimes(Longhurst, 2006). The polar biome is characterized by year-round near freezing temperatures making polar oceans salinity-rather than temperature-stratified (Carmack, 2007). The Arctic Ocean stands apart from the Antarctic Southern Ocean through its greater geographical isolation (Beszczynska-Moller, Woodgate, Lee, Melling, & Karcher, 2011) and having fresher surface waters due to the inflow from large rivers. This added freshwater isolates the upper photic zone from saltier deep waters, and seasonal freezing renders the entire Arctic Basin an ice-influenced ecosystem.

The marine Arctic food web is supported by photosynthetic activity, performed by microalgae inhabiting both the water column (phytoplankton) and sea ice (sea-ice algae). These algae are phylogenetically diverse, including green algae, cryptomonads, haptophytes, dinoflagellates and ochrophytes (e.g., diatoms, pelagophytes and chrysophytes) (Fig. 1) (Dorrell et al., 2021a; Li, McLaughlin, Lovejoy, & Carmack, 2009). The lineages that comprise these algae split from one another hundreds of millions of years in the past and have acquired photosynthetic capacity through multiple chloroplast endosymbiotic events (Dorrell et al., 2021a; Strassert, Irisarri, Williams, & Burki, 2021).

**Fig. 1.**
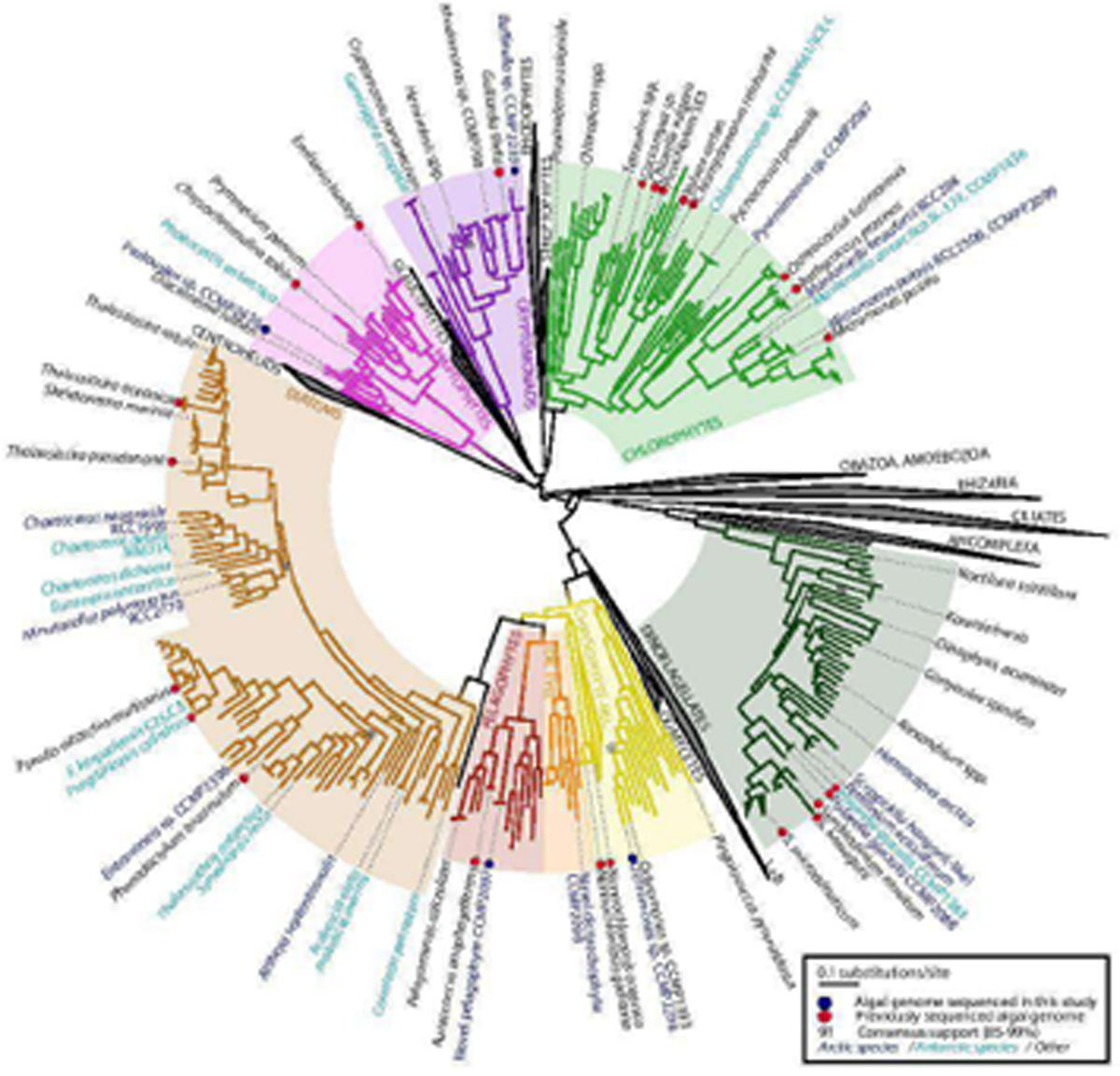
Distant evolutionary relationships of Arctic and Antarctic algae. Consensus ML topology of a 391 taxa x 39,504 aa alignment based on 250 conserved single-copy nuclear genes from across the eukaryotic tree of life (*22*); supplemented with all genomes and MMETSP transcriptomes from eight algal groups (cryptomonads, chlorophytes, chrysophytes, dictyochophytes, diatoms, dinoflagellates, haptophytes and pelagophytes) with at least one sequenced Arctic species. Branch colour corresponds to the phylogeny, and text colour the isolation site of each species considered. All sequenced algal genomes, sequenced Arctic and Antarctic algal species, and taxonomically representative taxa for each algal group are labelled. Genome libraries sequenced in this study are shown with blue circles.

Arctic-isolated algae show distinct physiological properties, in particular having viable growth temperatures that are far below those of relatives isolated from lower latitudes (Fig. 1- Figure Supplement 1) (Lovejoy et al., 2007). Different Arctic microalgae proliferate in different niches, from cold open water to ice, including microhabitats with variable salinity during ice formation and melting. During the early spring, high concentrations of nutrients in the Arctic Ocean surface facilitate diatom growth (Leu et al., 2015; Li et al., 2009; Lovejoy et al., 2007). However, from the late spring onwards, freshwater from river runoff and ice melt enforce vertical stratification of the Arctic Ocean, keeping inorganic nutrients below the photic zone. Low nutrient conditions favour smaller species with high surface-to-volume ratios and photo-mixotrophic life strategies (Bock et al., 2021; Jeong et al., 2021; Lie et al., 2018; McKie-Krisberg & Sanders, 2014). Ice-dwelling species are also exposed to salinity fluctuations, nutrient pulses and depletion, and have the ability to maintain viable cells during ice-free periods. Defining the genetic adaptations underpinning these small algal species is crucial as a baseline to understand their response to anthropogenic global change (Notz & Stroeve, 2016).

A small number of algal genomes has been previously assembled from polar biomes, including the Antarctic diatom *Fragilariopsis cylindrus*, the Antarctic chlorophytes *Chlamydomonas* sp. ICE-L and sp. UWO241, and the dinoflagellate *Polarella glacialis* (Mock et al., 2017; Stephens et al., 2020; X. Zhang, Cvetkovska, Morgan-Kiss, Hüner, & Smith, 2021; Z. Zhang et al., 2020). These libraries have yielded insights into the adaptation of algae to high latitudes, including genes encoding ice-binding proteins (IBPs) that facilitate tolerance to freezing conditions, and that have been acquired via horizontal transfers from cold-adapted bacteria (Mock et al., 2017; Raymond, 2011; Raymond & Kim, 2012; X. Zhang et al., 2021; Z. Zhang et al., 2020). We extend these insights by sequencing the genomes of four distantly related microalgae isolated from the Pikialasorsuaq/ Northwater Polynyna of the Arctic Ocean (Eegeesiak, Aariak, & Kleist, 2017). Comparing these libraries to sequenced algal genomes (Grigoriev et al., 2021) and transcriptomes from the Marine Microbial Eukaryote Transcriptome Sequencing Project (MMETSP) (Keeling et al., 2014), we reveal remarkable convergence in the coding content of these and other distantly related small Arctic algae. We further demonstrate, using genome-wide phylogenetic approaches (Dorrell et al., 2021a; Stiller et al., 2014) and environmental sequence data including *Tara* Oceans (Carradec et al., 2018; Ibarbalz et al., 2019; Pesant et al., 2015), that this is partly driven by horizontal gene transfers (HGT) between Arctic algal species. Our data reveal innovations underpinning an Arctic algal “metapan-genome”; and reposition our understanding of HGT as an effector of environmental adaptation that may even be restricted by geographic boundaries.

## Results

### New genome sequences from Arctic-isolated algae

To improve our understanding of Arctic algal evolution and adaptive diversity, genomes were sequenced from four species isolated from latitudes > 75° N. The four species are distantly related to each other (Fig. 1): *Baffinella* sp. CCMP2293, a cryptomonad, and Pavlovales sp. CCMP2436, a haptophyte, from pelagic environments; *Ochromonas* sp. CCMP2298, a chrysophyte from sea ice; and a novel pelagophyte species CCMP2097 from a brine pocket on the sea ice surface (Table S1, sheet 1) (Hamilton, Lovejoy, Galand, & Ingram, 2008; Terrado, Monier, Edgar, & Lovejoy, 2015). All four species have been demonstrated to grow optimally at low temperatures (<6 °C; Fig. 1 - Figure Supplement 1) (Keeling et al., 2014; Lovejoy et al., 2007; Terrado et al., 2015).

Phylogenetic contexts of the sequenced Arctic algae were assessed through a concatenated tree of 250 conserved single-copy nuclear genes (39,504 aa) for 391 taxa from across the eukaryotic tree of life (Fig. 1; Table S1, Sheets 2-3) (Burki et al., 2016), alongside densely sampled single-gene trees of nuclear 18S and plastid 16S rDNA (Fig. 1- Figure Supplements 2-5; Table S1, Sheets 4-7). *Baffinella* sp. CCMP2293 was resolved in an 18S analysis to a well-supported clade containing the conspecific Arctic cryptomonad *Baffinella frigidus* CCMP2045 (Fig. 1- Figure Supplement 2; (Daugbjerg, Norlin, & Lovejoy, 2018)), whereas the novel pelagophyte CCMP2097 was placed in 18S and 16S analyses near Arctic environmental isolates and members of the genus *Ankylochrysis* (Fig. 1- Figure Supplement 3) (Han et al., 2018); both were distant to other polar species with sequenced genomes or transcriptomes (Fig. 1). Pavlovales sp. CCMP2436 was placed, in all three analyses, at the base of the clade that includes the otherwise non-polar genus *Diacronema* (Bendif et al., 2011) (Fig. 1- Figure Supplement 4), while *Ochromonas* sp. CCMP2298 was found in the 18S and multigene analysis to be most closely related to the temperate species *Ochromonas* sp. CCMP1393 (Fig. 1; Fig. 1 - Figure Supplement 5) (Keeling et al., 2014; Lie et al., 2018).

Next, geographical distributions were calculated for 18S (V4 and V9 variable regions) and 16S (V4V5 regions) ribotypes from the *Tara* Oceans (including Polar Circle) Expeditions that were phylogenetically reconciled to each sequenced Arctic isolate (de Vargas et al., 2015; Ibarbalz et al., 2019; Sunagawa et al., 2015) (Fig. 2; Table S1, sheets 8-11). All four species were most frequently identified in surface water samples and 3 or 5 to 20 μm size fractions (Fig. 2- Figure Supplement 1). *Baffinella* sp. CCMP2293 and the novel pelagophyte CCMP2097 were relatively abundant, with > 10,000 total mapped ribotypes across all samples and size fractions, the majority of which were from Arctic Ocean stations (Fig. 2; Fig. 2- Figure Supplement 2 panels A, B). Pavlovales sp. CCMP2436 and *Ochromonas* sp. CCMP2298 were much rarer, with *Ochromonas* sp. CCMP2298 only detected in 16S V4-V5 ribotypes; and both were exclusively located in the Arctic (Fig. 2; Fig. 2- Figure Supplement 2 panels C, D). The relative abundances of all four species were negatively associated with temperature (Spearman *f*-test, P < 0.05) specifically in Northern hemisphere stations (Fig. 2- Figure Supplement 3; Table S1, sheet 12), suggesting cold-adaptation and predominant restriction to the Arctic. Similar results were obtained for 0.8-2000 μm size fraction samples, albeit with some presence of relatives of *Baffinella* sp. CCMP2293 and the novel pelagophyte CCMP2097 outside of the Arctic (Fig. 2- Figure Supplement 2 panels A, B).

**Fig. 2.**
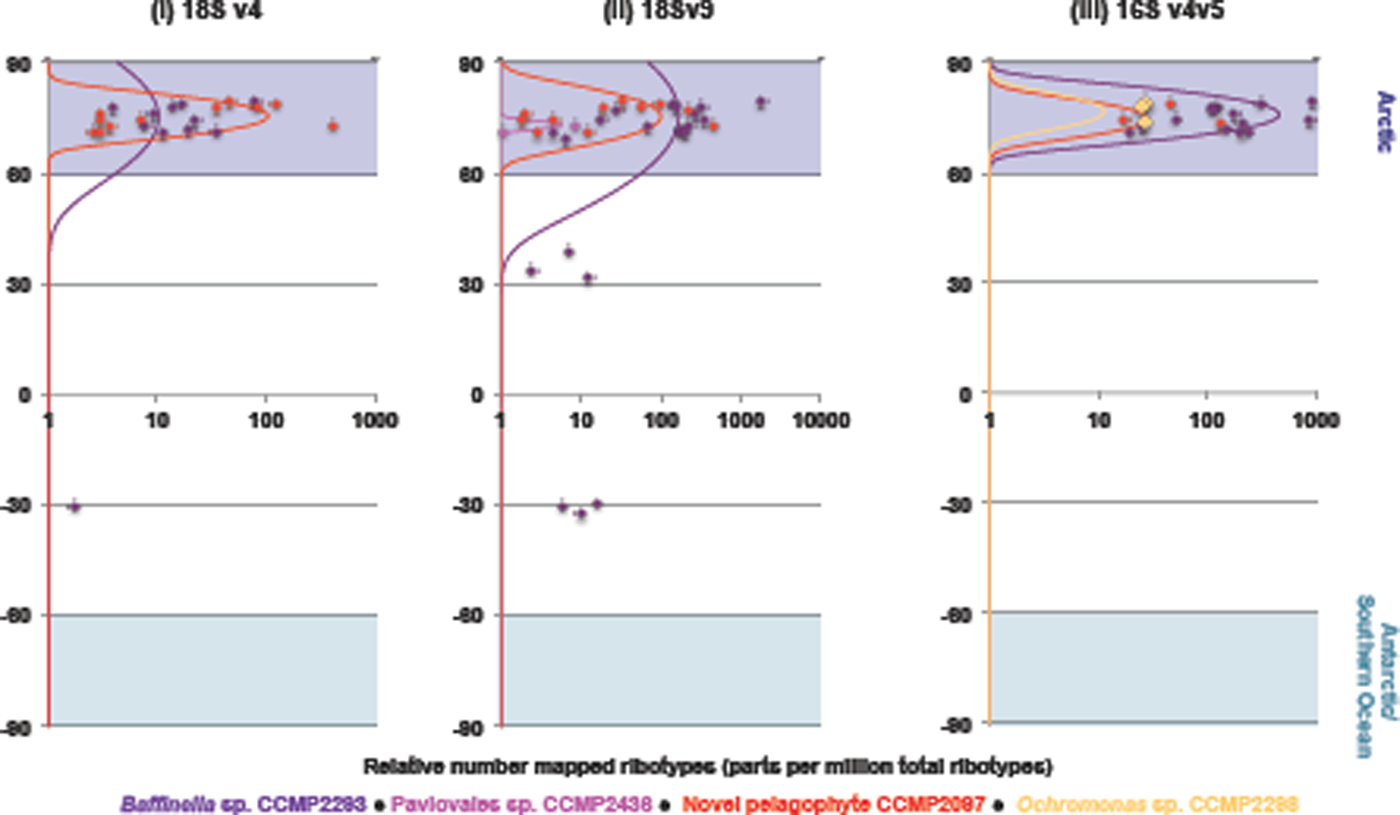
Arctic-specific distributions of sequenced algal genomes. Bell distributions of the total number of 18S V9, 18S V4, and 16S V4V5 ribotypes from *Tara* Oceans from the four sequenced algal taxa. Each point in each scatterplot corresponds to a *Tara* Oceans station, showing the relative proportion of ribotypes (expressed in parts per million total ribotypes) reconciled to each species from the surface and 3-20 μm (Arctic) or 5-20 μm (all other stations) size fractions. The lines show normal distributions of these abundances centred around the mean latitude at which each species is observed.

### Arctic algae possess expanded genomes with distinct composition

Possible genomic trends underpinning the biology of Arctic algal species were inferred across a dataset of 24 sequenced algal genomes (Grigoriev et al., 2021) and 296 MMETSP transcriptomes (Keeling et al., 2014) from eight groups (chlorophytes, chrysophyte-related species, cryptomonads, diatoms, dictyochophytes, dinoflagellates, haptophytes and pelagophytes) with at least one sequenced Arctic species, henceforth referred to as the “pan-algal dataset” (Table S1, sheet 2). The Arctic algal genomes were typically larger than sequenced non-Arctic relatives: most dramatically, the *Baffinella* sp. CCMP2293 genome (534.5 Mbp total content) is 6.13 times that of the non-Arctic cryptomonad *Guillardia theta* (87.2 Mbp; Table S2, sheet 1) (Curtis et al., 2012), although more moderate expansions (between 1.17 and 2.11 times the size of nearest sequenced relatives in the dataset) were observed for other Arctic algae. A broader analysis of the pan-algal dataset (Table S2, sheets 2, 3) revealed higher ratios of genes to PFAM protein domains (i.e., suggesting an expansion in genes without known PFAM functions), in Arctic than non-polar sequence libraries. This enrichment was statistically significant (one-way ANOVA, P < 0.05) for Arctic cryptomonads and chrysophytes compared to non-Arctic relatives (Table S2, sheet 3).

The Arctic species within the pan-algal dataset encoded greater proportions of small hydrophobic residues (alanine, glycine, and valine, one-way ANOVA, P= 0.013), and smaller proportions of charged residues (aspartate, glutamate, lysine and arginine, P= 0.032) than non-Arctic species (Table S2, sheets 4, 5). The enrichment in small hydrophobic residues was further found specifically in Arctic chrysophytes and pelagophytes compared to non-polar relatives (P < 0.05; Table S2, sheet 5).

Genome expansions have previously been identified in psychrophilic prokaryotes (Royo-Llonch et al., 2020)and eukaryotes (X. Zhang et al., 2021; Z. Zhang et al., 2020); as have expansions in hydrophobic residues in cold-adapted bacteria (Metpally & Reddy, 2009) and freshwater eukaryotic algae (Nelson et al., 2021)suggesting possible adaptive features to the cold, fresher Arctic waters.

### Convergence in protein domain content between distantly related small Arctic algae

Next, we considered the global similarity of Arctic algal genomes and transcriptomes to one another, considering PFAM domain distributions as a proxy (Fig. 3;(Mistry et al., 2020)). Phylogenetically-aware principal component analyses (phylPCA) of genomes (Fig. 3- Figure Supplement 1i) and transcriptomes (Fig. 3- Figure Supplement 1ii) revealed clusters of some Arctic species (e.g., Pavlovales sp. CCMP2436, the novel pelagophyte CCMP2097, and *Ochromonas* sp. CCMP2298), albeit with others (e.g., *Baffinella* sp. CCMP2293) forming outliers. To determine if Arctic species have converged on similar PFAM contents, pairwise Bray-Curtis indices were calculated between the PFAM distributions across the pan-algal dataset, considering genome and transcriptome libraries separately (Fig. 3- Figure Supplement 2; Table S2, sheets 6, 8-10) (Anderson et al., 2016; Horn et al., 2016; Nelson et al., 2021).To avoid artifacts caused by differences in phylogenetic proximity, species from the same taxonomic group were excluded (Fig. 3- Figure Supplement 2) prior to calculating mean Bray-Curtis values between pairs of libraries from different habitats (either: Arctic, Antarctic/ Southern Ocean, or “Other” for non-polar species).

**Fig. 3.**
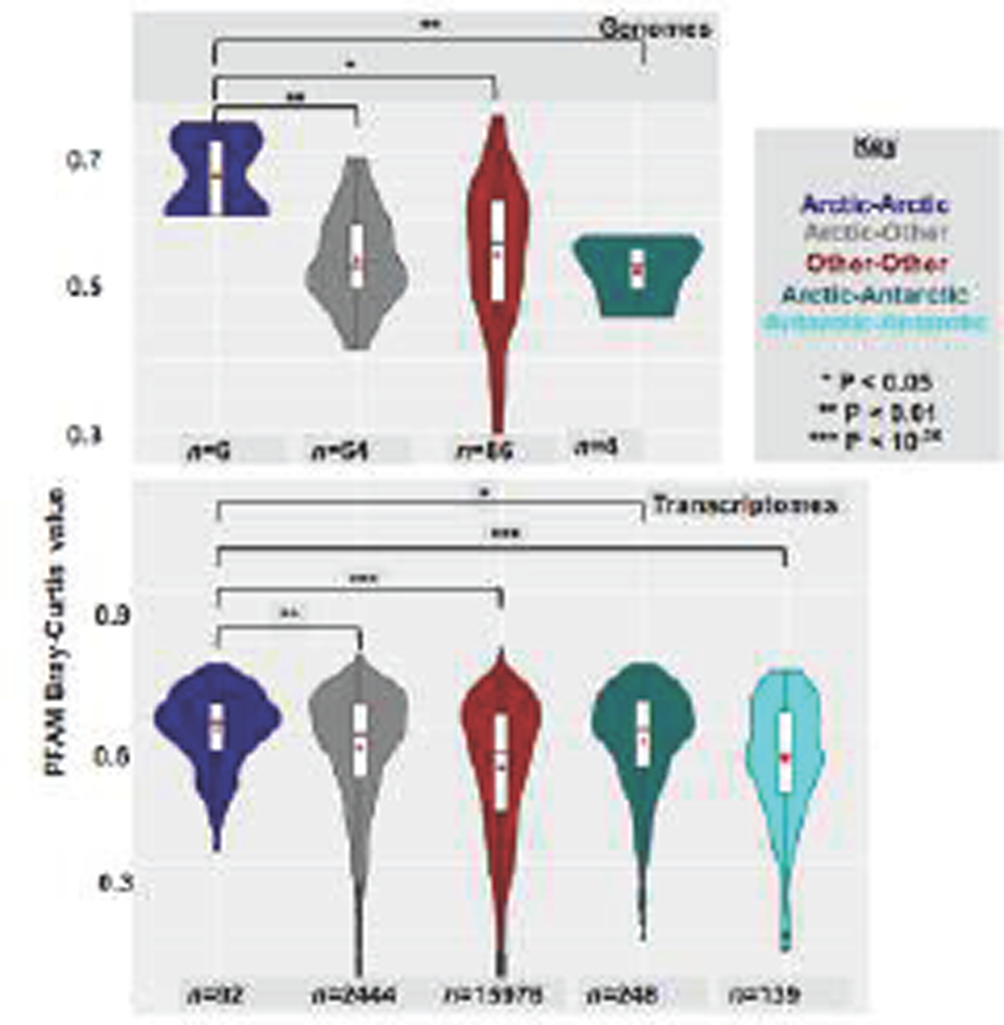
Convergence of PFAM domain contents of Arctic-specific algae. Violin plots of Bray-Curtis indices calculated between PFAM distributions of pairs of algal genomes (top) or transcriptomes (bottom), separated by habitat: Arctic (isolation site > 60°N), Antarctic (isolation site > 55°S) or Other (all intermediate latitudes). Comparisons between members of the same taxonomic group, and involving either freshwater or obligately non-photosynthetic species were excluded from the analyses. Genomic calculations involving Antarctic species are not shown due to the presence of only one Antarctic genome (*Fragilariopsis cylindrus*) in the pan-algal dataset. Significance values of one-way ANOVA tests of the separation of means (red dots) are provided between Arctic-Arctic species pairs, and all other forms of species pairs considered.

Pairs of Arctic species showed greater mean similarity in PFAM content (genome mean 0.663; transcriptome mean 0.596) than pairs of Arctic and non-polar species (genome mean 0.541; transcriptome mean 0.556); or pairs of non-polar species did to one another (genome mean 0.551; transcriptome mean 0.512; Fig. 3). The mean Bray-Curtis value observed between Arctic species pairs was significantly greater than that observed between pairs of Arctic and non-Arctic species (one-way ANOVA, genomes P = 6 x 10^-05^; transcriptomes P = 0.002); and between pairs of non-Arctic species (genomes P = 0.01; transcriptomes P = 0; Fig. 3). Greater similarity was noted between pairs of Arctic species than between Arctic and Antarctic species (e.g., *Fragilariopsis cylindrus*) (Mock et al., 2017) in both genomes (mean = 0.527, P = 0.0047) and transcriptomes data (mean 0.569; P = 0.049), indicating that this convergence is specific to the Arctic Ocean (Fig. 3). Statistically significant convergences were also observed between Arctic genomes considering the BUSCO-adjusted number of PFAMs shared between each library, suggesting it was independent of the completeness of Arctic and non-Arctic libraries in the pan-algal dataset (Fig. 3- Figure Supplement 3A); and from the Spearman rank index of PFAMs in the transcriptomes dataset (*34*) (Choi & Kim, 2007), indicating that this was not due solely to expansions in Arctic algal PFAM contents (Fig. 3- Figure Supplement 3B).

Convergences involved diverse species, with Pavlovales sp. CCMP2436 having greater convergence to the novel pelagophyte CCMP2097; and Arctic chlorophytes and dinoflagellates likewise having greater convergence to one another than to non-Arctic equivalents, for all metrics tested (Fig. 3- Figure Supplement 4; Table S2, Sheet 10). Exclusion of diatoms from the PFAM dataset indeed strengthened the separation between Arctic-Arctic species pairs and other biogeographical categories of species pairs in Bray-Curtis, BUSCO-corrected Bray-Curtis, and Spearman correlation calculations (Fig. 3- Figure Supplement 5; Table S2, sheet 10). This may point to separate trends in PFAM content between Arctic diatoms, and smaller size fraction picophytoplankton in the Arctic.

### Arctic-associated PFAMs are spread by within-Arctic horizontal gene transfer

Next, we determined which PFAMs were principally responsible for the convergence in PFAM content amongst Arctic algae (Fig. 4). We considered both occurrence in Arctic species (Table S2, sheet 6) and expansions or contractions in copy number in Arctic species compared to their closest non-Arctic relatives (Table S3, sheet 1) (X. Zhang et al., 2021), performed through a phylogenetically-calibrated CAFE (Computational Analysis of gene Family Evolution) across the pan-algal dataset (De Bie, Cristianini, Demuth, & Hahn, 2006). Multiple PFAM domains were more frequently detected, or more frequently underwent expansions (two-tailed chi-squared P < 10^-05^) in Arctic species within the dataset (Fig. 4; Table S3, sheet 2), with exemplar distributions of eight such PFAMs shown in Fig. 4- Figure Supplement 1.

**Fig. 4.**
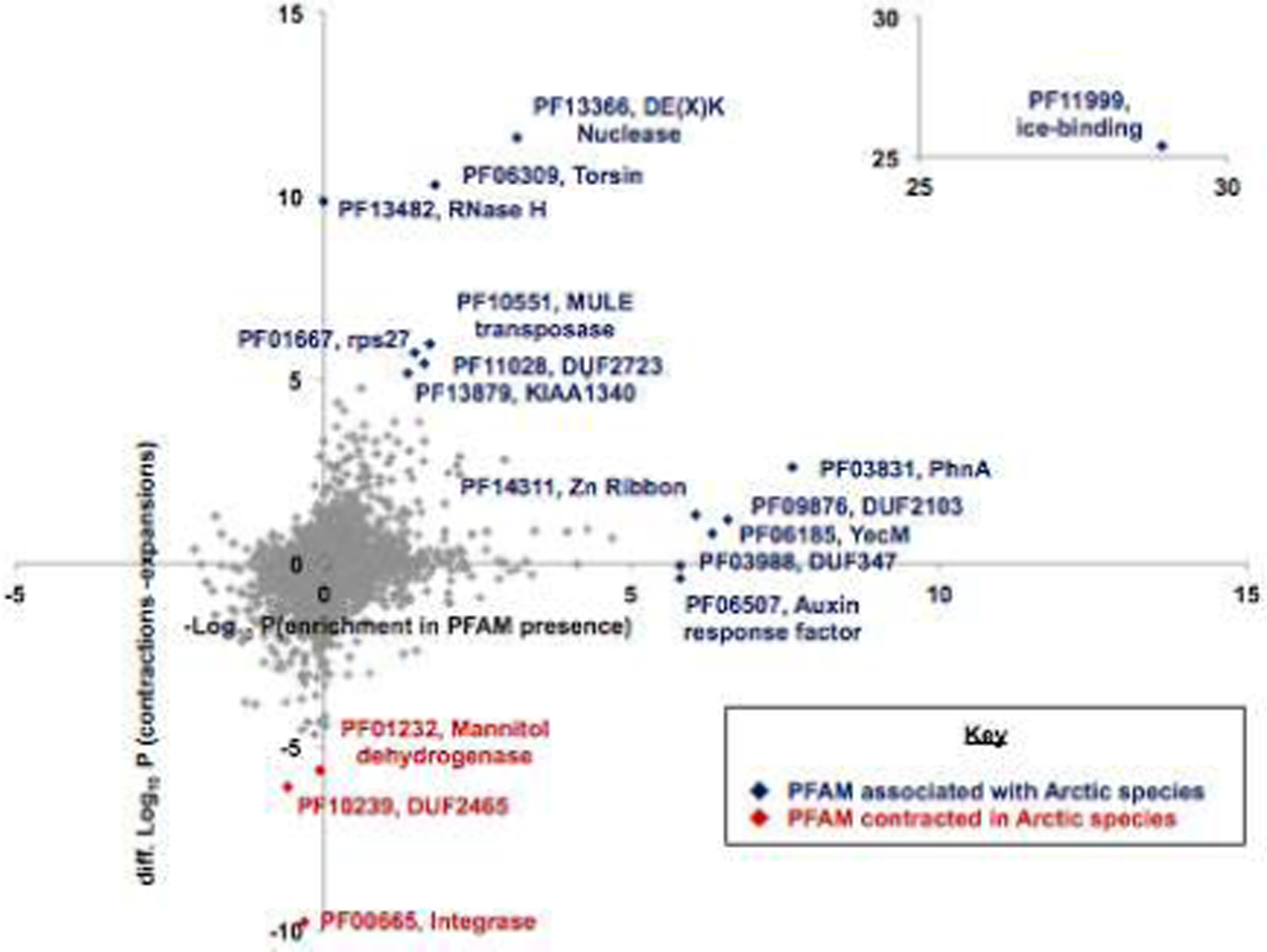
Arctic-specific expansions and contractions of PFAM domains. Scatterplot of 3,858 PFAMs, detected in at least one algal genome and one algal transcriptome, and having inferred to have undergone at least one expansion or contraction by CAFE of genome data and at least one expansion and contraction by CAFE of transcriptome data; showing possible enrichments and depletions in Arctic species. The horizontal axis shows the signed -log_10_ chi-squared P-value of the presence of PFAMs in Arctic versus non-Arctic species in the dataset. Positive values indicate the PFAM occurs more frequently than expected in Arctic species; and negative values that the PFAM occurs less frequently than expected in Arctic species. The vertical axis shows the -log_10_ chi-squared P-values for enrichment in expansions of each PFAM, inferred by CAFE, in Arctic compared to non-Arctic species, minus the –log_10_ chi-squared P-values of contractions in each PFAM in Arctic species, using the same methodology. Positive values indicate the PFAM is more frequently expanded in Arctic species and negative that it is more contracted in Arctic species than expected. PFAMs that are inferred to either be specifically associated (enriched in presence, or expanded) or not associated (contracted) in Arctic compared to non-Arctic species (P < 10^-05^) are labelled. One PFAM (PF11999, ice-binding domain) showing extreme enrichment in Arctic species is shown off-scale.

The most highly Arctic-associated PFAM domain, both considering presence and expansion frequencies (P < 10^-25^), was PF11999 (Figs. 4, 5), encoding ice-binding domains (Raymond, 2011; Raymond & Kim, 2012; Vance, Bayer-Giraldi, Davies, & Mangiagalli, 2019). Ice-binding domains, which typically complex with water through six threonine-rich domains within a larger hydrophobic manifold, play key roles in cryotolerance, variously allowing cold-adapted species to avoid osmolysis during freezing transitions within the cell, or allowing membrane surface transporters to be kept open, or even adhesion of cells to the ice surface if secreted (Raymond, 2011; Raymond & Kim, 2012; Vance et al., 2019). Ice-binding domains were accordingly detected in the overwhelming majority of the Arctic species in the dataset, with the greatest number (100 annotated in the transcriptome; 40 in the genome) in Pavlovales sp. CCMP2436 (Fig. 4- Figure Supplement 1). A broader analysis of environmental sequences containing a predicted PF11999 domain from *Tara* Oceans (Carradec et al., 2018) confirmed a strong polar signature (Fig. 5- Figure Supplement 1; Table S3, sheet 3). Nearly all of the individual *Tara* Oceans genes showed either exclusively Arctic or exclusively Antarctic distributions, with only thirty (out of 1,607 total sequences) found at intermediate latitudes, and only four distributed in both poles (Table S3, sheets 3, 6).

**Fig. 5.**
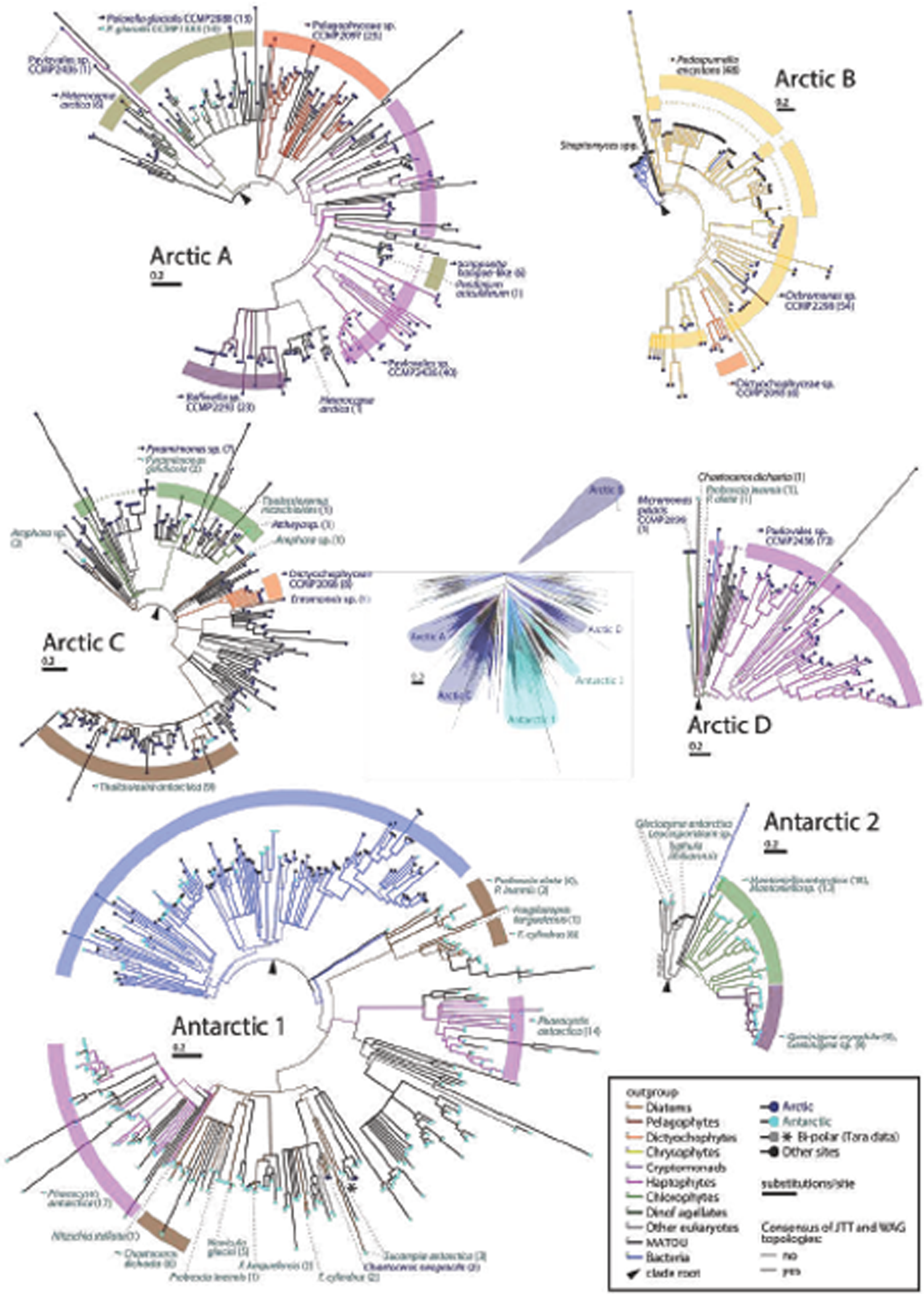
HGT of ice-binding domain sequences between Arctic algae. Consensus best-scoring tree topology obtained with RaxML under JTT and WAG substitution models for a 4862 branchx193 aa alignment of all ice-binding domain (PF11999) sampled from UniRef, JGI algal genomes, MMETSP, and *Tara* Oceans. Branches are shaded by evolutionary origin and leaf nodes by biogeography (either: isolation location of cultured accessions where recorded; or on oceanic regions for which > 70% total abundance of each *Tara* unigene could be recorded). One *Tara* unigene (asterisked) shows bipolar distributions (> 35% total abundance in both Arctic and Antarctic/ Southern Oceans). Thick branches indicate presence of a clade in both best-scoring tree outputs. The tree in the centre shows an overview of the global topology obtained; four clades of algal IBPs with probable within-Arctic transfer histories and two clades of algal IBPs with probable within-Antarctic IBPs are shown as magnified circular topologies. Numbers in parentheses identify the number of non-identical branches (i.e., gene sequences) identified in each named species. The earliest-diverging branch in each clade, relative to the remaining global tree topology, is marked with an arrow. From these rooting points, probable horizontal transfer events can be inferred e.g. from monophyletic groups of sequences, positioned within paraphyletic groups of sequences from a different phylogenetic derivation.

To investigate the evolutionary history underpinning the differential enrichment of ice-binding domains in Arctic species, we constructed a 4,862 branch tree of ice-binding domains, from all 4 query genomes, all 317 genomes and transcriptomes in the pan-algal dataset, all of UniRef (Suzek, Huang, McGarvey, Mazumder, & Wu, 2007), and all sequences from *Tara* Oceans; labelling each sequence by the phylogenetic origin of the underlying species and, where known, the geographical location (Fig. 5; Table S3, sheets 3, 7). The tree topology of ice-binding sequences did not match the underlying species phylogenies, but was separated in predominantly Arctic- and Antarctic-clades; consistent with within-ocean HGT (Fig. 5). “Arctic clade A” contains the dinoflagellate *Heterocapsa arctica*; followed by the distantly related dinoflagellate *Polarella glacialis* (Dorrell et al., 2017; Stephens et al., 2020). Of note, *P. glacialis* contains both Arctic (CCMP2088) and Antarctic (CCMP1383) isolated strains, which resolve as sister-groups (with multiple direct orthologues, (Stephens et al., 2020)). The position of both strains within an otherwise Arctic-isolated clade within the tree thus implies that *P. glacialis* was ancestrally present in the Arctic and that CCMP1383 subsequently arrived in the Antarctic (Stephens et al., 2020). Arctic clade A further contains, in probable order of acquisition: the novel pelagophyte CCMP2097; Pavlovales sp. CCMP2436; *Baffinella* sp. CCMP2293; and finally dinoflagellates within the *Scrippsiella hangoei/ Peridinium aciculiferum* species complex, which are distant from *H. arctica* or *P. glacialis* and have freshwater Arctic distributions (Craveiro, Daugbjerg, Moestrup, & Calado, 2017; Dorrell et al., 2017; Keeling et al., 2014). Similarly, Clade B contained putative transfers between *Ochromonas* sp. CCMP2298, the boreal freshwater non-photosynthetic chrysophyte *Pedospumella encystans* (Beisser et al., 2017; Dorrell et al., 2019; Grossmann, Bock, Schweikert, & Boenigk, 2016) and an Arctic dictyochophyte, CCMP2098 (Terrado et al., 2015); Clade C consisted of transfers between the Arctic dictochophyyte CCMP2098 and the chlorophyte *Pyramimonas* sp. CCMP2087 (Lovejoy et al., 2007); and Clade D consisted of HGT between Pavlovales sp. CCMP2436, the Arctic chlorophyte *Micromonas* sp. CCMP2099, and multiple Arctic *Tara* Oceans sequences (clade D) (Lovejoy et al., 2007; McKie-Krisberg & Sanders, 2014). Finally, two Antarctic-specific clades indicate parallel horizontal transfers of ice-binding proteins within Antarctic algae, specifically between the haptophyte *Phaeocystis antarctica* and multiple diatoms including *Fragilariopsis cylindrus)* (Gast, Moran, Dennett, & Caron, 2007; Keeling et al., 2014; Mock et al., 2017), with Antarctic bacteria as a probable outgroup (Brinkmeyer et al., 2003; Muñoz-Villagrán et al., 2018) (clade 1), and between the cryptomonad *Geminigera cryophila,* the chlorophyte *Mantoniella antarctica* and the Antarctic fungi *Glaciozyma antarctica* and *Leucosporidium* sp. AY30 (Lovejoy et al., 2007; Turchetti et al., 2011) (clade 2).

The topology of the IBD tree was corroborated by an internal BLAST search of the alignment to itself (Table S3, sheets 3, 8, 9), which demonstrated that (excluding hits to congeneric species, and to environmental sequence isolates) Antarctic and Arctic species typically retrieved best-scoring hits to other algae from the same oceanic region (Fig. 5- Figure Supplement 2). These included enrichments in reciprocal hits between *Heterocapsa arctica, Baffinella* sp. CCMP2293 and Pavlovales sp. CCMP2436 (Arctic clade A); *Pedospumella encystans, Ochromonas* sp. CCMP2298 and the dictyochophyte CCMP2098 (Arctic clade B); and *Mantoniella antarctica* and *Geminigera cryophila* (Antarctic clade 2).

Several additional PFAMs were significantly enriched in Arctic species (Fig. 4; Fig. 4- Figure Supplement 1). These included PF03988/ DUF347, which is likely to encode an efflux transporter implicated in metal stress responses in the diatom *Thalassiosira pseudonana* (Davis, Hildebrand, & Palenik, 2006) and the boreal actinomycete *Frankia* (Furnholm & Tisa, 2014), and previously shown to be expanded in *Scrippsiella hangoei* (Stephens, Ragan, Bhattacharya, & Chan, 2018). A second example was PF03831 (PhnA, YjdM), which functions as an uptake protein or inducer involved in alkyl-phosphonate metabolism (Chetouani, Glaser, & Kunst, 2001; Kulakova et al., 2001), and was previously shown to be expanded in bacteria from the phycosphere of Antarctic seaweeds (Cid et al., 2018). Phylogenetically and biogeographically-resolved analyses of each PFAM, using *Tara* Oceans-enriched datasets (Table S3, sheets 4-7), revealed probable HGTs between Pavlovales sp. CCMP2436 and *Scrippsiella hangoei* in the PF03988/ DUF347 tree (Fig. 5- Figure Supplement 3); and between the novel pelagophyte CCMP2097 and the Arctic dictyochophyte CCMP2098 in the PF03831/ PhnA domain tree (Fig. 5- Figure Supplement 4).

### Widespread occurrence of within-Arctic HGT across Arctic algal genomes

Next, we investigated the probable frequency of within-Arctic HGT across all available genomes of Arctic algae (Fig. 6). We adapted a protocol from previous studies based on LASTal analysis (Kiełbasa, Wan, Sato, Horton, & Frith, 2011) and linear regression (Dorrell et al., 2019; Metpally & Reddy, 2009; Stiller et al., 2014) (Fig. 6- Figure Supplement 1; Table S4, sheet 2). This involved searching two genomes from a given algal group, one Arctic and the other non-Arctic (a “*comparative pair*”) against a mixed library of Arctic and non-Arctic genomes and transcriptomes from another algal group (Fig. 6- Figure Supplement 1A). As both members of the comparative pair are of equivalent phylogenetic distance to all species within the reference library, the number of LAST best hits obtained from the Arctic and non-Arctic queries for each reference should be correlated, with deviations indicating convergences (e.g., HGT with the query species) (Fig. 6- Figure Supplement 1B).

**Fig. 6.**
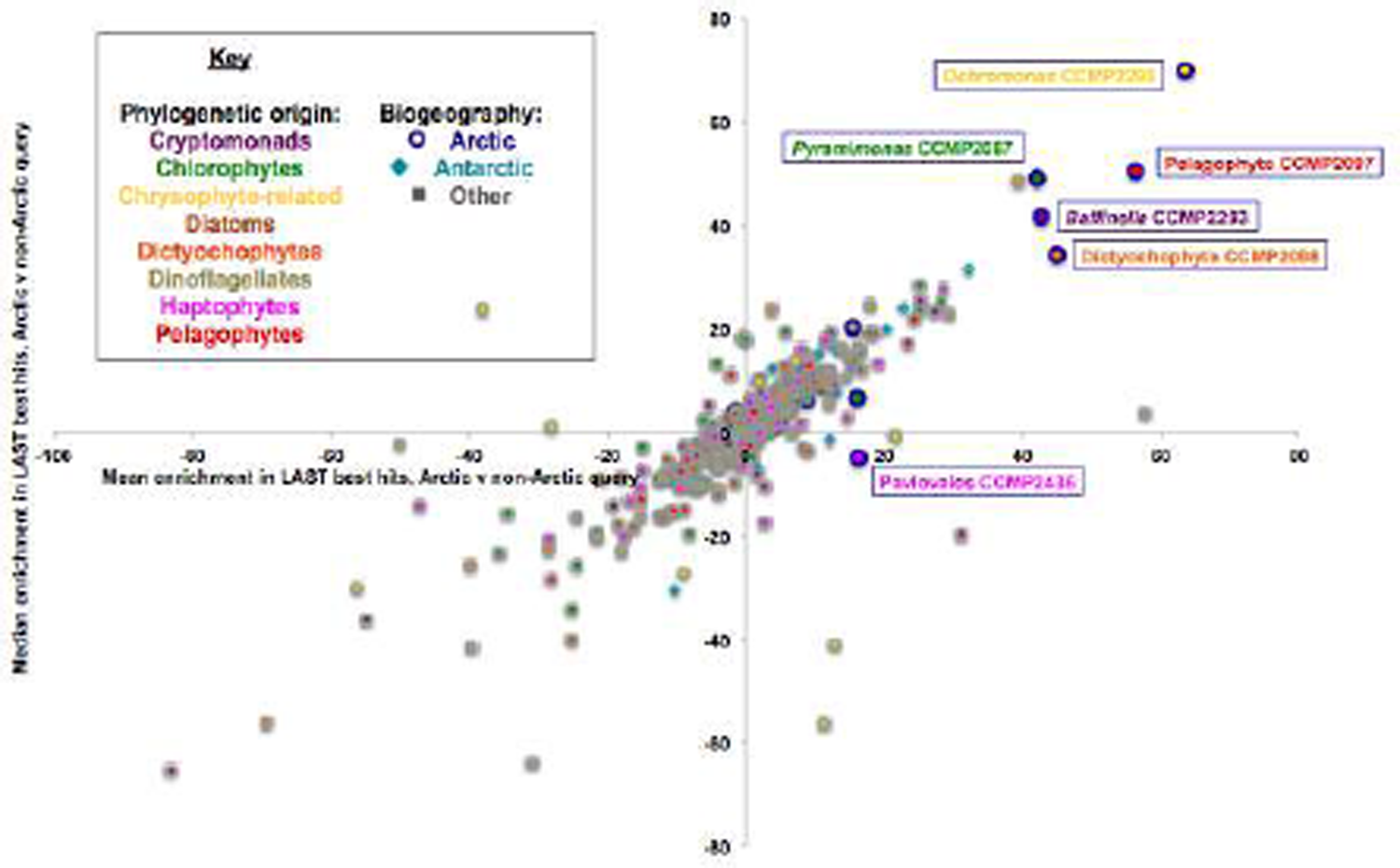
Similarity in Arctic algal genome content inferred by LAST top-hit analysis. Scatterplot showing mean (horizontal axis) and median enrichments (vertical axis) in LAST top hits from the complete genomes of *comparative pairs* of Arctic algal species and phylogenetically equivalent non-Arctic species, searched across taxon-specific reference libraries trom the pan-algal dataset. Species are shaded by phylogenetic origin (inner colour) and biogeographical origin (outer colour). Five Arctic species that show strong enrichments in LAST top-hits to other Arctic queries, plus Pavlovales sp. CCMP2436 which uniquely amongst the four sequenced species does not, are labelled. Deviation values were calculated as (CCMP2097_obs_ – CCMP2097_exp|*Aureococcus*_, CCMP2293_obs_ - CCMP2293_exp|*Guillardia*_, CCMP2436_obs_ - CCMP2436_exp*|Chrysochromulina*_, and CCMP2298_obs_ - CCMP2298_exp|*Nannochloropsis*_) where “obs” is the observed number of best hits for a given reference species and “exp” is the expected number based on linear regression from the number of best hits obtained with the non-Arctic query within the same comparative pair. Searches between query and reference libraries from the same taxonomic group were excluded from all calculations.

Deviations for multiple comparative pairs of Arctic and non-Arctic species were calculated for each species in the pan-algal dataset (Fig. 6). The five species that showed the greatest positive deviations in favour of Arctic queries (mean residual > 40, median residual > 30) were themselves small Arctic algae (the dictyochophyte CCMP2098, *Baffinella* sp. CCMP2293, *Pyramimonas* sp. CCMP2087, the pelagophyte CCMP2097, and *Ochromonas* sp. CCMP2298; Fig. 6). A further five Arctic species showed smaller but significant (chi=squared P < 0.05) enrichments in Arctic best hits, of which only one (*Entomoneis* sp.) was a diatom; whereas non-Arctic (including Antarctic) algae within the reference dataset showed smaller deviations (Fig. 6; Table S4, sheet 3).

To explicitly recover additional within-Arctic HGTs, we used genes from each sequenced Arctic genome that produced LAST best hits involving other Arctic algal species, as seed proteins for the generation of manually curated trees (Table S4, sheets 4; Fig. 6- Figure Supplement 2). Each tree was enriched with the best homologues of the query sequence from all 317 algal genomes and transcriptomes in the pan-algal dataset, a further 82 reference prokaryotic and eukaryotic genomes sampled for taxonomic representation and the inclusion of polar isolated species, and from a previously assembled dataset of 151 combined genome and transcriptome libraries covering the entire tree of life; i.e., up to 559 total homologues (Table S4, sheet 5) (Dorrell et al., 2019; Dorrell et al., 2021a).

Following manual curation (Table S4, sheets 6-8), 34 gene clusters were identified that supported within-Arctic HGTs (RAxML threshold bootstrap support 50%; Fig. 6- Figure Supplement 3; Table S4, sheet 9), including a well-supported ice-binding protein clade probably corresponding to “Arctic clade A” in the global IBD phylogeny (Fig. 4). Although a diverse range of Arctic species were detected in the within-Arctic HGT clades, including all four query genomes, the dictyochophyte CCMP2098, *Pyramimonas* sp. CCMP2087 and multiple Arctic dinoflagellates (*Heterocapsa, Scrippsiella, Peridinium* sp.), not one tree resolved HGTs between the four query species and Arctic diatoms (Table S4, sheet 9).

Several of the genes inferred to have been horizontally transferred between Arctic algae correspond to proteins which may carry out Arctic-adaptive functions (Table S4, sheet 9). These include an alcohol dehydrogenase and a CCCH Zn-finger PFAM domain shared between *Baffinella* sp. CCMP2293, Pavlovales sp. CCMP2436 and the novel pelagophyte CCMP2097, which are implicated in cold and salinity stress responses in *Arabidopsis* (Song, Liu, & Ma, 2019; Y. Wang et al., 2017); and a tellurite resistance protein, which was shared between *Baffinella sp.* CCMP2293 and *Ochromonas* sp. CCMP2298 and has been previously detected in polar bacteria (Muñoz-Villagrán et al., 2018). The within-Arctic HGT genes were significantly enriched (chi-squared P < 10^-05^) in signal peptides, consistent with endomembrane or secretory localisations, compared to other genes in the query genomes (Table 1; Fig. 6- Figure Supplement 4; Table S4, sheet 9); as has been noted in other meta-analyses of HGT in non-Arctic eukaryotic microbes (Dorrell et al., 2021a; Eme, Gentekaki, Curtis, Archibald, & Roger, 2017; Irwin, Pittis, Richards, & Keeling, 2021). Finally, considering homologues in *Tara* Oceans data, five gene families were identified in which all *Tara* Oceans genes phylogenetically reconciled to the within-Arctic HGT clade had exclusively Arctic distributions, and a further seven in which a majority of the phylogenetically reconciled *Tara* Oceans unigenes were exclusively Arctic (Fig. 6- Figure Supplement 5; Table S4, sheets 10-11); confirming their probable Arctic-specific functions.

## Discussion

We have harnessed newly sequenced genomes alongside a densely sampled dataset of genomes and transcriptomes, as well as environmental data from *Tara* Oceans to unveil evolutionary trends in algae isolated from the Arctic Ocean. Our data suggests convergence in the coding content of diverse small Arctic algae, linked to the presence of Arctic-specific genes, although with the exclusion of Arctic diatoms. We further show that within-Arctic HGT, exemplified by genes coding for ice-binding domains, is an important component of this convergent evolution. The overall results position within-ocean HGT as a mechanism of environmental adaptation, which may occur via biotic interactions characteristic of Arctic species such as photo-mixotrophy (McKie-Krisberg & Sanders, 2014; Søgaard et al., 2021) and viral infection (Irwin et al., 2021; Nelson et al., 2021); or even the direct exchange of genetic material through the Arctic water column or sea ice (Raymond, 2011; Raymond & Kim, 2012).

It remains to be determined whether within-Arctic convergences in sequenced species, which were predominantly sampled from Baffin Bay, extend to other regions within the Arctic Ocean, and algal groups. Deeper sequencing of Arctic species, either from cultured isolates, or from Arctic-specific Metagenome Assembled Genomes (Cao et al., 2020; Vorobev et al., 2020), will be instrumental in determining their broader genomic diversity. In particular, deeper sequencing of Arctic diatoms will be instrumental to understanding their different coding contents to small algal species, which may underpin their different relative seasonal niches in the Arctic (Joli, Monier, Logares, & Lovejoy, 2017; Li et al., 2009), and particularly considering the large numbers of diatom-specific HGT events visualised in previous phylogenomic studies of pan-algal HGTs (Dorrell et al., 2021a; Vancaester, Depuydt, Osuna-Cruz, & Vandepoele, 2020). Moreover, the regulation and functions of individual genes inferred to have undergone within-Arctic HGT await characterisation, e.g., using comparative transcriptomic approaches (Liang, Koester, Liefer, Irwin, & Finkel, 2019; Mock et al., 2017; Terrado et al., 2015) or through the expression and characterisation of candidate genes in transformable model algae. Finally, it remains to be determined to what extent discrete protein functional architectures have independently and convergently evolved in Arctic-, Antarctic-, and other cryophilic and ice-adapted (e.g., montane) algal species, as in the case of ice-binding domains, which may have independently originated multiple times within the tree of life (Stewart et al., 2021; Vance et al., 2019). Further functional, phylogenetic and environmental investigation of the Arctic algal metapan-genome may unveil effectors of the fitness or decline of small Arctic algae in a warming and freshening ocean environment (Li et al., 2009); and facilitate our understanding of the environmental fragility and future ecology of this important ocean biome.

## Materials and Methods

### Cultures and nucleic acid isolation

The algae were all isolated from Northern Baffin Bay within the Pikialasorsuaq/ Northwater Polynya (Eegeesiak et al., 2017)in June 1998 using a serial selection-dilution technique until a single species was isolated, and have been maintained as mono-algal cultures in L medium without Si at a salinity of 30 and ca. 4 °C and under continuous illumination since (Terrado et al., 2015). Prior to growing the sub-cultures used for DNA sequencing, the culture was transferred to L medium with added antibiotics (Terrado et al., 2015) to minimize bacterial contamination, which was assessed via light microscopy.

Nucleic acids were harvested from the batch culture in late exponential phase by centrifugation at 3000 *g* for 30 minutes at 4 °C. The supernatant was discarded and pellets were frozen in liquid nitrogen and stored at −80 °C until nucleic acid extraction. RNA and DNA was collected for whole genome sequencing following the U.S. Department of Energy Joint Genome Institute (DOE JGI) protocols (Rio et al., 2006).

### Sequencing

Genomes in this study were sequenced on an Illumina platform using a combination of a 400-800 bp Tight Insert library with 1.5 kb, 4 kb, and 8 kb insert Cre-LoxP Recombination (CLR) libraries. For the pelagophyte CCMP2097, Ligation-free paired end (LFPE) libraries were additionally used.

For Tight Insert libraries, 2-3 µg of DNA was sheared to 400 bp or 800 bp using a Covaris LE220 pulse focused sonicator (Covaris) and size selected using a Pippin Prep (Sage Science). The fragments were end-repaired, A-tailed and ligated to Illumina compatible adapters (IDT, Inc) using a KAPA-Illumina library creation kit (KAPA biosystems). For CLR libraries, 5-25 µg of DNA was sheared using a Covaris g-TUBE (Covaris) and gel size-selected for 1.5 kb, 4 kb, and 8 kb, respectively. The sheared DNA was end-repaired and ligated with biotinylated adapters containing loxP, circularized via recombination by a Cre excision reaction (NEB), and randomly sheared using a Covaris LE220 (Covaris). For LFPE libraries, 15-25 µg of DNA was sheared using HydroShear (Genomic Solutions) and gel size selected for 4.5 kb and 8 kb, respectively. The sheared DNA was end-repaired, ligated with biotinylated adapters, circularized by Intra-Molecular Hybridization, then digested with T7 Exonuclease and S1 Nuclease (Invitrogen).

Sheared ligation fragments were end-repaired and A-tailed using the KAPA-Illumina library creation kit (KAPA biosystems) followed by immobilization of mate pair fragments on streptavidin beads (Invitrogen). Illumina compatible adapters (IDT, Inc) were ligated to the mate pair fragments and amplified with 8-12 cycles of PCR (KAPA Biosystems).

All four transcriptomes were sequenced using stranded Illumina RNA-Seq protocols. Poly(A)+ RNA was isolated from 10 ug total RNA using a Dynabeads mRNA isolation kit (Invitrogen), repeated twice to remove all residual rRNA contamination; then fragmented using RNA Fragmentation Reagents (Ambion) at 70 °C for 3 min, targeting fragments around 300 bp, and purified using AMPure SPRI beads (Agencourt). Reverse transcription was performed using random hexamer primers (Fermentas) and SuperScript II (Invitrogen), with annealing, elongation and inactivation steps of 65 °C for 5 min, 42 °C for 50 min, and 70 °C for 10 min, respectively. Purified cDNA was used for second strand synthesis, with a dNTP mix where dTTP was replaced by dUTP at 16 °C for 2 h. Targeted (300 bp) double stranded cDNA fragments were purified using AMPure SPRI beads; then were blunt-ended, poly A-tailed, and ligated with TruSeq adaptors using Illumina DNA Sample Prep Kit (Illumina), and purified again. Second strand cDNA was removed through dUTP digestion with AmpErase UNG (Applied Biosystems). Digested cDNA was again cleaned up with AMPure SPRI beads, amplified by 10 cycles PCR with Illumina TruSeq primers, and finally cleaned again with AMPure SPRI beads.

Both genome and transcriptome libraries were quantified using a next-generation sequencing library qPCR kit (KAPA Biosystems) and run on a LightCycler 480 real-time PCR instrument (Roche). The quantified libraries were prepared for sequencing utilizing a TruSeq paired-end cluster kit (v3), and a cBot instrument (Illumina) to generate a clustered flow cell, and sequenced on an Illumina HiSeq2000 sequencer using a TruSeq SBS sequencing kit, v3, following a 2×100 or 2×150 indexed run recipe.

After sequencing, the genomic fastq files were screened for phix contamination. Reads composed of >95% simple sequence were removed. Illumina reads <50bp after trimming for adapter and quality (q<20) were removed. The remaining Illumina reads were assembled using AllPathsLG (Gnerre et al., 2011) parameters: DATA_SUBDIR=data RUN=run SUBDIR=assem1 TARGETS=standard OVERWRITE=True THREADS=6 CLOSE_UNIPATH_GAPS=False). The resulting assembled scaffolds were screened against all bacterial proteins and organelle sequences from GenBank nr (Wheeler et al., 2006) and removed if found to be a contaminant. Illumina transcriptome reads were QC filtered to remove contamination and assembled into consensus sequences using Rnnotator v. 2.5.3 (Martin & Wang, 2011). Each genome was annotated using the JGI Annotation Pipeline, which detects and masks repeats and transposable elements, predicts genes, characterizes each conceptually translated protein with sub-elements such as domains and signal peptides, chooses a best gene model at each locus to provide a filtered working set, clusters the filtered sets into draft gene families, ascribes functional descriptions (such as GO terms and EC numbers), and creates a JGI genome portal in PhycoCosm (https://phycocosm.jgi.doe.gov/) with tools for public access and community-driven curation of the annotation (Grigoriev et al., 2021; Kuo, Bushnell, & Grigoriev, 2014).

Residual contamination, including from Arctic bacteria, was eliminated from each genome using a custom pipeline based on phylogenetic assignation of all genes within each contig via BLAST best-hit search against a combined tree of life library, consisting of complete copies of UniRef, JGI genomes, and MMETSP algal transcriptomes (Dorrell et al., 2019; Dorrell et al., 2021a). Only genes assigned to the same contig as at least one gene of inferred vertical origin (*Baffinella* sp. CCMP2293-cryptomonads; Pavlovales sp. CCMP2436-haptophytes; *Ochromonas* sp. CCMP2298 and the novel pelagophyte CCMP2097-ochrophytes) were retained (Table S4, Sheet 1). A second round of cleaning was performed by reciprocal tBLASTn/ BLASTx searches of each peptide sequence against the corresponding MMETSP nucleotide transcriptome libraries (Keeling et al., 2014) that had been decontaminated through a previously published pairwise BLAST similarity analysis (Marron et al., 2016) (Table S4, Sheet 1). Only gene models that retrieved a reciprocal best hit with bidirectional threshold value 10^-05^ were retained for downstream analyses. Between 3,056 (*Ochromonas* sp. CCMP2298) and 11,568 gene models (Pavlovales sp. CCMP2436) were confirmed to be of unambiguously non-contaminant origin for each genome, and retained for analysis of within-Arctic HGT (Table S4, sheet 1). Full details of the decontamination pipeline employed are provided in the linked supporting database https://osf.io/3pmxb/ (Dorrell et al., 2021b)I n the folder “Within-Arctic HGTs”.

Comparisons of key properties between the assembled genomes, and their closest assembled relatives (accessed January 2018) in cryptomonads, haptophytes, pelagophytes, and chrysophyte-related species (*16*) are provided in Table S2, Sheets 1-5. Per-gene analyses of the amino acid compositions of each genome and transcriptome included in this study, as defined below, are provided in the linked supporting database https://osf.io/3pmxb/ (Dorrell et al., 2021b) in the folder “Genome quantitative analysis”.

### Multigene phylogeny

A pan-algal dataset of 21 genomes, including the four sequenced in this study (accessed January 2018) (Grigoriev et al., 2021) and 296 decontaminated MMETSP transcriptomes (Keeling et al., 2014; Marron et al., 2016) was assembled for eight algal groups for which at least one sequenced Arctic genome or transcriptome has been completed: these being chlorophytes, chrysophyte-related species (including synchromophytes, pinguiophytes and eustigmatophytes ( Dorrell et al., 2021a)), cryptomonads, diatoms, dictyochophytes, dinoflagellates, haptophytes and pelagophytes. The isolation sites of each sequenced species, and minimum and maximum recorded viable growth temperatures, were manually confirmed by comparing to the accession records in public culture collections, considering synonymous culture identifiers where present (Gachon et al., 2013; Guiry et al., 2014; Vaulot, Le Gall, Marie, Guillou, & Partensky, 2004), and are provided in Table S1, Sheet 1.

Reference sequences from a previously assembled eukaryotic multigene tree (Burki et al., 2016; Strassert et al., 2021)w ere enriched using diamond v0.9.30.131 (Buchfink, Xie, & Huson, 2015) with orthologues from all members of this dataset. To infer phylogenies, single-gene datasets were aligned by MAFFT v7.407 (Katoh, Rozewicki, & Yamada, 2017) using the L-INS-i refinement and a maximum of 1000 iterations, then trimmed by trimAl v1.4 (Capella-Gutiérrez, Silla-Martínez, & Gabaldón, 2009) using the -gt 0.5 setting. ML trees were inferred by IQ-TREE v 1.6.12 (Nguyen, Schmidt, von Haeseler, & Minh, 2015) using the LG+F+G model. Paralogs were manually removed from single-gene datasets, considering their branching and alignment coverage. The concatenated multi-gene alignment was finally trimmed to remove sites with more than 20% gaps with trimAl (-gt 0.8) and sites exhibiting the fastest exchange rates with TIGER(-exc 10 -b 10; (Cummins & McInerney, 2011)) analogously to the reference multi-gene matrix (Burki et al., 2016). The final ML tree was reconstructed using the Posterior Mean Site Frequency (PMSF) model of IQ-TREE on a LG+F+G guide tree, with the multi-gene matrix treated as a single partition to eliminate small sample LBA bias(H. C. Wang, Susko, & Roger, 2019). Incorporated gene and tree topologies are provided in Table S1, sheets 2-3; and single-gene alignments and topologies, and concatenated alignments are provided in the linked supporting database https://osf.io/3pmxb/ (Dorrell et al., 2021b) in the folder “Multigene tree topologies”.

### 18S and 16S phylogeny

18S rDNA and chloroplast 16S rDNA sequences corresponding to cultured isolates, and uncultured environmental isolates from the Arctic and Antarctic/Southern Oceans were downloaded for four algal groups (cryptomonads; haptophytes; chrysophyte-related taxa, including chrysophytes, synchromophytes, pinguiophytes and eustigmatophytes; and pelagophytes/ dictyochophytes) (Dorrell et al., 2021a), from GenBank nr (March 2020) (*59*). These were supplemented with orthologues searched for each group from MMETSP nucleotide sequence libraries by BLASTn, using a randomly selected downloaded query from the group considered, and a threshold e-value of 10^-05^. Homologous sequences were identified by a reciprocal BLASTn search against a complete copy of the *Arabidopsis thaliana* chloroplast, mitochondria and nuclear genome enriched with the query sequence; only sequences that retrieved the query as the BLAST best hit were retained (Sato, Nakamura, Kaneko, Asamizu, & Tabata, 1999). A complete set of query sequences are provided in Table S1, sheet 4.

Retained homologues were aligned using MAFFT (v. 7.407) under the --auto and --adjustdirection settings; and manually edited to remove non-homologous, fragmented and chimeric sequences (Katoh et al., 2017). Curated alignments were trimmed with trimAl under the -gt 0.5 and -gt 0.8 settings (Capella-Gutiérrez et al., 2009); and tree topologies were inferred from trimmed alignments using MrBayes v 3.2.1 and RAxML v 8.0 as integrated into the CIPRES web server (Miller et al., 2015; Stamatakis, 2014). MrBayes trees were run with two chains for 600,000 generations with burnin fractions of 0.5, and were manually verified in each case to have reached a final convergence statistic of ≤0.1 prior to calculation of the consensus topology; whereas RAxML trees were run by default for 450 bootstrap replicates, with automatic bootstopping applied. Curated alignments, individual and consensus tree topologies are provided in Table S1, sheets 4-7; and in the linked supporting database https://osf.io/3pmxb/ (Dorrell et al., 2021b) in the folder “18S and 16S trees”.

### Tara Oceans calculations

Ribotypes corresponding to the 18S rRNA (V4 and V9 variable regions) and 16S rRNA (V4V5 regions) sequences of each species were identified from version 2 (Arctic-inclusive) of *Tara* Oceans data (Ibarbalz et al., 2019) using the 18S and 16S trees defined above as a guide. Briefly, the complete 18S or 16S sequence of the species in question was searched by BLASTn against all 18S or 16S plastidial sequences previously assigned to the same taxonomic group as the query (*Baffinella* sp. CCMP2293: cryptomonads; Pavlovales sp. CCMP2436: haptophytes; *Ochromonas* sp. CCMP2298: chrysophytes; novel pelagophyte CCMP2097: pelagophytes), with threshold e-value 10^-05^. Matching sequences were then searched against the complete curated 18S or 16S alignments for each group using BLASTn, and only sequences that yielded BLAST best hits against either the query sequence or its immediate sister groups in the 18S or 16S tree topology were retained for subsequent analysis.

Retained ribotypes were realigned against the reference library using MAFFT under the --auto setting, manually trimmed to retain only the 18S V4, 18S V9, or 16S V4V5 regions, and finally trimmed with trimAl under the –gt 0.5 setting (Capella-Gutiérrez et al., 2009; Katoh et al., 2017); and trees were calculated from curated alignments using RAxML with automated bootstopping, as defined above(Stamatakis, 2014). Finally, ribotypes that were inferred from the best-scoring RAxML tree topology to be more closely related to the Arctic query species than the nearest cultured non-Arctic reference; and had a minimum 97% nucleotide similarity to the Arctic query species 18S or 16S rDNA sequence as assessed by BLASTn search; were retained for the calculation of absolute and total relative abundances, expressed as the proportion of all 18S V4, 18S V9 or 16S V4V5 ribotypes present in a sample. Curated alignments and tree topologies are provided in Table S1, sheets 8-9; tabulated individual and total read abundances are provided in Table S1, sheets 10-11; and all raw data pertaining to the identification of matching ribotypes by BLAST and phylogeny, and calculation of quantitative abundance trends, are provided in the linked supporting database https://osf.io/3pmxb/ (Dorrell et al., 2021b) in the folder “TARA Oceans calculations”.

### Quantitative analysis of PFAM content

PFAM distributions for each algal genome in the pan-algal dataset (both newly and previously sequenced accessions) were reannotated for this study using InterProScan and an updated (December 2020) version of the PFAM database from the constituent fasta files for each genome (Jones et al., 2014; Mistry et al., 2020). PFAM annotation files for each MMETSP transcriptome were manually downloaded from the source accession; and cleaned using a previously defined pipeline which compares the relative BLAST similarity between pairs of nucleotide sequences in each MMETSP library to identify transcripts of potential contaminant origin (Dorrell et al., 2019; Marron et al., 2016). Tabulated PFAM outputs are provided in Table S2, sheet 6; and complete PFAM lists per gene for each decontaminated library are provided in the linked supporting database https://osf.io/3pmxb/ (Dorrell et al., 2021b) in the folder “PFAM Bray-Curtis distributions”.

Phylogenetically-aware PCA was performed on both genome and transcriptome datasets (Revell & Graham Reynolds, 2012), due to strong taxonomic effects observed using crude phylogeny-independent analyses. Edited versions of the multigene tree topology (Fig. 1) retaining only genome- or transcriptome branches, were generated using MEGA version X as phylogenetic templates (Kumar, Stecher, Li, Knyaz, & Tamura, 2018). Input and output data for PCA are provided for user exploration in Table S2, sheet 7; and in the linked supporting database https://osf.io/3pmxb/ (Dorrell et al., 2021b) in the folder “CAFE and phylPCA”.

Similarity in PFAM content between different algal PFAM libraries were calculated using Bray-Curtis (Anderson et al., 2016; Horn et al., 2016) and Spearman indices (Choi & Kim, 2007; Nelson et al., 2021) following previous studies; and the total number concordant PFAMs between each library pair were additionally normalised against the total number of complete (single-copy or duplicated) eukaryotic BUSCOs retrieved in each library (Simão, Waterhouse, Ioannidis, Kriventseva, & Zdobnov, 2015). A schematic diagram explaining the methodology used, and the different types of convergence visible using each technique, is provided in Fig. 3- Figure Supplement 2.

For Bray-Curtis and Spearman calculations, algae were divided into three functional categories: Arctic (species isolated from > 60°N); Antarctic (species isolated from > 55°S); and all remaining taxa from intermediate latitudes grouped as “Other”. To avoid introducing biases due to comparing taxa of unequal phylogenetic distance, Bray-Curtis and Spearman calculations were only compared between pairs of algal libraries from different taxonomic groups, and to avoid biases due to comparing taxa with highly divergent life-strategies, all freshwater and secondarily non-photosynthetic taxa were removed from the final dataset. These values were then used to calculate mean values, and perform one-way ANOVAs of difference, within and between different groups of algae in the dataset, separated by biogeography into “Arctic”, “Antarctic” and “Other” taxa, as above. Tabulated Bray-Curtis and Spearman outputs are provided in Table S2, sheets 8-12; and all raw data, including alternative format outputs, are provided in the linked supporting database https://osf.io/3pmxb/ (Dorrell et al., 2021b) in the folder “PFAM Bray-Curtis distributions”.

### Identification of Arctic-associated PFAMs

PFAMs whose presence or absence were specifically associated with Arctic species were assessed by calculating the frequency with which the PFAM was recovered in Arctic, compared to non-Arctic species (“Antarctic” or “Other”) in the dataset (Table S2, sheet 6). These frequencies were used to calculate a ratio and a chi-squared P-value of enrichment of the PFAM in Arctic species in the dataset with cutoff P-value 10^-05^. Only PFAMs that were detected in both Arctic genomes and Arctic transcriptomes were considered as possible candidates for enrichment; and “Freshwater” and “Non-Photosynthetic” species were excluded from the dataset to avoid introducing artifacts by comparing species with highly divergent life strategies.

To accurately identify expansions and contractions in PFAM content across each species, high-frequency PFAMs (defined as PFAMs for which the maximum frequency minus minimum frequency observed was greater than 100 across the entire dataset) were first fragmented into smaller orthogroups. Briefly, profiles were extracted for each high frequency PFAM using hmmfetch (Potter et al., 2018), and then re-annotated for all libraries using hmmsearch with threshold expect value 10^-05^. Proteins containing these domains were used to run OrthoFinder v 2.4.1 (Emms & Kelly, 2019) with inflation value 1.3 to avoid over-fragmentation of orthogroups. Orthogroups were filtered (maximum – minimum frequency > 2) to remove uninformative examples, then the presence of PFAM domains (defined as presence in at least 10% of protein space within the orthogroup) were called for each orthogroup.

Computational Analysis of gene Family Evolution (CAFE) was performed on the composite set of low-frequency PFAMs, and orthogroups decomposed from high-frequency PFAMs, for separate sets of genome- and transcriptome-only libraries, using MEGA-edited versions of the previously generated multigene reference tree, as per the phylPCA analysis above. (Kumar et al., 2018) A single gamma rate for the lambda and error model was assessed from the low-frequency PFAM dataset across the entire phylogenetic tree. The CAFE outputs obtained were used to calculate enrichment ratios and P-values for “expansions”, defined as signed positive CAFE scores, and “contractions”, defined as signed negative CAFE scores for Arctic species within the dataset, using methodology as defined above. Summarised CAFE outputs and P-values are provided in Table S3, sheets 1-2; and all raw data, including the prior decomposition of PFAMs into orthogroups, are provided in the linked supporting database https://osf.io/3pmxb/ (Dorrell et al., 2021b) in the folder “CAFE and phylPCA”.

### Environmentally supported phylogenies of Arctic-associated PFAMs

The environmental distributions of sequences containing three PFAMs (PF03831, PF03988, and PF11999) whose presence was significantly associated with Arctic species in the enrichment analysis above were investigated in meta-transcriptome and meta-genome sequence data from the *Tara* Oceans Expedition, inclusive of sequences from the Arctic Ocean (Carradec et al., 2018; Ibarbalz et al., 2019). Sequences containing each PFAM were extracted using hmmer with gathering threshold option (Potter et al., 2018); and were classified into four categories based on the sum of all relative abundances (normalised against the total number of meta-T or meta-G unigenes sequenced from the sample) across all stations and size/depth fractions. These were: “Arctic” sequences (> 70% summed relative abundances contained in stations of > 60 °N); “Antarctic” sequences (> 70% summed relative abundances contained in stations of > 55°S); “Bipolar” sequences, (> 70% summed relative abundances contained in stations of > 60°N or > 55°S, including >20% each in stations > 60°N and stations > 55°S); and “Other” (< 70% summed relative abundances contained in stations of > 60°N and > 55°S).

The *Tara* Oceans sequences were aligned against all sequences containing the PFAM concerned from all algal genomes and transcriptomes considered within this study; and all sequences containing the PFAM in UniRef (downloaded March 2020) (Suzek et al., 2007); using mafft under the --auto setting, followed by a more stringent round of alignment using --gap_open_penalty 12, -- gap_extension_penalty 3 and --maxiteration 2 settings to remove poorly aligned sequences (Katoh et al., 2017). Cultured sequence accessions were manually labelled with “Arctic” and “Antarctic” provenance considering the isolation site of the species, where recorded (Guiry et al., 2014).

Alignments were manually trimmed after each step, and subsequently trimmed with TrimAl using the –gt 0.5 setting (Capella-Gutiérrez et al., 2009); and finally used to infer best-scoring trees with RAxML using the PROTGAMMAGTR, PROTGAMMAJTT and PROTGAMMAWAG substitution matrices (Stamatakis, 2014). Environmental sequence calculations, and individual and consensus topologies for each tree are provided in Table S3, sheets 3-7; and in the linked supporting database https://osf.io/3pmxb/ (Dorrell et al., 2021b) in the folder “Environmental PFAM calculations”.

### Global identification of within-Arctic HGTs

Global biases in algal genome or transcriptome composition, which may reveal underlying HGT events, were inferred using a linear regression pipeline adapted from previous studies (Dorrell et al., 2021a; Stiller et al., 2014)(Fig. 6- Figure Supplement 1). To avoid incorporating biological contaminants into these analyses, all four Arctic genomes were initially refined to a “best confidence” set of genes, as described above, consisting of genes present on an assembled contig with at least one gene of clear vertical origin (*Baffinella* sp. CCMP2293: cryptomonads; Pavlovales sp. CCMP2436: haptophytes; *Ochromonas* sp. CCMP2298: chrysophytes and novel pelagophyte CCMP2097: pelagophytes) based on BLASTp best-hit analysis of a combined library from across the tree of life as defined below (Dorrell et al., 2021a); and presence of a corresponding transcript in decontaminated MMETSP transcriptome assemblies for the species, inferred by reciprocal BLASTp best-hit searches (Keeling et al., 2014).

Next, a *comparative pair* of query genomes from a specific algal lineage, one Arctic and one non-Arctic (e.g., *Baffinella* sp. CCMP2293, and *Guillardia theta*, from cryptomonads (Curtis et al., 2012)) were searched against a composite dataset of Arctic and non-Arctic genomes and transcriptomes from another algal group (e.g., chlorophytes) via LAST (Kiełbasa et al., 2011), and the best-scoring reference sequences (defined by bitscore) to each query gene were recorded. As both *Baffinella* CCMP2293 and *G. theta* are of equivalent phylogenetic distance to all chlorophytes, the number of LAST best hits obtained by the *Guillardia* genome to each reference chlorophyte sequence should act as a predictor of the number of hits obtained by the *Baffinella* genome, with positive deviations from this, as inferred by linear regression, potentially relating to more recent HGTs between the *Baffinella* genome and the reference species. A schematic diagram explaining the methodology; and exemplar scatterplot outputs, are provided in Fig. 6- Figure Supplement 1. These analyses were repeated independently for four comparative pairs of Arctic and non-Arctic query genomes (*Baffinella* CCMP2293 v *Guillardia theta;* Pavlovales CCMP2436 v *Chrysochromulina tobin*; *Ochromonas* CCMP2298 v *Nannochloropsis gaditana*; and the novel pelagophyte CCMP2097 v *Aureococcus anophageferrens*), and combined reference libraries generated from the eight algal lineages considered in this study (chlorophytes, chrysophytes, cryptomonads, diatoms, dictyochophytes, dinoflagellates, haptophytes and pelagophytes), with the exception of within-taxon comparisons (e.g., cryptomonad query genomes were not compared to the cryptomonad reference dataset).

Details of the scripts, and tabulated LAST best hit frequencies, are provided in Table S4, sheets 2-3; and in the linked supporting database https://osf.io/3pmxb/ (Dorrell et al., 2021b) in the folder “Within-Arctic HGTs”. Altogether, 1,322 genes from Arctic genomes that either retrieved an Arctic LAST best hit when searched against at least one of the algal reference libraries; or that themselves were retrieved as LAST best hits; or corresponded to MMETSP transcripts retrieved as LAST best hits to another Arctic genome query were used as seed sequences for subsequent identifications of within-Arctic HGT (schematic workflow presented in Fig. 6- Figure Supplement 2). Briefly, these seed sequences were searched by BLASTp against a previously assembled composite reference library (Dorrell et al., 2019; Dorrell et al., 2021a) containing a complete copy of UniRef (Suzek et al., 2007); all JGI algal genomes (Grigoriev et al., 2021); and decontaminated copies of the MMETSP (Keeling et al., 2014; Marron et al., 2016) and OneKP transcriptomes (Carpenter et al., 2019; Initiative, 2019); along with independently sequenced reference transcriptomes e.g., from diatoms (Dorrell et al., 2021a) and chrysophytes (Beisser et al., 2017); and here manually annotated by isolation site (Arctic, Antarctic or Other as defined above). 229 genes that (i) did not yield a BLAST best hit against the same taxonomic group as the query (*Baffinella* sp. CCMP2293: cryptomonads; Pavlovales sp. CCMP2436: haptophytes; *Ochromonas* sp. CCMP2298: chrysophytes and novel pelagophyte CCMP2097: pelagophytes) and (ii) included at least one Arctic species in the best ten BLASTp hits were retained for subsequent analysis. Query genes searched are provided in Table S4, sheet 4; and the ten best BLASTp hits for each query sequence in Table S4, sheet 6.

A full set of orthologues were assembled for each retained query sequence by retrieving the LAST best hit for 151 non-redundant taxonomic groups from the all tree of life library used above (Dorrell et al., 2019; Dorrell et al., 2021a); the pan-algal dataset of genomes and transcriptomes used in this study; and a further set of 33 eukaryotic reference genomes and 59 prokaryotic reference genomes, sampled to include diverse representation from the tree of life, including multiple Arctic and Antarctic prokaryotes (Table S4, sheet 5); along with all sequences identified from the genomes and MMETSP transcriptomes of each of the four Arctic algae sequenced in this study that could be retrieved by LAST with threshold e-value 10^-05^. Clusters for pairs of seed sequences that were found to show greater similarity to one another than any non-seed sequence within the cluster via internal BLAST searched were merged, yielding 215 merged gene clusters of non-redundant seed sequences and a total set of approx. 300-500 orthologues from across the tree of life (Fig. 6- Figure Supplement 2).

Alignments were constructed for each merged gene cluster using MAFFT with the --auto setting; followed by a second round of alignment with the --gap_open_penalty 12 --gap_extension_penalty 3 --max_iteration 2 settings; then trimmed using TrimAl with the –gt 0.5, -resoverlap 0.75 and - seqoverlap 0.8 settings; followed by a final round of alignment with MAFFT (Capella-Gutiérrez et al., 2009; Katoh et al., 2017). Clusters containing poorly aligned or fragmented seed sequences were manually identified and rejected at each step, retaining 129 approved gene clusters (Fig. 6- Figure Supplement 2). Approved gene clusters were used to generate RAxML consensus trees, with 300 bootstrap replicates and the PROTGAMMAJTT substitution model. 21 of the realised trees were found to contain monophyletic groups of multiple Arctic species with > 50% bootstrap support; and a further 13 contained groups consisting of a majority of Arctic species with some Antarctic or non-polar species, with > 50% support for the ancestral node (Fig. 6- Figure Supplement 2; Table S4, Sheet 9). Finalised alignments and consensus tree topologies are provided in Table S4, sheets 7-8; and all intermediate alignment processing steps, and individual bootstrap replicates for each tree, are provided in the linked supporting database https://osf.io/3pmxb/ (Dorrell et al., 2021b) in the folder “Candidate HGT trees”.

Inferred functions were inspected for all genes from each Arctic query genome identified at each stage of the pipeline (all decontaminated genes; all genes with a within-Arctic LAST best hit; all genes with an Arctic top ten BLAST best-hit; and all genes confirmed by single-gene trees to be participants of within-Arctic HGTs). The assessed functions included KEGG and KOG annotations, assessed by BlastKOALA and GhostKOALA (Kanehisa & Sato, 2020); PFAM domains, inferred by InterProScan (Jones et al., 2014); absolute and ranked rpkm values of transcripts corresponding to each gene in corresponding MMETSP transcriptomes derived under different physiological conditions (Terrado et al., 2015); and predicted subcellular localisations, inferred using ASAFind v 2.0 used in conjunction with Signal P v 3.0 (Gruber, Rocap, Kroth, Armbrust, & Mock, 2015), HECTAR using a default scoring matrix (Gschloessl, Guermeur, & Cock, 2008), and WolfPSort, taking the consensus annotations obtained with animal, fungal and plant reference models (Horton et al., 2007). Tabulated functions associated with each sequence, and two-tailed chi-squared enrichment values for each function in within-Arctic HGTs, are provided in Table S4, sheets 9-12. Finally, *Tara* Oceans unigenes corresponding to within-Arctic HGTs were identified by hmmer using HMMs composed of all Arctic isolate sequences from the untrimmed alignments for each gene family, as above. Peptide sequences of the unigenes retrieved using this approach were first searched against all sequences in the corresponding gene family alignment by BLASTp with the - max_target_seqs 1 parameter. Unigenes that yielded a best BLAST hit against an Arctic species were realigned against the gene family alignment with mafft under the --auto setting; and used to build neighbour-joining trees with 100 replicates in GeneIOUS v. 10.0.9. Unigenes that mapped to within-Arctic HGT clade within these trees were extracted, and used to calculate total relative environmental abundances across all depths and size fractions in *Tara* Oceans data, as above. Details of the BLAST outputs, alignments, phylogenetically reconciled TARA sequences and relative abundance calculations, are provided in Table S4, sheets 10-11; and in the linked supporting database https://osf.io/3pmxb/ (Dorrell et al., 2021b) in the folder “Within-Arctic HGTs”.

### Data deposition

The genome assemblies and annotations are available from JGI PhycoCosm portal (Grigoriev et al., 2021)and have been deposited in DDBJ/ENA/GenBank with the following URLs and NCBI accessions: CCMP2293:https://phycocosm.jgi.doe.gov/Crypto2293_1, PRJNA223438; CCMP2436: https://phycocosm.jgi.doe.gov/Pavlov2436_1 PRJNA223446; CCMP2298: https://phycocosm.jgi.doe.gov/Ochro2298_1, PRJNA171379; CCMP2097:https://phycocosm.jgi.doe.gov/Ochro2298_1, PRJNA210205;.

All additional supporting data, including the composition of the global and taxonomic reference datasets used in this study; comparisons to previously published genome assemblies (Armbrust et al., 2004; Derelle et al., 2006; Gobler et al., 2011; Grigoriev et al., 2021; Hovde et al., 2015; Lin et al., 2015; Lommer et al., 2012; Mock et al., 2017; Radakovits et al., 2013; Rastogi et al., 2018; Read et al., 2013; D. M. Wang et al., 2014; Worden et al., 2009); 18S, 16S and multigene trees; *Tara* Oceans Expedition calculations; phylPCA, CAFE and PFAM frequency analyses; LAST and phylogenetic analyses of within-Arctic HGTs and Arctic-specific gene expansions and contractions are provided via a dedicated osf.io database (Dorrell et al., 2021b) https://osf.io/3pmxb/. Data is classified within this database by project, each of which contains an associated README file describing the folder contents, and associated methodology for its production.

## Author contributions

CL isolated the strains and coordinated the global study. MP and REE grew and maintained the culture, with MPB responsible for nucleic acid isolation and quality control. KB, JS, and IVG coordinated genome projects, and RGD coordinated functional analysis of each species. AK, JJ, LH, AS, CP, AL, YP and KL produced the genome data; and RGD, AK, ASS, NZ, ZF, FI, JJPK and ER analysed the genome data. AMGNV, JM, NJF, FRJV, VL, BH, JBD, CdV and PW provided additional supporting data. RGD, AK, ER, ZF, CL, CB and IVG wrote the manuscript with comments on different sections by co-authors.

## Acknowledgments

The work conducted by the U.S. Department of Energy Joint Genome Institute, a DOE Office of Science User Facility, is supported by the Office of Science of the U.S. Department of Energy under Contract No. DE-AC02-05CH11231. RGD and AMGNV acknowledge funding from a CNRS Momentum Fellowship (2019-2021) awarded to RGD. CL received support from the Discovery program of Natural Science and Engineering Council (NSERC, Canada) and a pilot project grant from Genome Québec. CB acknowledges funding from the European Research Council (ERC) under the European Union’s Horizon 2020 research and innovation programme (grant agreement No. 835067; Diatomic), the French Government ‘Investissements d’Avenir’ programmes MEMO LIFE (ANR-10-LABX-54), PSL* Research University (ANR-1253 11-IDEX-0001-02), France Génomique (ANR--10-INBS-09) and OCEANOMICS (ANR-11-BTBR-0008), and Research Grant ‘Green Life in the Dark’ (RGP0003/2016) from the Human Frontier Science Program. CB, FMI and JJPK acknowledge support from ECOS Sud-Argentine program (AT08ST18). ZF acknowledges support from the J.W. Fulbright Commission of the Slovak Republic and computational resources supplied by the project “e-Infrastruktura CZ” (e-INFRA LM2018140) provided within the program Projects of Large Research, Development and Innovations Infrastructures. The authors thank E. Virginia Armbrust (University of Washington) and Jackie Collier (Stony Brook Univ. New York) for permission to use a *Pseudo-nitzschia multiseries* genome for comparative PFAM analysis, and *Aplanochrytrium kerguelense, Auriantiochytrium limacinum,* and *Schizochytrium aggregatum* transcriptomes for phylogenetic dataset construction, respectively. The authors respectfully acknowledge that the research within this project benefits from the sampled indigenous biodiversity of the Inuit Nunangat, and was partially performed on the traditional lands and Treaty 6 and unceded territories of multiple First Canadian Nations. This article is contribution number XX of the *Tara* Oceans project.

---

**Fig. 1 - Figure supplement 1.**
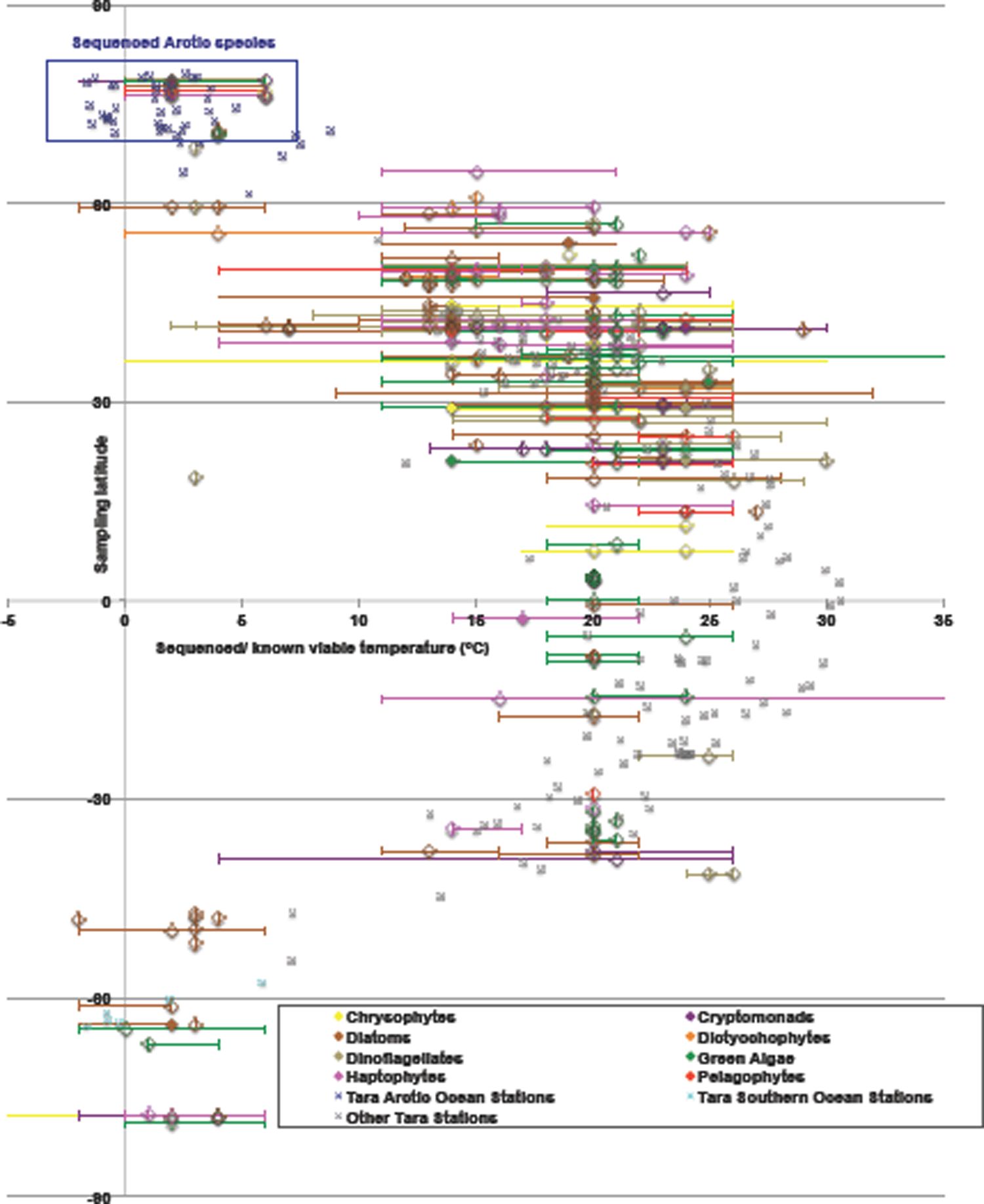
Scatterplot of sampled latitude, and growth temperatures, of all geolocalised algal genomes and transcriptomes within the pan-genomic dataset assembled for this study. Taxa are shaded by phylogeny and biogeography per Fig. 1. Data were manually verified for each culture by comparing the synthesis of the genome and transcriptome portal data (Grigoriev et al., 2021; Keeling et al., 2014) with recorded permissible growth temperatures for each corresponding culture collection accession, taking into account synonymous strain names housed in different collections. Tara Oceans data is taken from PANGAEA entries for each station (Pesant et al., 2015). Growth temperatures are provided as ranges, centred around the experimental temperature (if recorded) at which the genome or transcriptome library was generated. The Arctic species sequenced in this study, which are not viable at temperatures > 6C, are surrounded by a blue box.

**Fig. 1 - Figure supplement 2.**
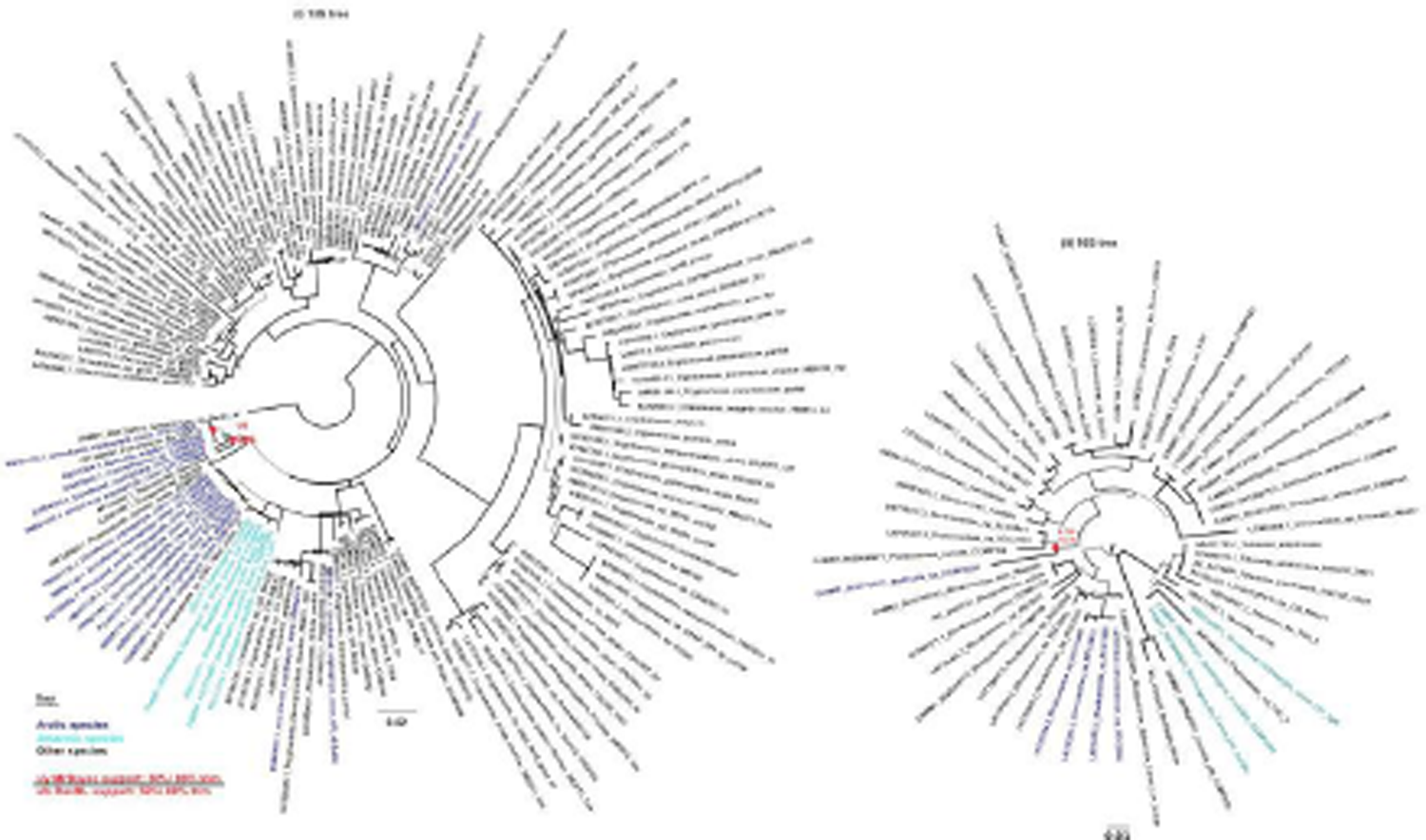
Phylogenetic position of Baffinella sp. CCMP2293 inferred from cultured 18S sequences. This figure shows the consensus topologies inferred for (left) a 143 taxa x 1699 nt 18S rDNA alignment and (right) a 53 taxa x 1178 nt alignment of all cryptomonad NCBI and MMETSP sequences, trimmed to 80% occupancy. The 18S tree is rooted on a *Goniomonas* outgroup (Cenci et al., 2018) and the 16S tree on the midpoint. Thick branches indicate clade presence in both MrBayes consensus trees obtained. Leaf nodes are shaded by biogeographical origin. MrBayes consensus and RAxML best-scoring tree values retrieved for nodes unifying *Baffinella* sp. CCMP2293 with a clade of mainly Arctic cryptomonads, including the conspecific CCMP2045 (Daugbjerg et al., 2018), and with the genus *Proteomonas*, are shown with red circles.

**Fig. 1 - Figure Supplement 3.**
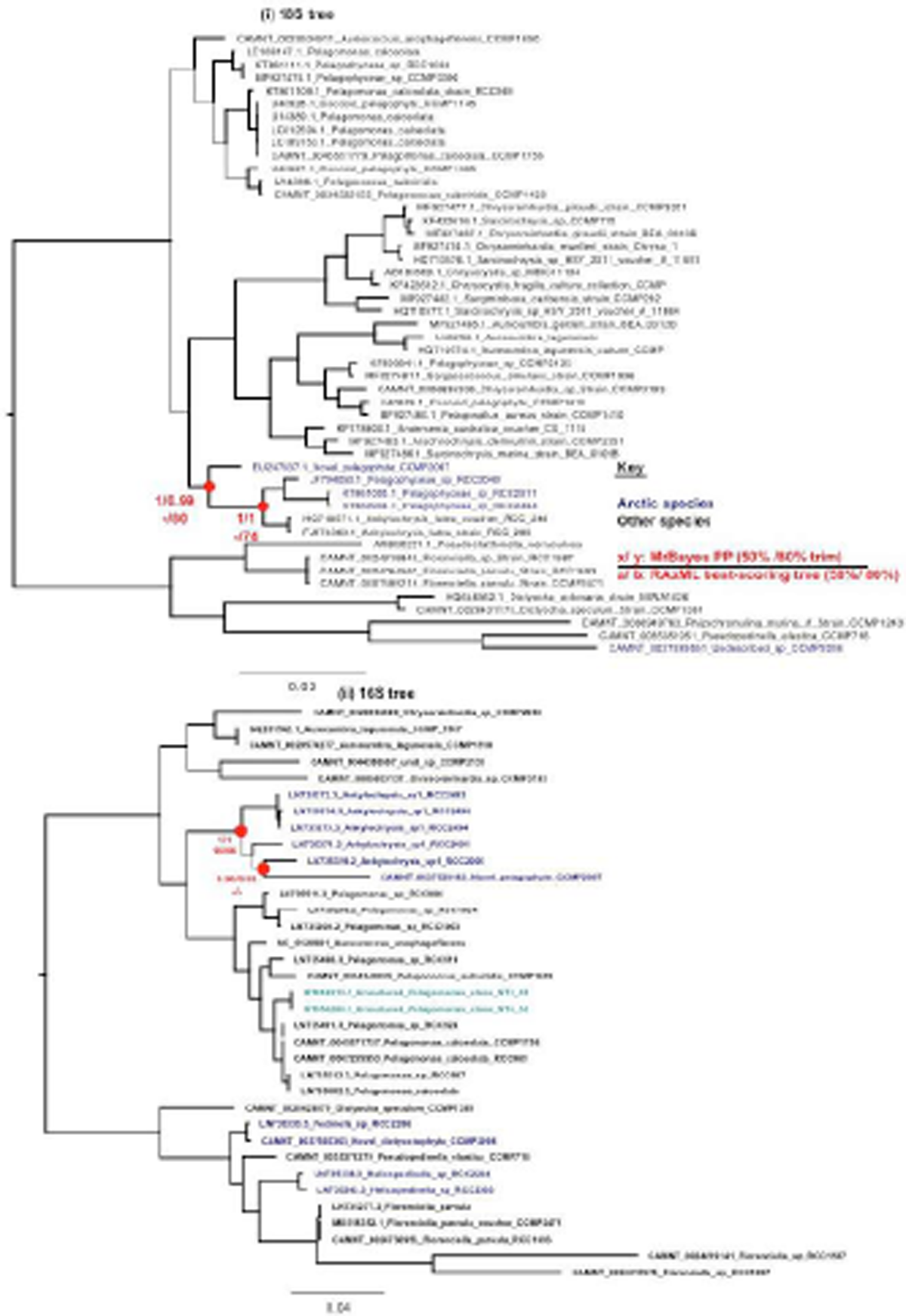
Phylogenetic position of the novel pelagophyte CCMP2097 inferred from cultured 18S and 16S sequences. Consensus MrBayes and RAxML topologies realised for (i) a 55 taxa x 1648 nt 18S alignment; and (ii) a 35 taxa x 819 nt 16S alignment of pelagophyte and dictyochophyte NCBI and MMETSP 18S sequences, rooted between pelagophytes and dictochophytes (Dorrell et al., 2021a), and trimmed to 80% occupancy. Thick bars indicate clade recovery in all tree topologies. Leaf nodes are shaded by biogeographical origin. Support values for nodes, placing CCMP2097 at within a predominantly Arctic clade including the genus *Ankylochrysis* (Han et al., 2018), are shown with red circles.

**Fig. 1 - Figure supplement 4.**
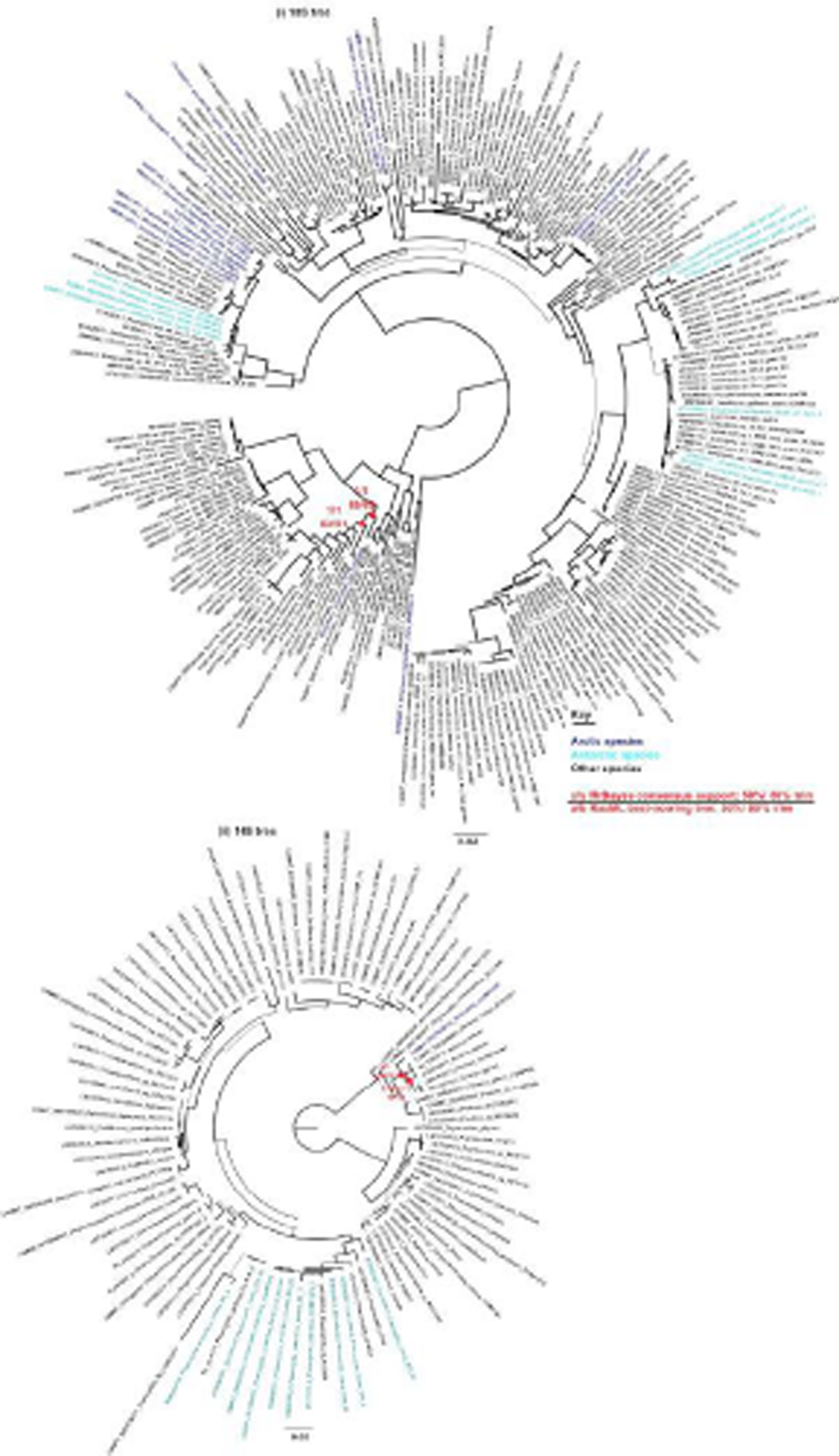
Phylogenetic position of Pavlovales sp. CCMP2436 inferred from cultured 18S and 16S sequences. Consensus MrBayes and RAxML topologies inferred for (i) a 241 taxa x 1679 nt 18S alignment and (ii) a 94 taxa x 846 nt 16S alignment of NCBI and MMETSP haptophyte trimmed to 80% occupancy and rooted on the split between pavlovophyte and prymnesiophyte sequences (Bendif et al., 2011). Thick branches indicate recovery of a clade in both tree topologies. Leaf nodes are shaded by biogeographical origin. Support values for two nodes realised using MrBayes and RAxML, which place CCMP2436 as an isolated basal member of the pavlovophytes,and related to the otherwise non-Arctic genus *Diacronema* sensu lato, are indicated with red circles.

**Fig. 1 - Figure Supplement 5.**
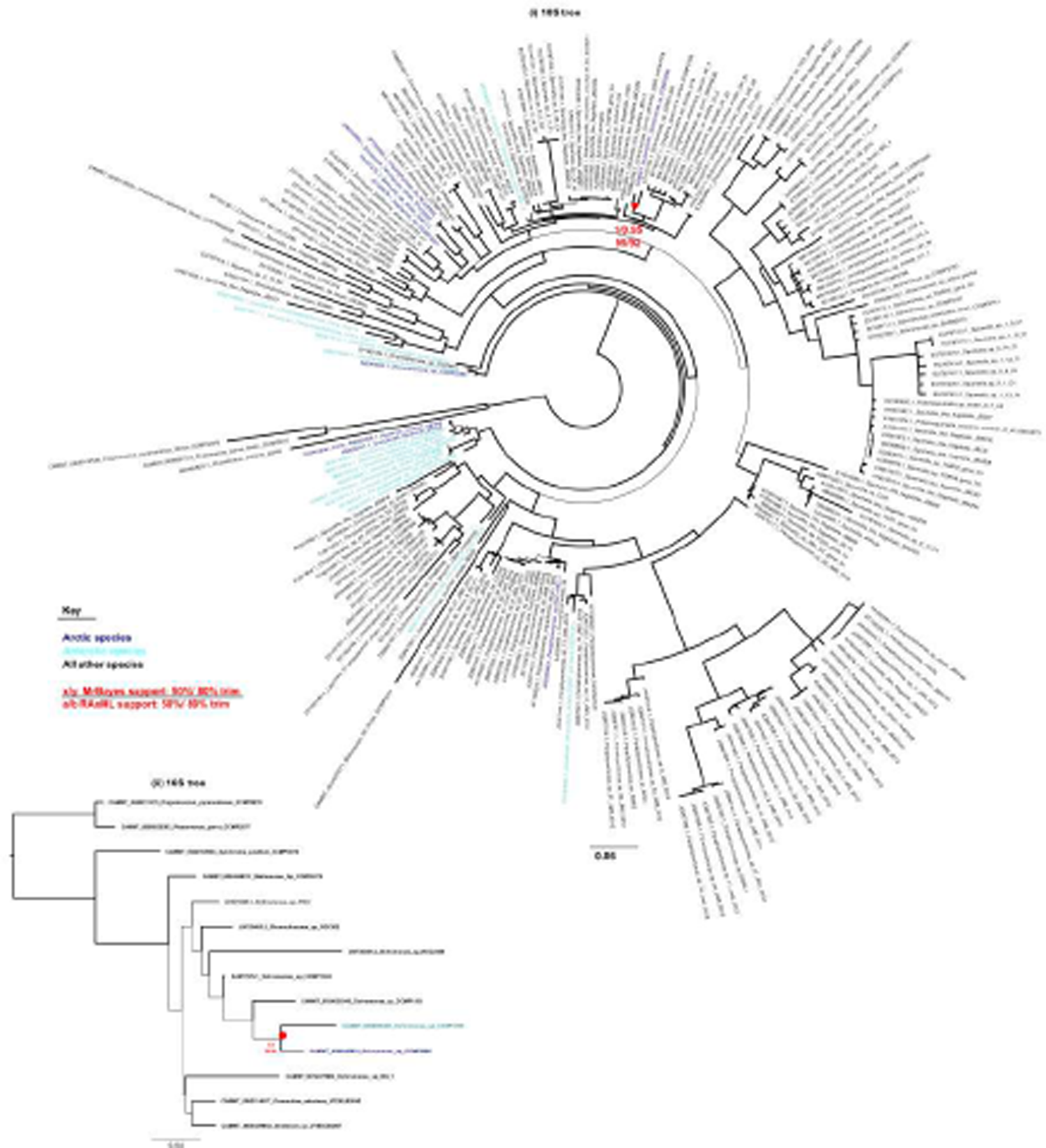
Phylogenetic position of *Ochromonas* CCMP2298 inferred from cultured 18S and 16S sequences. Consensus MrBayes and RAxML topology resolved for (top) a 219 taxa x 1636 nt alignment for NCBI and MMETSP 18S rDNA sequences from and (bottom) a 15 taxa x 1383 nt alignment (trimmed to remove sites with <80% occupancy) of chrysophytes, synurophytes, and a pinguiophyte outgroup (Dorrell et al., 2021a). Thick branches indicate presence of a node in the MrBayes consensus and RaXML best-scoring tree topology. Leaf nodes are shaded by biogeographical origin. Consensus values retrieved for one node linking CCMP2298 to the mesophilic *Ochromonas* sp. CCMP1393 (18S) and to *Ochromonas* sp. CCMP1899 (16S) in trees calculated with alignments trimmed to 50% and 80% occupancy are shown with red circles.

**Fig. 2 - Figure Supplement 1.**
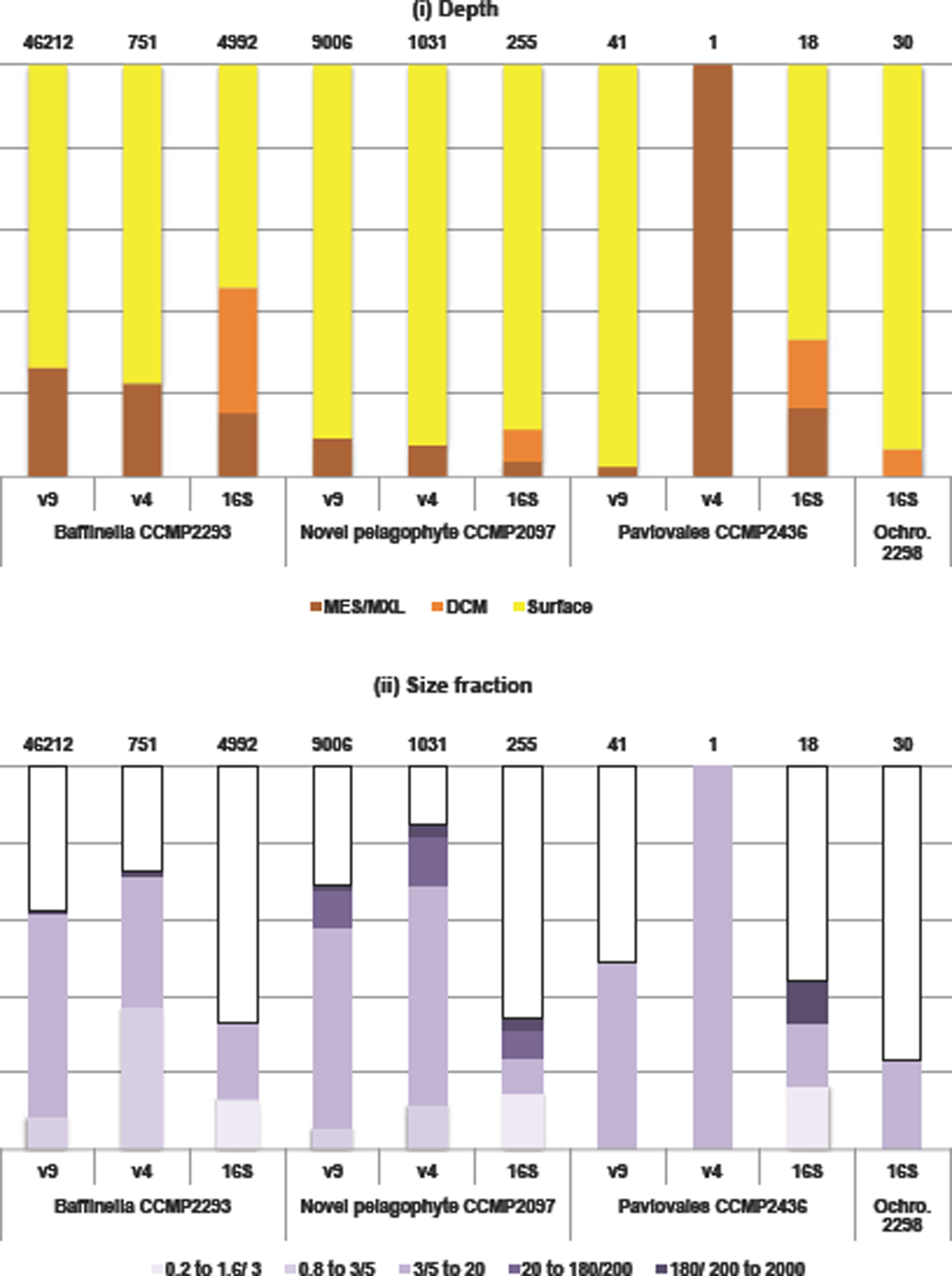
Total read counts of *Tara* Oceans ribotypes phylogenetically resolved to be closely related to Arctic native species: divided by **(i)** depth (either surface, deep chlorophyll maximum, or mesopelagic/ mixed layer), and **(ii)** size fraction. Each species is predominantly distributed in surface waters, small (3-5/ 20 μm) size fractions, and Arctic stations. The total numbers of reconciled ribotypes are shown on the above each plot. Samples with no mapped ribotypes (*Ochromonas* sp. CCMP2298 was not detected in 18S v4 or 18S v9 ribotype data hence these values are not shown).

**Fig. 2 - Figure Supplement 2.**
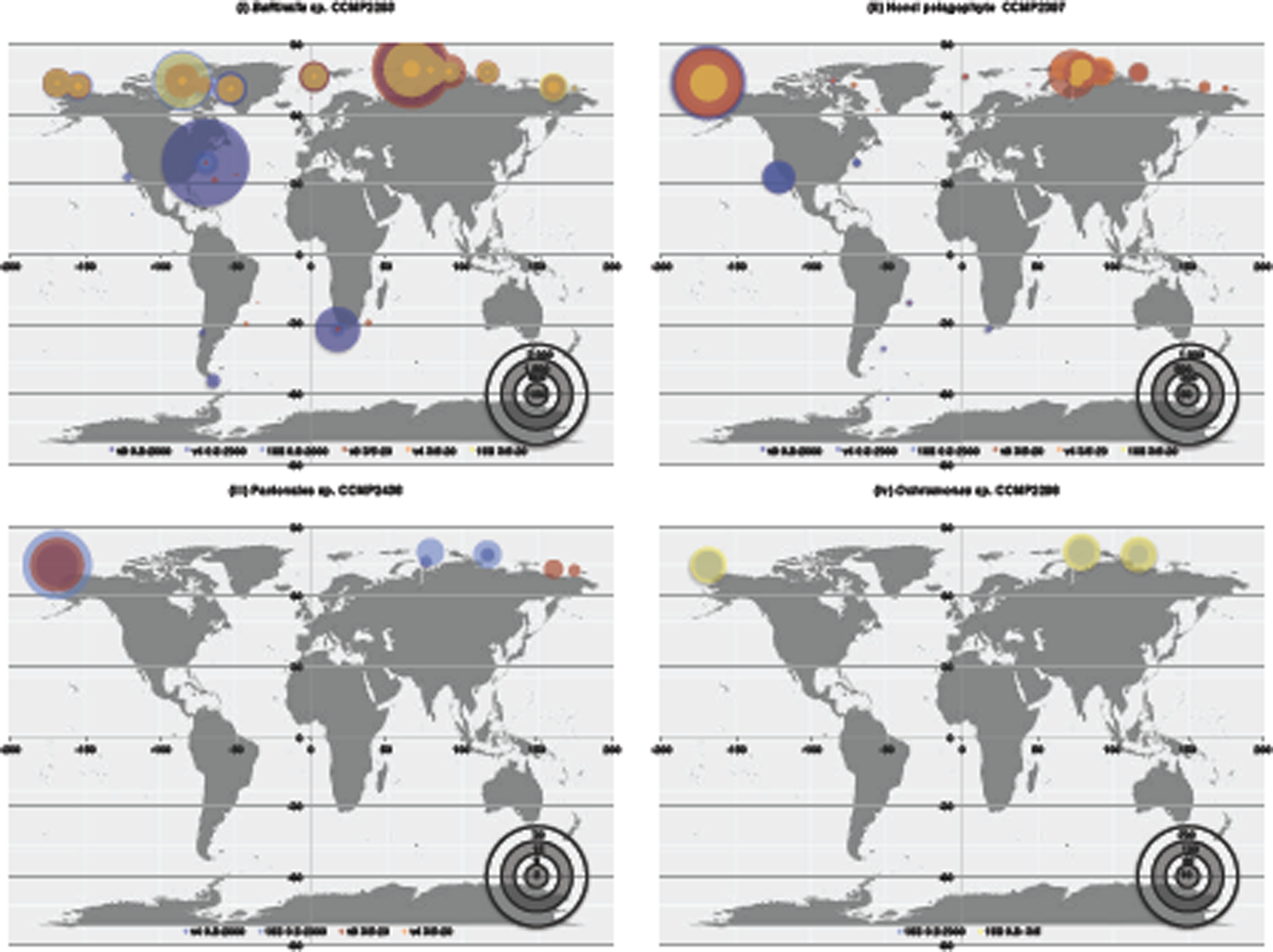
Relative abundances (expressed as parts per million) of 18S v4, 18S v9 and 16S v4v5 ribotypes of Arctic isolated algae: (A) *Baffinella sp*. CCMP2293**, (B)** novel pelagophyte CCMP2097, **(C)** Pavlovales sp. CCMP2436 and **(D**) *Ochromonas* sp. CCMP2298; calculated from the Tara Oceans and Tara Oceans Polar Circle expeditions. Ribotypes corresponding to each species were identified by BLASTn (threshold similarity 97%) followed by phylogenetic reconciliation to cultured accessions; and abundance calculations are shown for surface samples and size fractions (0.8-2000 and 3/5-20 μm) in which the greatest number of corresponding OTUs were counted.

**Fig. 2 - Figure Supplement 3.**
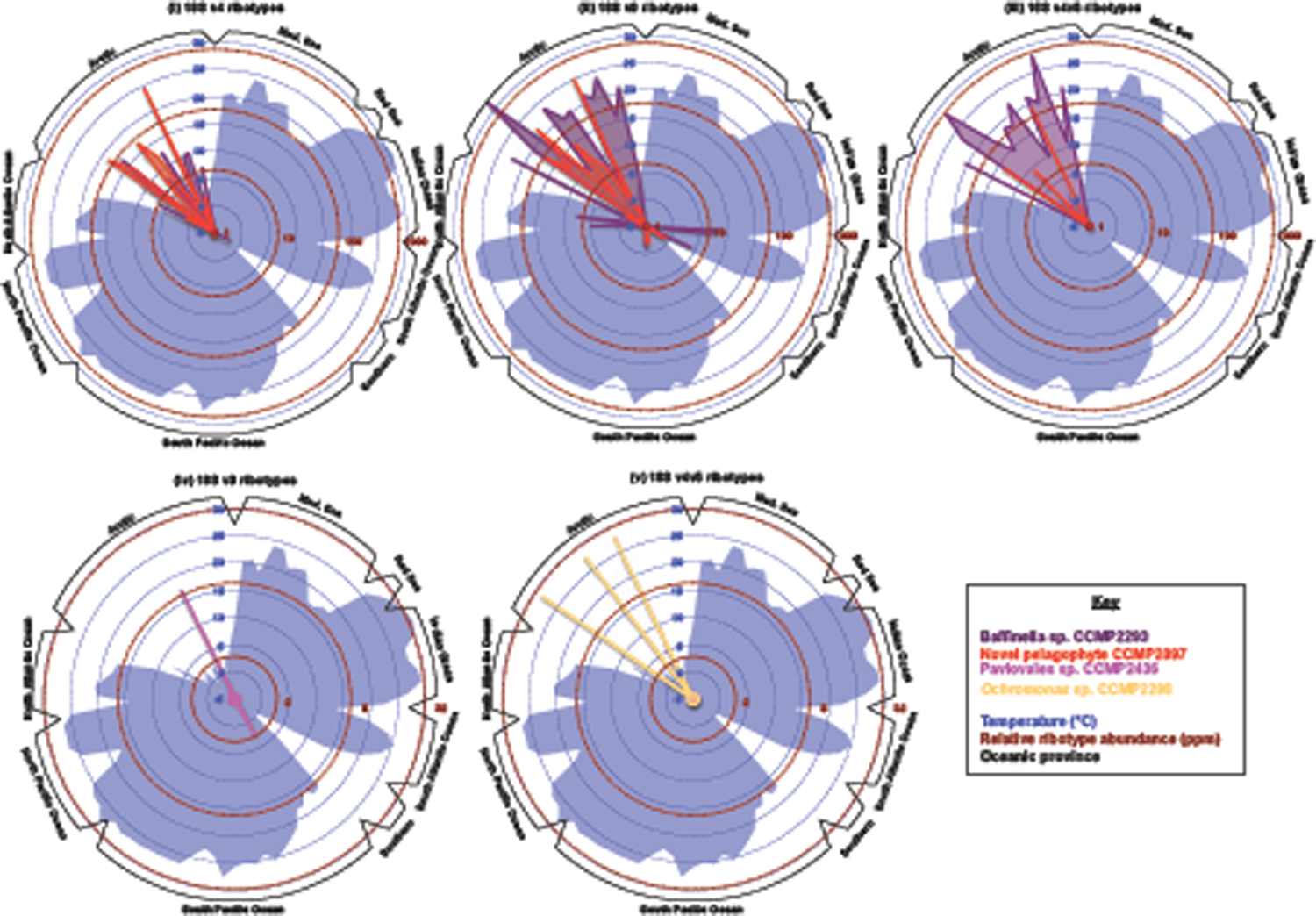
Cold water preferences of Arctic native algae. This figure shows radar plots of the relative abundances of (i-iii) *Baffinella* sp. CCMP2293 and the novel pelagophyte CCMP2097; (iv) Pavlovales sp. CCMP2436 and (v) *Ochromonas* sp. CCMP2298 in the 3/5-20 μm size fraction and surface samples of Tara Oceans station, divided by station provenance, and compared to environmental temperature. All four species show strong localisation preferences to cold stations in the Northern hemisphere (i.e. the Arctic Ocean) without an equivalent enrichment in cold Southern Ocean stations.

**Fig. 3 - Figure Supplement 1.**
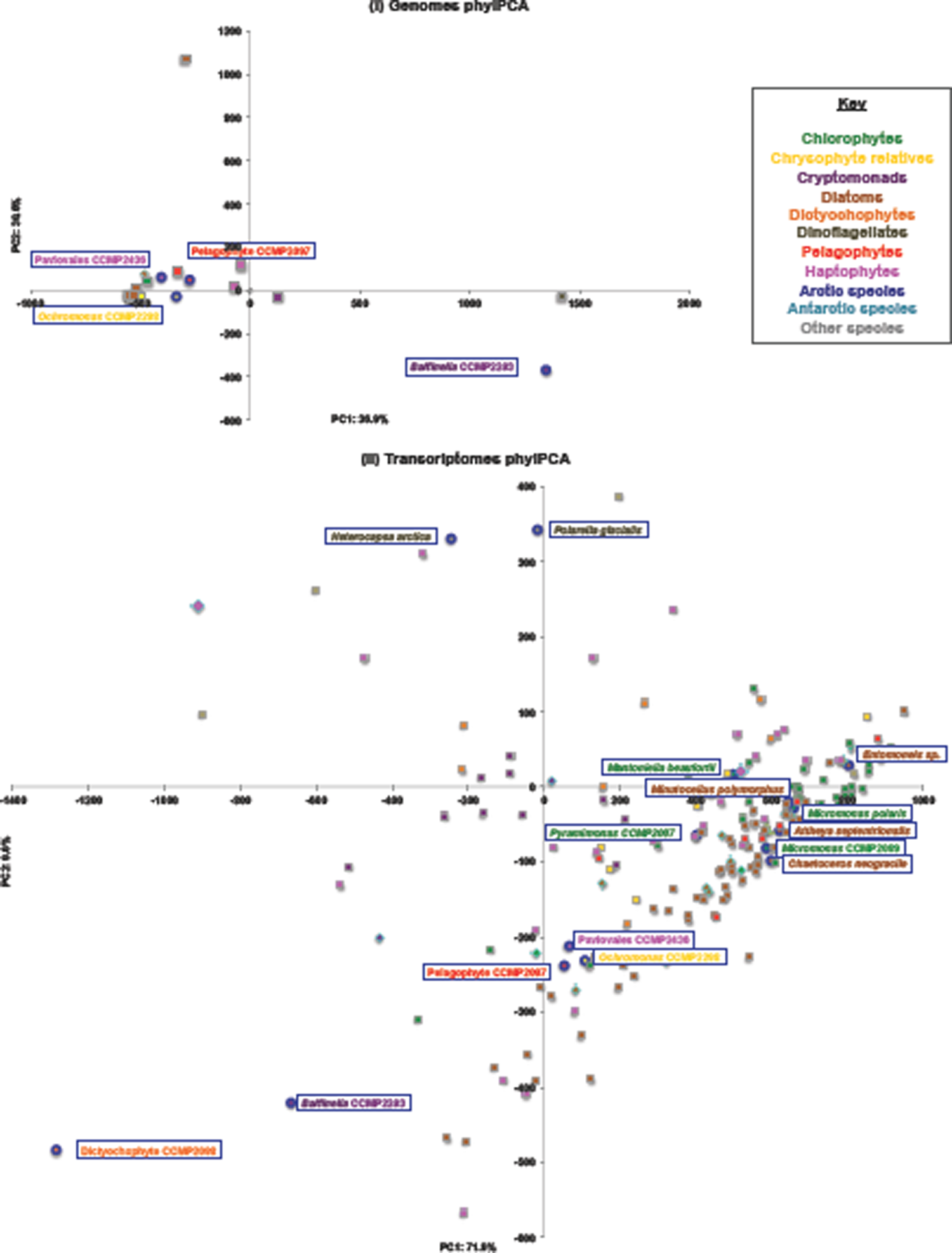
**Phylogenetically-aware principal component analyses of PFAM content in the pan-algal dataset**, separately realised (i) for genome and (ii) for transcriptome data. Full coordinate values are provided in Table S2, sheet 6. Each coordinate point is coloured by phylogeny (fill) and biogeographical origin (line). The all other Arctic species in the datasets are labelled.

**Fig. 3 - Figure Supplement 2.**
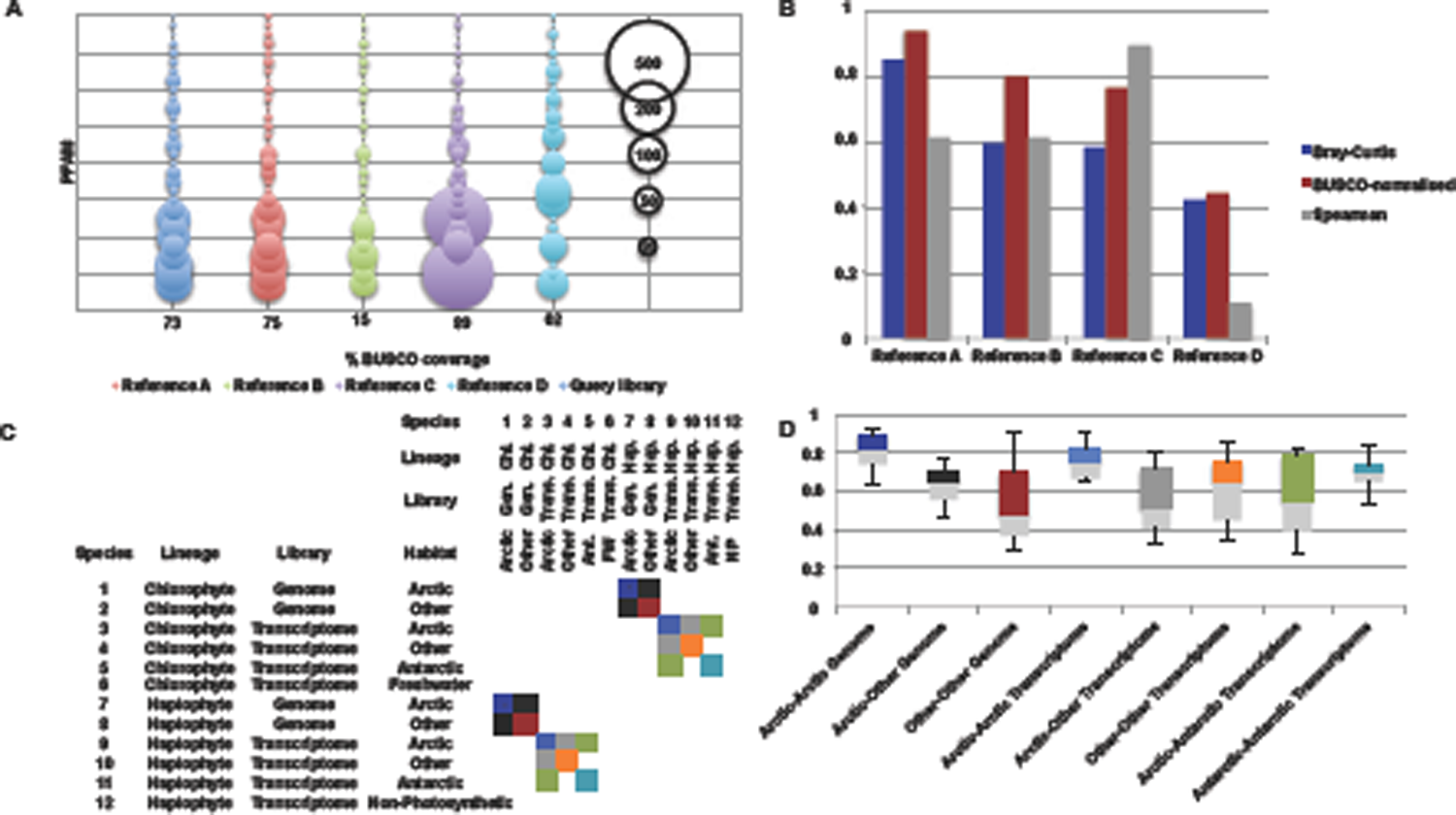
**Exemplar calculations of PFAM convergence. A**: bubble plot showing the distribution of 30 PFAMs in five hypothetical libraries. The query library is convergent to Reference A; Reference B is also convergent, but fragmented; Reference C is also convergent, but with specific expansions in individual PFAMs; and Reference D is non-convergent. **B**: calculated Bray-Curtis, BUSCO-normalised shared PFAM, and Spearman correlation values between the query library and References A-D. **C**: exemplar heatmap of pairwise comparisons between 12 libraries, and **D**: the convergence calculations made between them, shaded per the sampled intersecting cells in the heatmap. Of note, comparisons within individual lineages, between genomes and transcriptomes, and involving either freshwater or non-photosynthetic species were excluded to minimise latitude-independent biases on PFAM convergence estimates.

**Fig. 3 - Figure Supplement 3.**
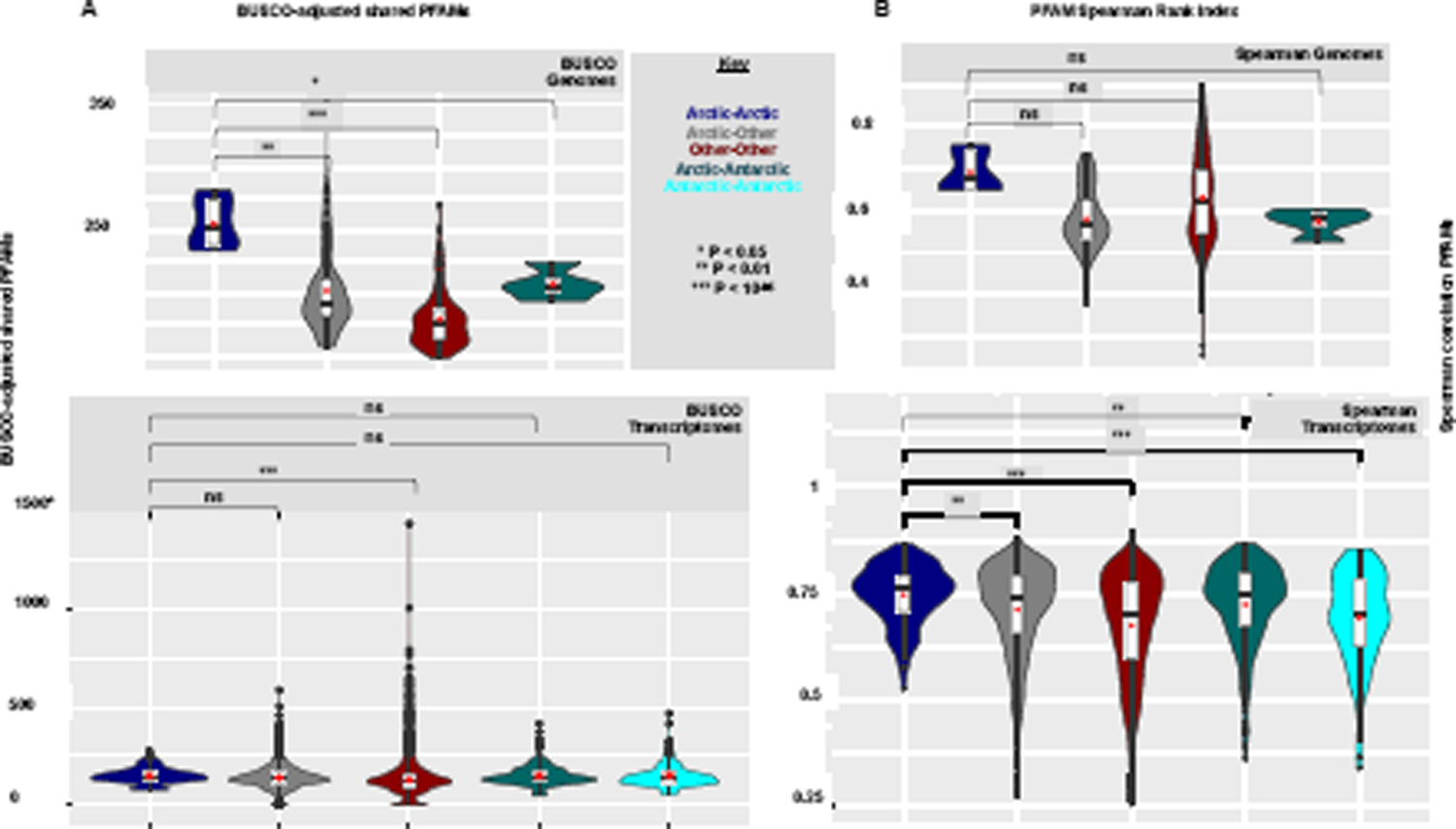
**Alternative metrics for identification of Arctic-Arctic PFAM convergences. A**: violin plot of pairwise numbers of total number of shared PFAMs normalised on % recovered complete (single or duplicated) eukaryotic BUSCOs, and **B**: violin plot of Spearman correlation coefficients between PFAM distributions; from Arctic, Antarctic and other algal genomes (top) and transcriptomes (bottom) within the pan-genomic dataset. Violin plots are shown as per Fig. 3. Comparisons between members of the same taxonomic group, and involving either freshwater or obligately non-photosynthetic species were excluded from the analyses. Genomic calculations involving Antarctic species are not shown due to the presence of only one Antarctic genome (*Fragilariopsis cylindrus*) in the global dataset. Significance values of one-way ANOVA tests of the separation of means (red dots) are provided between Arctic-Arctic species pairs, and all other forms of species pairs considered.

**Fig. 3 - Figure Supplement 4.**
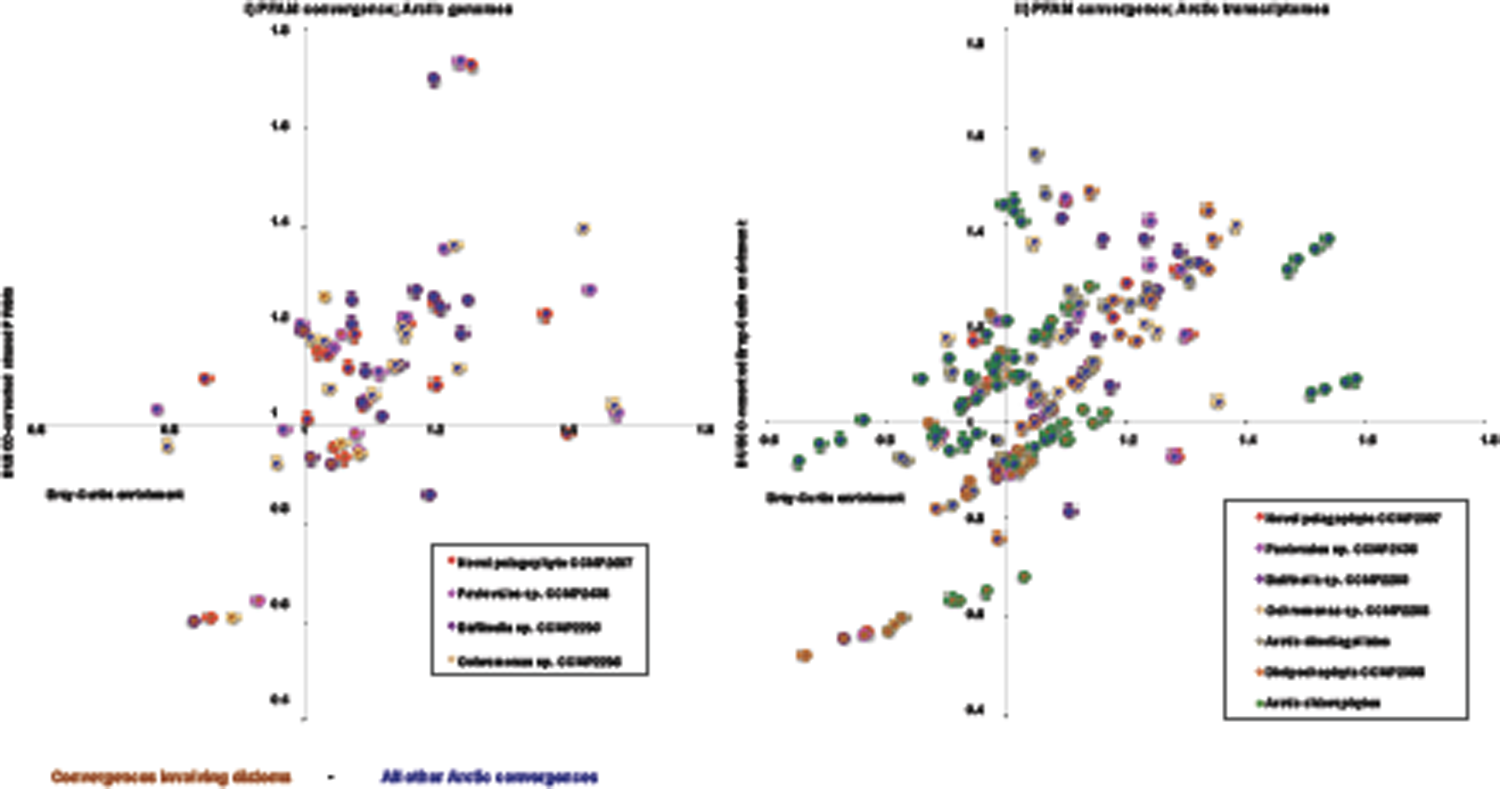
**Convergences in Arctic PFAM content principally involve non-diatom taxa**. This figure shows scatterplots of showing crude anti-Bray-Curtis values (horizontal axis), and BUSCO-normalised numbers of shared PFAMs (vertical axis), for similarity in PFAM content between different pairs of Arctic species in the pan-genomic dataset. Each value consists of the convergence value between a given Arctic query species and another Arctic reference, normalised against the mean Bray-Curtis or BUSCO-normalised Bray-Curtis value calculated between the query species against all non-Arctic species belonging to the same algal lineage as the reference. Values of >1 indicate a potential convergence in the PFAM content of pairs of Arctic species compared to phylogenetically equivalent references. Plots **(i)** and **(ii)** show respectively PFAM convergences involving query Arctic genomes; and PFAM convergences involving query Arctic transcriptomes from seven algal groups (chlorophytes, chrysophytes, cryptomonads, dictyochophytes, dinoflagellates, haptophytes and pelagophytes). The outer colour of each point is shaded by the taxonomic origin of the query species as per Fig. 1. The inner colour of each point is shaded by the taxonomic origin of the reference species: convergences involving diatom references are shaded brown, and convergences involving all other reference lineages are dark blue. In each plot, while multiple combinations of query and reference Arctic species are observed to have normalised convergence values > 1, suggesting Arctic-Arctic convergences, combinations involving diatom references strikingly have normalised reference values close to 1, or even less than 1, suggesting limited convergence in PFAM content between Arctic diatoms and other Arctic algal groups.

**Fig. 3 - Figure Supplement 5.**
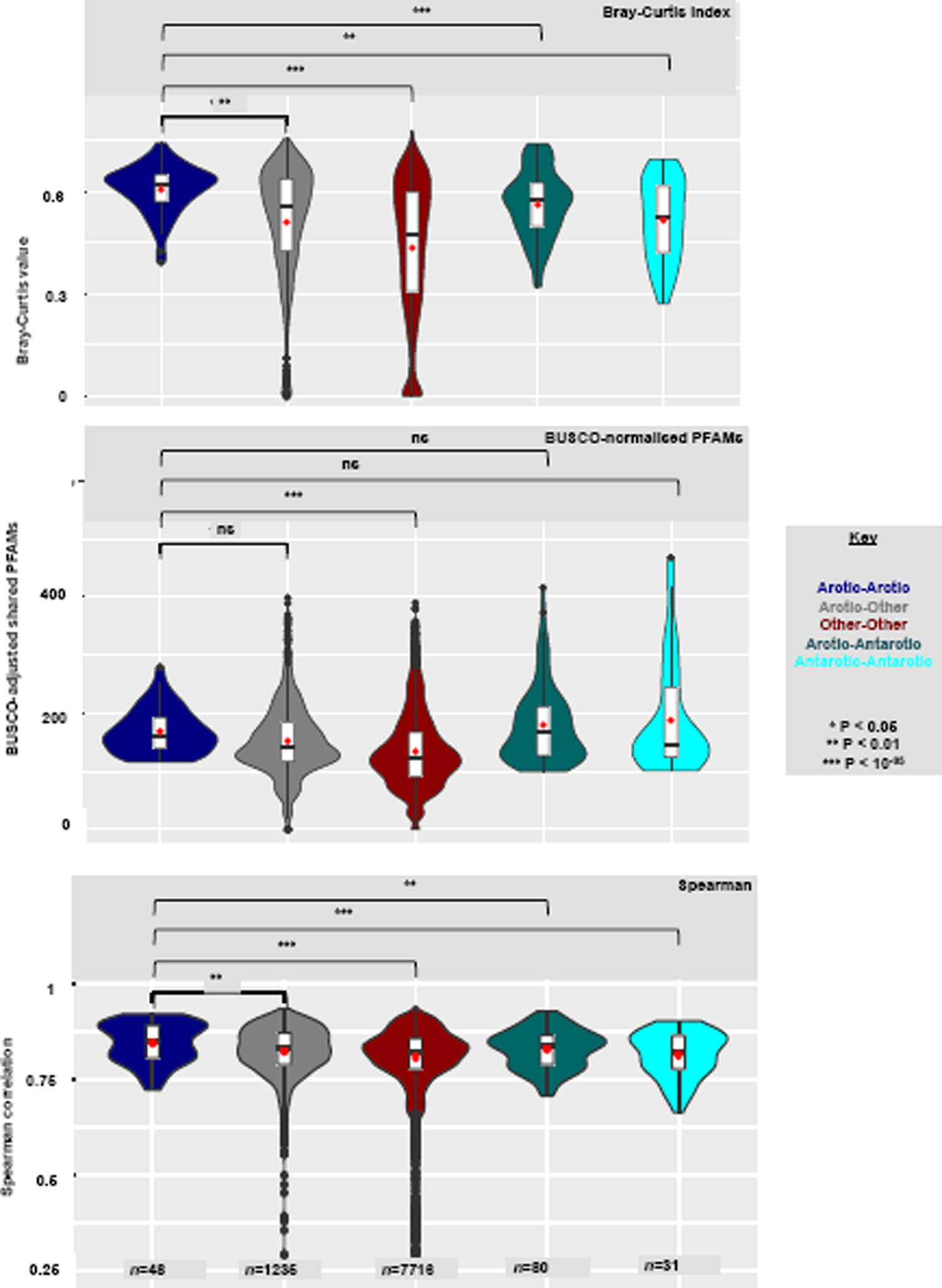
Violin plots of Arctic-Arctic convergence indices across transcriptome datasets exclusive of diatoms. Violin plots are shown as per Fig. 3. Diatom-exclusive genome plots are not shown due to the absence of Arctic diatom genomes from the pan-algal dataset.

**Fig. 4 - Figure Supplement 1.**
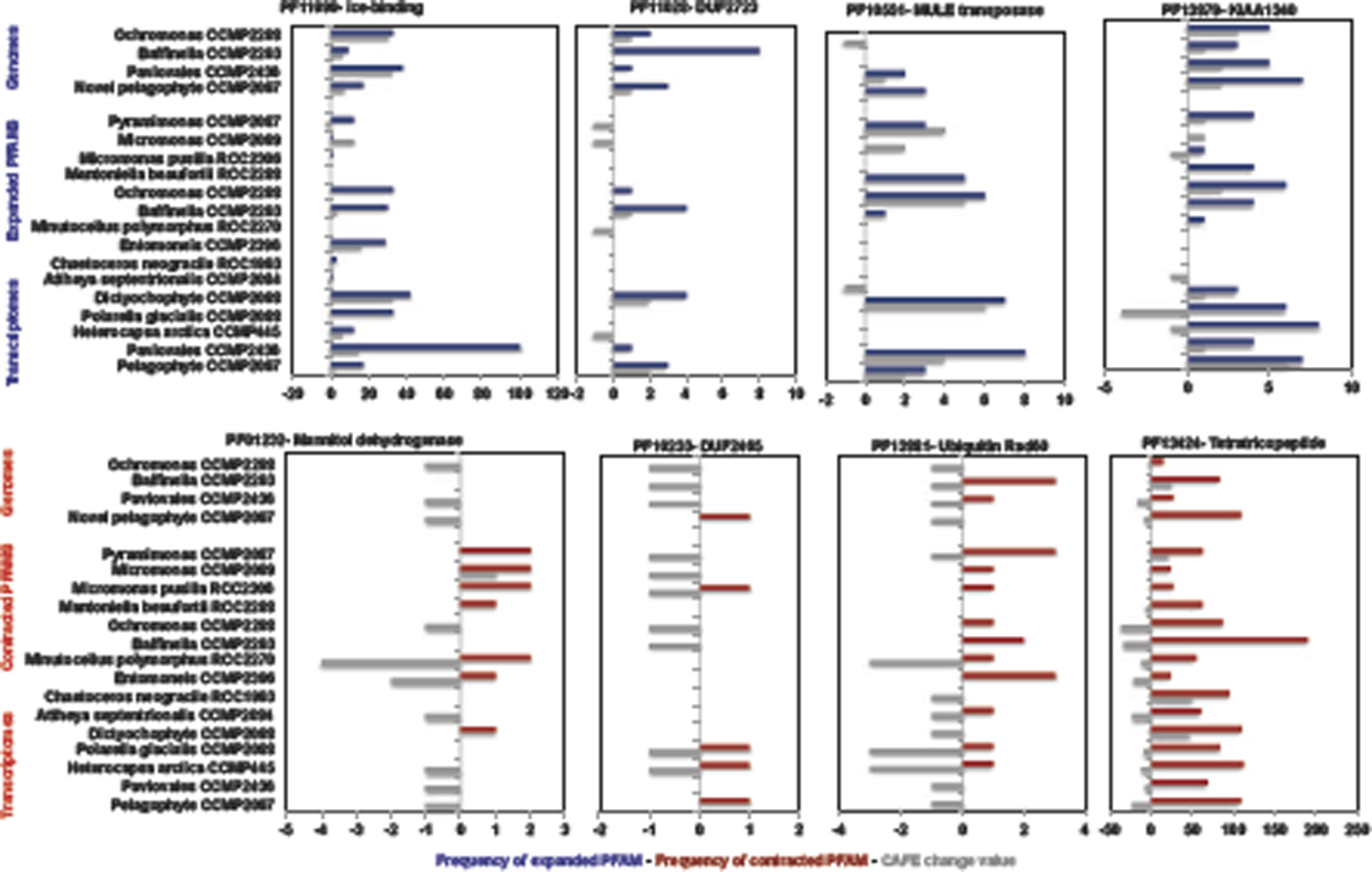
**Bar plots showing (coloured bars) the total number and (grey bars) CAFE scores**, across all studied Arctic species, for four PFAMs (blue) inferred to be significantly more frequently expanded and four PFAMs (red) significantly more contracted (one-tailed chi-squared test, P< 10^-05^) in Arctic compared to non-Arctic species across the pan-algal dataset.

**Fig. 5 - Figure Supplement 1.**
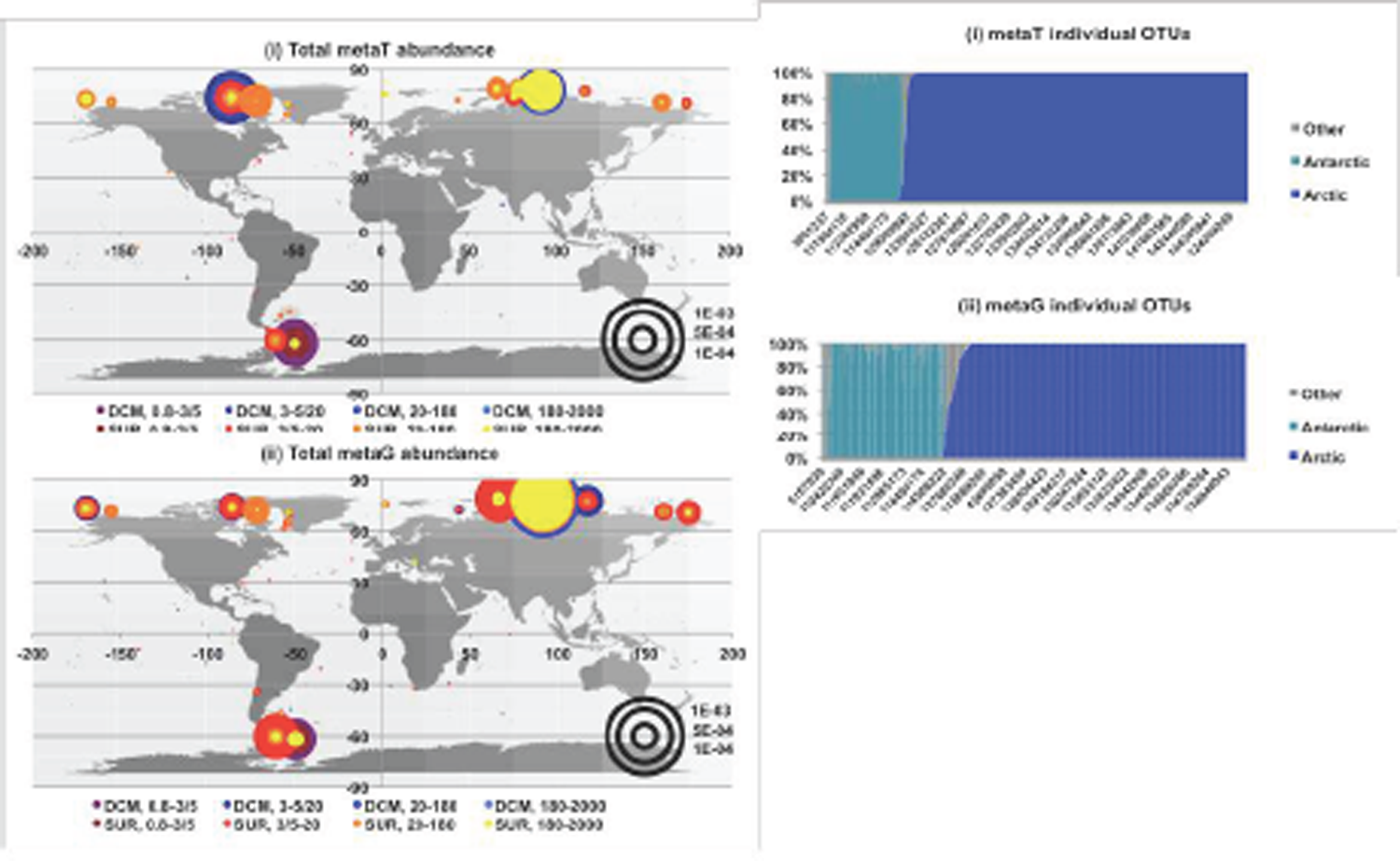
Environmental distributions of ice-binding domains in *Tara* Oceans data. (Left) global distribution plots, and (right) bar charts of individual proportional abundances of ice-binding domain (PF11999) containing sequences in (i) *Tara* Oceans meta-transcriptome and (ii) meta-genome sequence data, including the *Tara* Polar Circle survey. Stations are classified into Arctic (>60N), Antarctic (>55S) and Other latitudes, per Fig. 2. IBD-containing meta-genes show predominantly polar distributions, which are typically either restricted to the Arctic or Antarctic stations sampled, with only 4/ 1711 sampled meta-genes represented by >35% of their total global abundance in both Arctic and Antarctic libraries.

**Fig. 5 - Figure Supplement 2.**
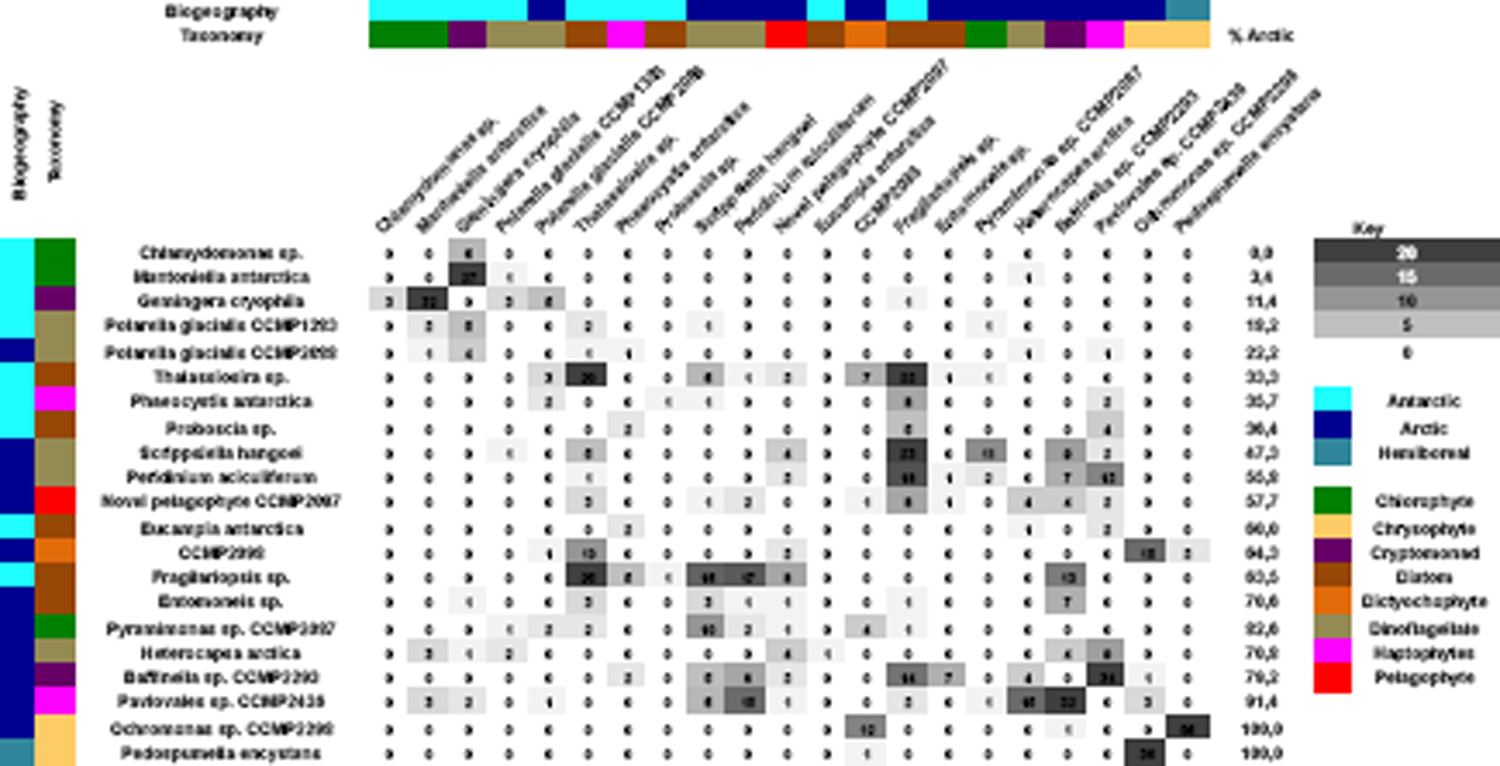
Internal BLAST best hit searches within the IBD alignment. This figure shows the number of times each Arctic- and Antarctic-native eukaryotic algae either retrieves, or is retrieved, as the best-scoring (lowest e-value) hit in a BLASTp search of the IBD alignment against itself (Table S3, sheets 3, 8, 9), excluding hits to members of the same genus (with *Scrippsiella* and *Peridinium* treated as congeneric; *38*) and Tara Oceans meta-genes. Sequences are shaded by taxonomy and biogeography as per Fig. 5, and are ranked in ascending order of the % of best-hits retrieved that are to Arctic eukaryotic algae. Species with fewer than three IBD sequences in the final alignment are not shown.

**Fig. 5 - Figure Supplement 3.**
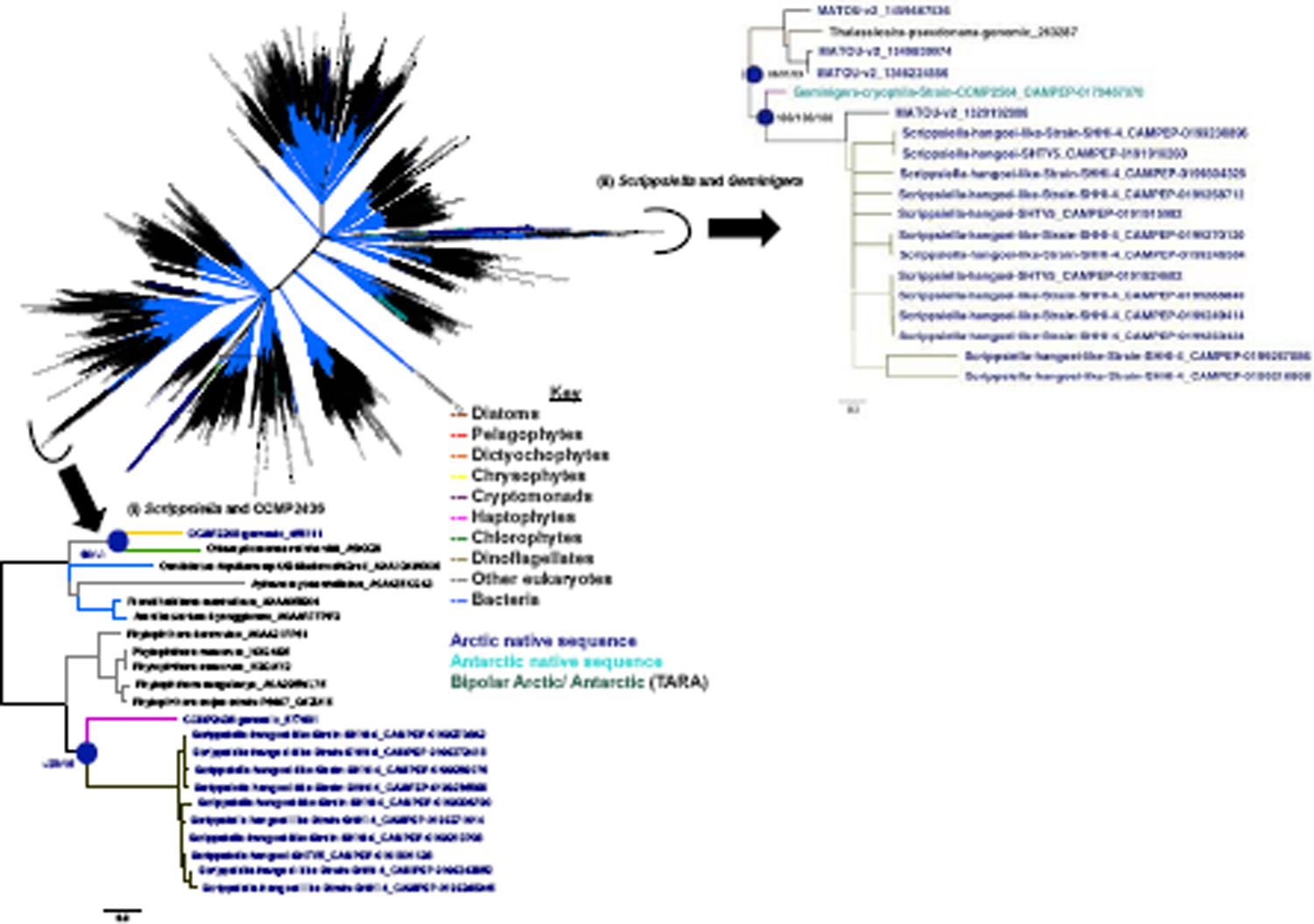
Diversity and polar horizontal gene transfers of DUF347 domains. This figure summarises the consensus best-scoring tree topology obtained with RaxML under GTR, JTT and WAG substitution models for a 3942 branch x 240 aa alignment of all DUF347 domains (PF03988) sampled from UniRef, jgi algal genomes, MMETSP, *Tara* Oceans, and an independent transcriptome survey of Baffin Island Northwater communities. Branches are shaded by evolutionary origin and leaf nodes by biogeography (either: isolation location of cultured accessions where recorded; or on oceanic regions for which > 70% total abundance of each Tara meta-gene could be recorded). Thick branches indicate presence of a clade in all three best-scoring tree outputs. Top: overview of the global topology obtained; bottom: magnified topology of a two clades containing the Arctic freshwater dinoflagellate *Scrippsiella* and **(i)** Pavlovales sp. CCMP2436, or **(ii)** the Antarctic cryptomonad *Geminigera*. Support values for two nodes linking *Scrippsiella, Geminigera* and multiple Arctic-only TARA meta-genes are shown with labelled circles.

**Fig. 5 - Figure Supplement 4.**
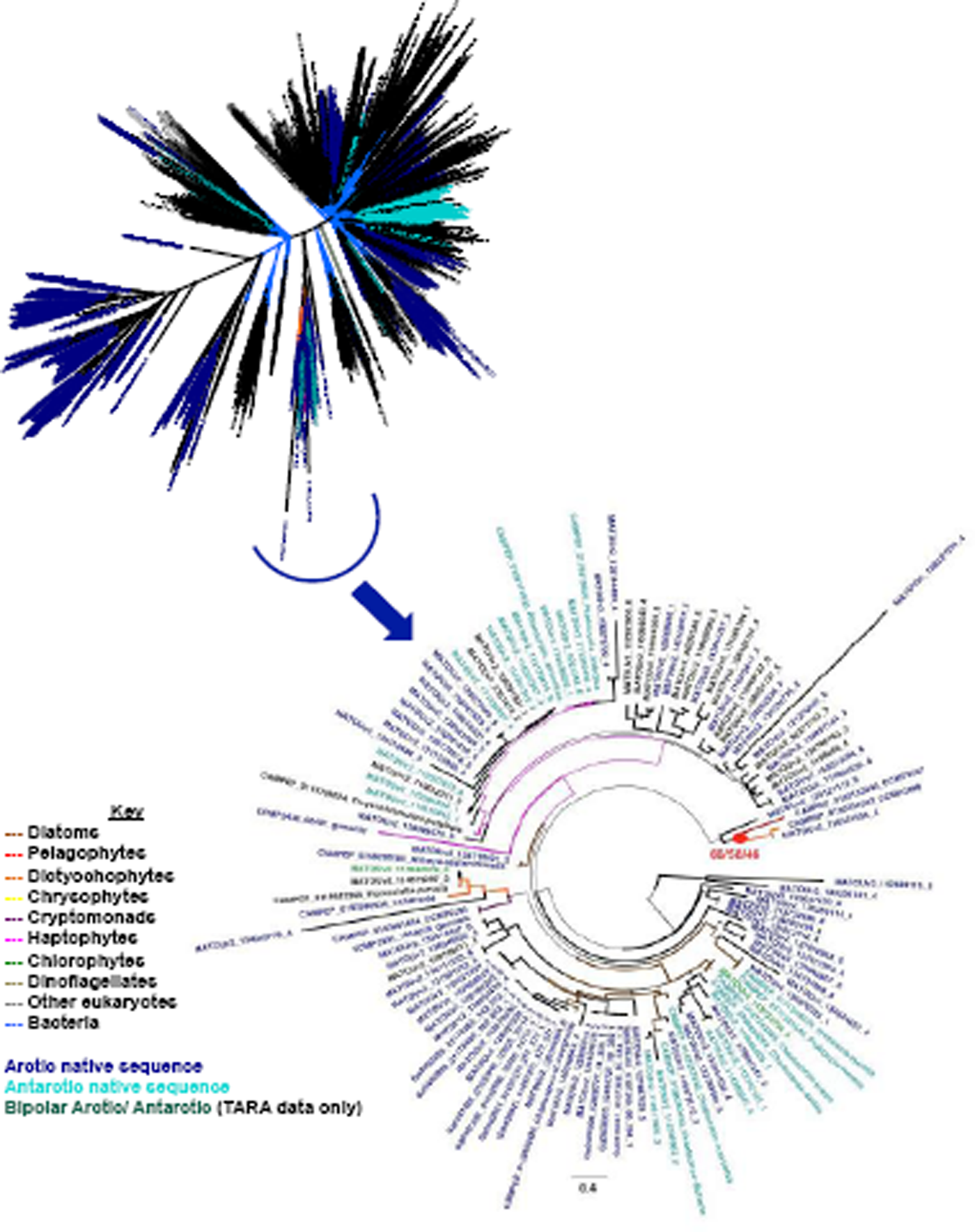
Diversity and polar horizontal gene transfers of PhnA domains. This figure summarises the consensus best-scoring tree topology obtained with RaxML under GTR, JTT and WAG substitution models for a 2934 branch x 113 aa alignment of all PhnA domains (PF03831) sampled from UniRef, jgi algal genomes, MMETSP, *Tara* Oceans, and an independent transcriptome survey of Baffin Island Northwater communities. Branches are shaded by evolutionary origin and leaf nodes by biogeography (either: isolation location of cultured accessions where recorded; or on oceanic regions for which > 70% total abundance of each *Tara* meta-gene could be recorded). Thick branches indicate presence of a clade in all three best-scoring tree outputs. Top: overview of the global topology obtained; bottom: magnified topology of a single clade showing evidence of horizontal gene transfer between Arctic- and Antarctic eukaryotic algae. A well-supported clade of Arctic pelagophyte (CCMP2097) and dictyochophyte (CCMP2098) sequences is shown with a red circle.

**Fig. 6 - Figure Supplement 1.**
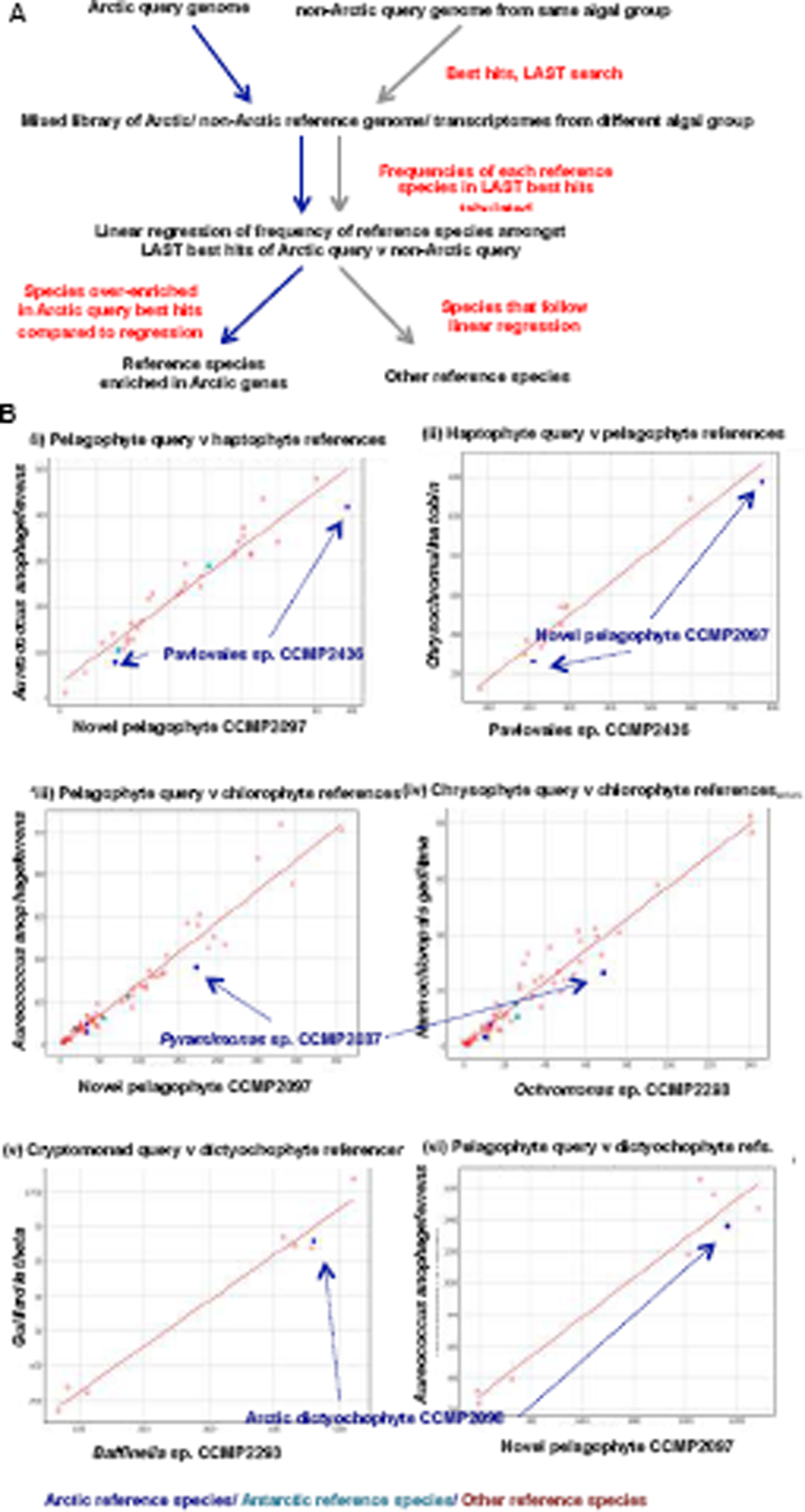
Identification of species enriched in Arctic-specific genes by LAST best hit search. A: Outline of the methodology used. Two query genomes (one Arctic, one not) from a query algal genome are searched against a mixed library of sequences from another algal group by LAST. As the query genomes are the same phylogenetic distance from the reference group each reference species should be retrieved as a LAST best hit a proportionate number of times by each query, with references that deviate from a linear relationship enriched in Arctic-specific genes. **B**: exemplar scatterplots of the number of LAST best hits obtained with Arctic (horizontal) and non-Arctic (vertical) queries against different algal groups; showing **(i, ii)** a bidirectional enrichment in LAST best hits between Pavlovales sp. CCMP2436 and the novel pelagophyte CCMP2097 in searches between haptophyte and pelagophyte libraries; **(iii, iv)** an enrichment in *Pyramimonas* sp. CCMP2087 in LAST best hits with Arctic pelagophyte and chrysophyte queries; and **(v, vi)** an enrichment in the Arctic dictychophyte CCMP2098 in LAST best hits with Arctic cryptomonad and pelagophyte queries.

**Fig. 6 - Figure Supplement 2.**
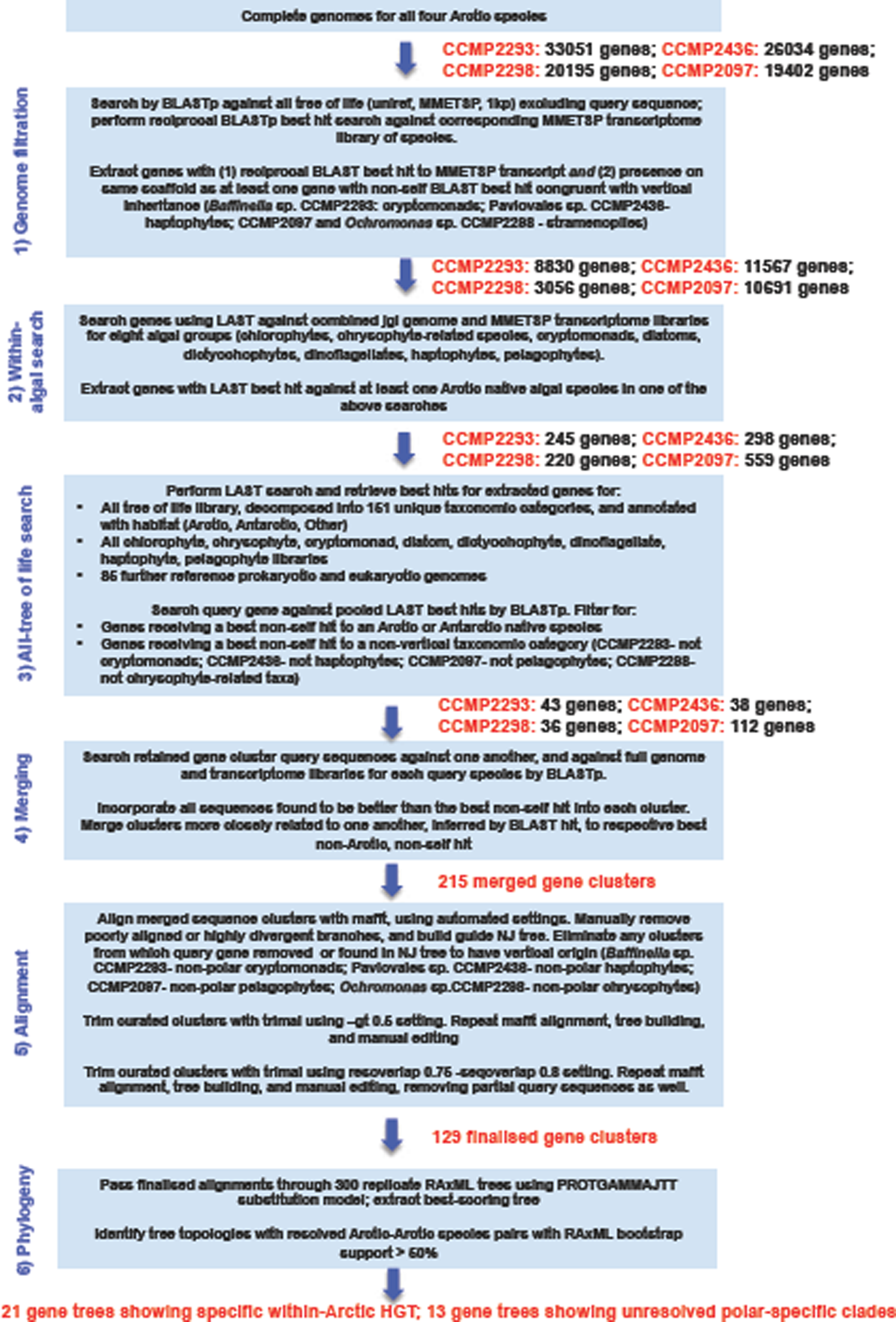
Pipeline of the phylogenomic approach used to identify within-Arctic HGTs in Arctic algal genomes.

**Fig. 6 - Figure Supplement 3.**
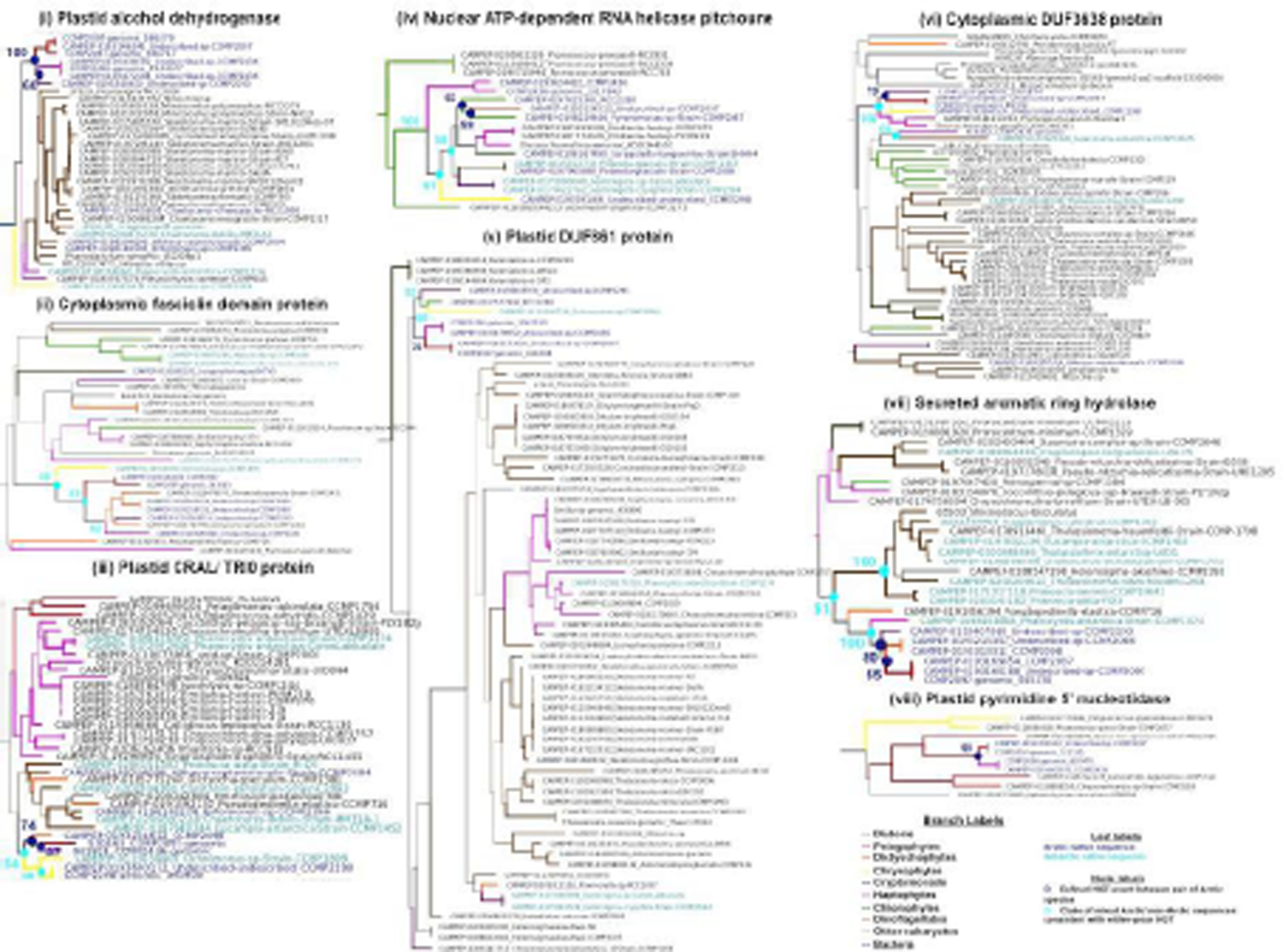
Eight exemplar within-Arctic HGT clusters identified by whole-genome phylogenetic profiling. The tree topologies shown consist of the proposed within Arctic-HGT clade and its two closest sister-groups; full tree topologies for these and 26 additional identified within-Arctic HGT trees are shown in Table S4, Sheet 9; and full nexus format tree topologies prepared using the same graphical format as detailed below are available through https://osf.io/3pmxb/files (Dorrell et al., 2021b) in «Supporting data > Exemplar within-Arctic HGT trees ». Branches are shaded by phylogenetic origin, leaves by biogeographical origin, and nodes resolving within-Arctic HGT events with > 50% bootstrap support are identified with coloured circles.

**Fig. 6 - Figure Supplement 4.**
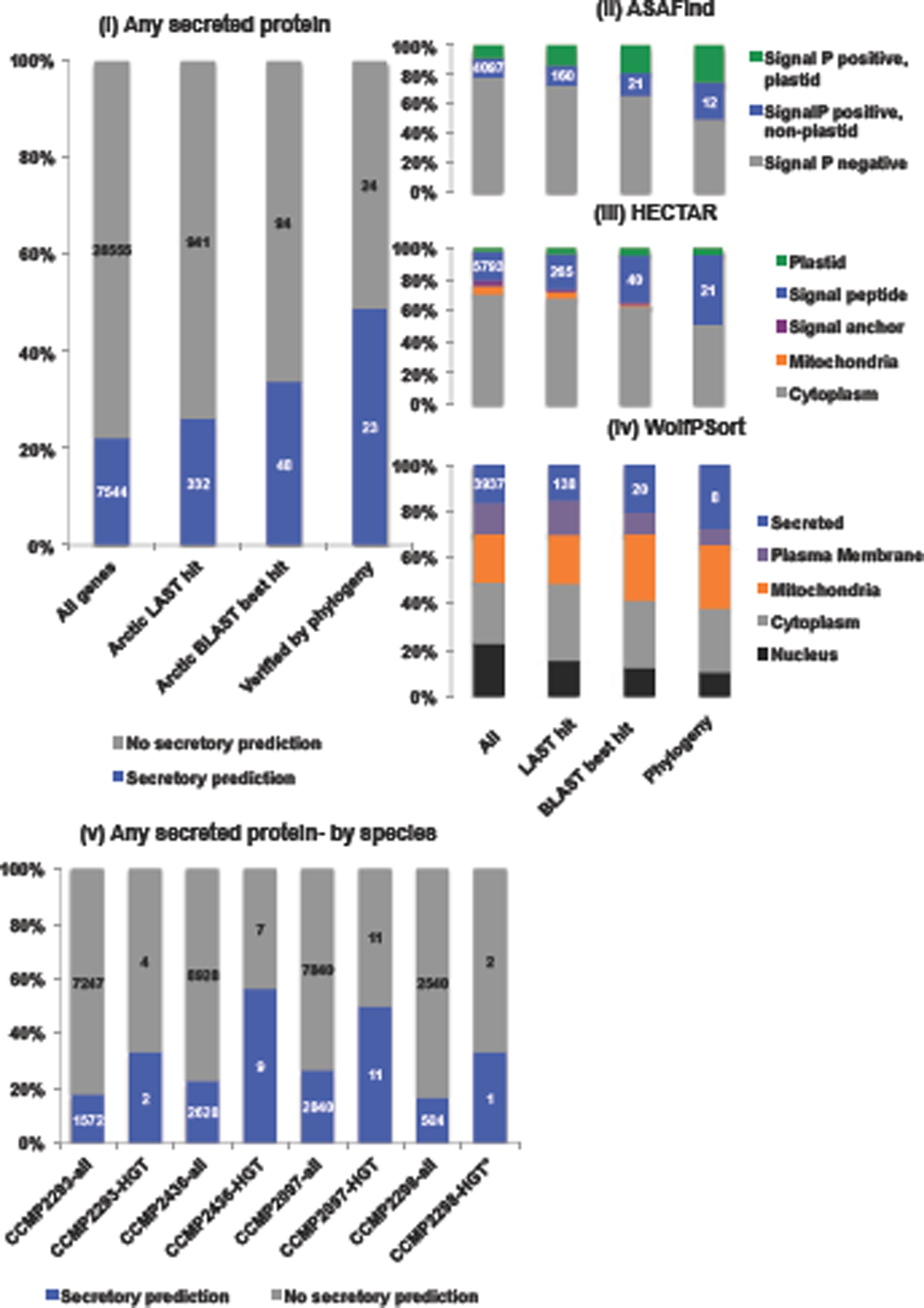
Secretory bias in within-Arctic HGTs. Bar charts showing **(i-iv)** the targeting predictions of all proteins in all Arctic algal genomes inspected for within-Arctic HGT; the total number of proteins yielding Arctic LAST or BLAST best hits per Fig. 6**- figure supplement 2**; and the total non-redundant proteins attributed to within Arctic-HGT; and **(v)** the total number of inspected proteins, and phylogenetically verified within-Arctic HGTs for each species. Targeting predictions were performed using ASAFind with SignalP v 3.0 (Gruber, Rocap, Kroth, Armbrust, & Mock, 2015), HECTAR under default conditions (Gschloessl, Guermeur, & Cock, 2008), and WolfPSort, with consensus animal, plant and fungal reference datasets (Horton et al., 2007) with proteins inferred to have a signal peptide and non-plastid prediction by either ASAFind or HECTAR, or an extracellular prediction by WolfPSort, inferred to be potentially secreted proteins.

**Fig. 6 - Figure Supplement 5.**
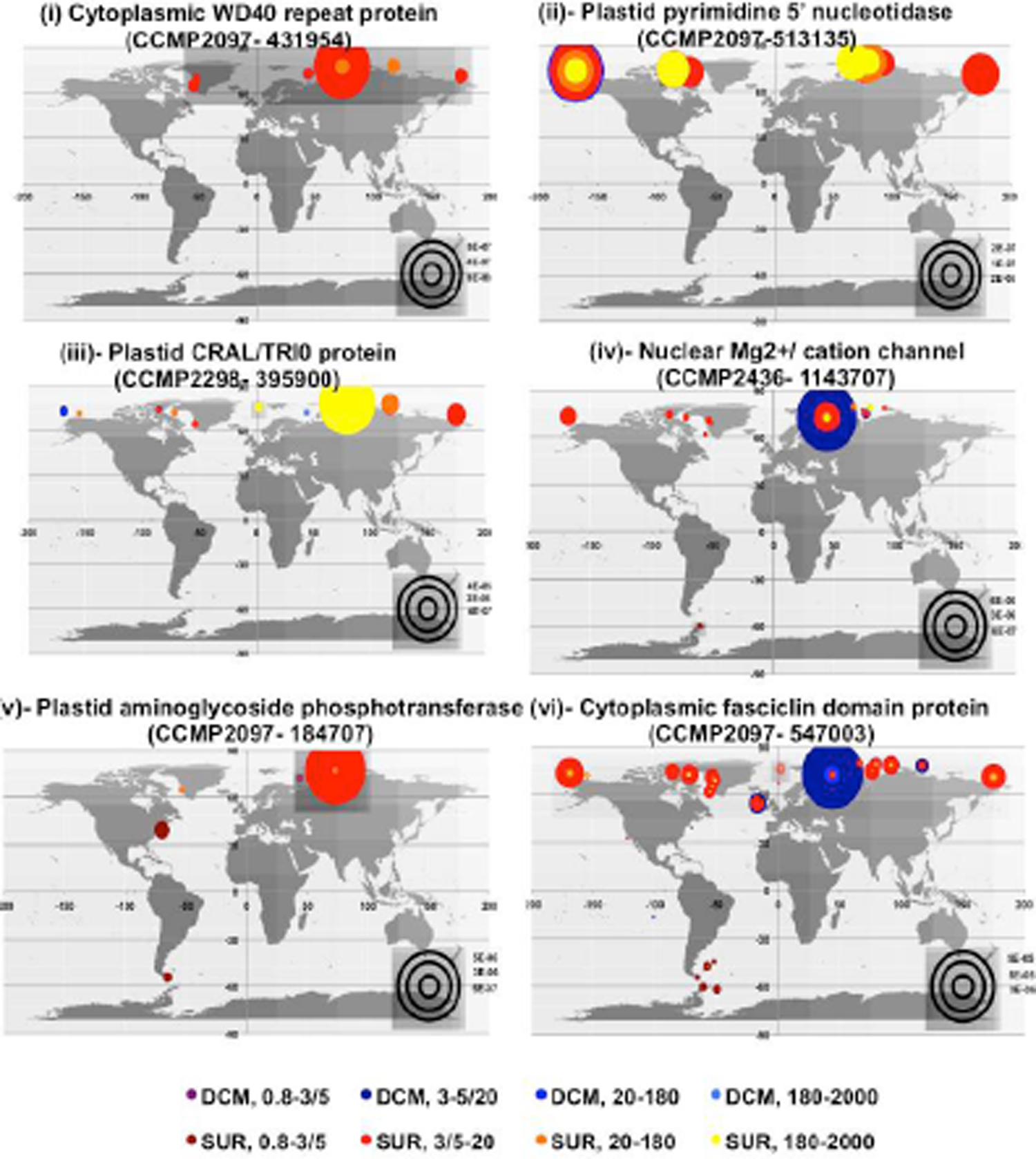
Environmental distributions of *Tara* Oceans meta-transcriptome homologues of six exemplar gene clusters phylogenetically inferred to have undergone within-Arctic HGT. This figure provides distributions for the environmental equivalents of within-Arctic HGT clade identified in Table S4, Sheet 9, as resolved by a combined hmmer, BLAST best-hit, MAFFT alignment and NJ tree pipeline. Total relative abundances are provided for all size fractions, and at both surface and deep chlorophyll maximum depth samples. Complete tabulated homologue sequences and abundances for all tractable genes in both meta-genome and meta-transcriptome data are presented in Table S4, sheets 10-11; and individual plots of total relative abundances and individual relative abundances for each phylogenetically reconciled gene are available through https://osf.io/3pmxb/files (Dorrell et al., 2021b) in «Supporting data > Within-Arctic HGTs > TARA distributions ».

## References

1. Anderson, E. L., Li, W., Klitgord, N., Highlander, S. K., Dayrit, M., Seguritan, V., … Jones, M. B. (2016). A robust ambient temperature collection and stabilization strategy: Enabling worldwide functional studies of the human microbiome. Sci Rep, 6, 31731. doi:10.1038/srep31731

2. Armbrust, E. V., Berges, J. A., Bowler, C., Green, B. R., Martinez, D., Putnam, N. H., … Rokhsar, D. S. (2004). The genome of the diatom *Thalassiosira pseudonana*: ecology, evolution, and metabolism. Science, 306(5693), 79–86.

3. Beisser, D., Graupner, N., Bock, C., Wodniok, S., Grossmann, L., Vos, M., … Boenigk, J. (2017). Comprehensive transcriptome analysis provides new insights into nutritional strategies and phylogenetic relationships of chrysophytes. PeerJ, 5, e2832. doi:10.7717/peerj.2832

4. Bendif, e. M., Probert, I., Hervé, A., Billard, C., Goux, D., Lelong, C., … Véron, B. (2011). Integrative taxonomy of the Pavlovophyceae (Haptophyta): a reassessment. Protist, 162(5), 738–761. doi:10.1016/j.protis.2011.05.001

5. Beszczynska-Moller, A., Woodgate, R. A., Lee, C., Melling, H., & Karcher, M. (2011). A synthesis of exchanges through the main oceanic gateways to the Arctic Ocean. Oceanography, 24(3), 82–99. doi:10.5670/oceanog.2011.59

6. Bock, N. A., Charvet, S., Burns, J., Gyaltshen, Y., Rozenberg, A., Duhamel, S., & Kim, E. (2021). Experimental identification and in silico prediction of bacterivory in green algae. ISME J. doi:10.1038/s41396-021-00899-w

7. Brinkmeyer, R., Knittel, K., Jürgens, J., Weyland, H., Amann, R., & Helmke, E. (2003). Diversity and structure of bacterial communities in Arctic versus Antarctic pack ice. Appl Environ Microbiol, 69(11), 6610–6619.

8. Buchfink, B., Xie, C., & Huson, D. H. (2015). Fast and sensitive protein alignment using DIAMOND. Nat Methods, 12(1), 59–60. doi:10.1038/nmeth.3176

9. Burki, F., Kaplan, M., Tikhonenkov, D. V., Zlatogursky, V., Minh, B. Q., Radaykina, L. V., … Keeling, P. J. (2016). Untangling the early diversification of eukaryotes: a phylogenomic study of the evolutionary origins of Centrohelida, Haptophyta and Cryptista. Proc Biol Sci, 283(1823). doi:10.1098/rspb.2015.2802

10. Cao, S., Zhang, W., Ding, W., Wang, M., Fan, S., Yang, B., … Zhang, Y.-Z. (2020). Structure and function of the Arctic and Antarctic marine microbiota as revealed by metagenomics. Microbiome, 8(1), 47. doi:10.1186/s40168-020-00826-9

11. Capella-Gutiérrez, S., Silla-Martínez, J. M., & Gabaldón, T. (2009). trimAl: a tool for automated alignment trimming in large-scale phylogenetic analyses. Bioinformatics, 25(15), 1972–1973. doi:10.1093/bioinformatics/btp348

12. Carmack, E. C. (2007). The alpha/beta ocean distinction: a perspective on freshwater fluxes, convection, nutrients and productivity in high-latitude seas. Deep-Sea Res. Part II, 54(23-26), 21.

13. Carpenter, E. J., Matasci, N., Ayyampalayam, S., Wu, S., Sun, J., Yu, J., … Wong, G. K. (2019). Access to RNA-sequencing data from 1,173 plant species: the 1000 Plant transcriptomes initiative (1KP). Gigascience, 8(10). doi:10.1093/gigascience/giz126

14. Carradec, Q., Pelletier, E., Da Silva, C., Alberti, A., Seeleuthner, Y., Blanc-Mathieu, R., … Coordinators, T. O. (2018). A global ocean atlas of eukaryotic genes. Nat Commun, 9(1), 373. doi:10.1038/s41467-017-02342-1

15. Cenci, U., Sibbald, S. J., Curtis, B. A., Kamikawa, R., Eme, L., Moog, D., … Archibald, J. M. (2018). Nuclear genome sequence of the plastid-lacking cryptomonad *Goniomonas avonlea* provides insights into the evolution of secondary plastids. BMC Biol, 16(1), 137. doi:10.1186/s12915-018-0593-5

16. Chetouani, F., Glaser, P., & Kunst, F. (2001). FindTarget: software for subtractive genome analysis. Microbiology (Reading), 147(Pt 10), 2643–2649. doi:10.1099/00221287-147-10-2643

17. Choi, I. G., & Kim, S. H. (2007). Global extent of horizontal gene transfer. Proc Natl Acad Sci USA, 104(11), 4489–4494. doi:10.1073/pnas.0611557104

18. Cid, F. P., Maruyama, F., Murase, K., Graether, S. P., Larama, G., Bravo, L. A., & Jorquera, M. A. (2018). Draft genome sequences of bacteria isolated from the *Deschampsia antarctica* phyllosphere. Extremophiles, 22(3), 537–552. doi:10.1007/s00792-018-1015-x

19. Craveiro, S. C., Daugbjerg, N., Moestrup, Ø., & Calado, A. J. (2017). Studies on *Peridinium aciculiferum* and *Peridinium malmogiense* (=*Scrippsiella hangoei*): comparison with *Chimonodinium lomnickii* and description of *Apocalathium* gen. nov. (Dinophyceae). Phycologia, 56(1), 21–35. doi:10.2216/16-20.1

20. Cummins, C. A., & McInerney, J. O. (2011). A method for inferring the rate of evolution of homologous characters that can potentially improve phylogenetic inference, resolve deep divergence and correct systematic biases. Syst Biol, 60(6), 833–844. doi:10.1093/sysbio/syr064

21. Curtis, B. A., Tanifuji, G., Burki, F., Gruber, A., Irimia, M., Maruyama, S., … Archibald, J. M. (2012). Algal genomes reveal evolutionary mosaicism and the fate of nucleomorphs. Nature, 492(7427), 59–65. doi:10.1038/nature11681

22. Daugbjerg, N., Norlin, A., & Lovejoy, C. (2018). *Baffinella frigidus* gen. et sp. nov. (Baffinellaceae fam. nov., Cryptophyceae) from Baffin Bay: morphology, pigment profile, phylogeny, and growth rate response to three abiotic factors. J Phycol, 54(5), 665–680. doi:10.1111/jpy.12766

23. Davis, A. K., Hildebrand, M., & Palenik, B. (2006). Gene expression induced by copper stress in the diatom *Thalassiosira pseudonana*. Eukaryot Cell, 5(7), 1157–1168. doi:10.1128/EC.00042-06

24. De Bie, T., Cristianini, N., Demuth, J. P., & Hahn, M. W. (2006). CAFE: a computational tool for the study of gene family evolution. Bioinformatics, 22(10), 1269–1271. doi:10.1093/bioinformatics/btl097

25. de Vargas, C., Audic, S., Henry, N., Decelle, J., Mahé, F., Logares, R., … Coordinators, T. O. (2015). Ocean plankton. Eukaryotic plankton diversity in the sunlit ocean. Science, 348(6237), 1261605. doi:10.1126/science.1261605

26. Derelle, E., Ferraz, C., Rombauts, S., Rouzé, P., Worden, A. Z., Robbens, S., … Moreau, H. (2006). Genome analysis of the smallest free-living eukaryote *Ostreococcus tauri* unveils many unique features. Proc Natl Acad Sci U S A, 103(31), 11647–11652. doi:10.1073/pnas.0604795103

27. Dorrell, R. G., Azuma, T., Nomura, M., Audren de Kerdrel, G., Paoli, L., Yang, S., … Kamikawa, R. (2019). Principles of plastid reductive evolution illuminated by nonphotosynthetic chrysophytes. Proc Natl Acad Sci USA, 116(14), 6914–6923. doi:10.1073/pnas.1819976116

28. Dorrell, R. G., Klinger, C. M., Newby, R. J., Butterfield, E. R., Richardson, E., Dacks, J. B., … Bowler, C. (2017). Progressive and biased divergent evolution underpins the origin and diversification of peridinin dinoflagellate plastids. Mol Biol Evol, 34(2), 361–379. doi:10.1093/molbev/msw235

29. Dorrell, R. G., Villain, A., Perez-Lamarque, B., Audren de Kerdrel, G., McCallum, G., Watson, A. K., … Blanc, G. (2021a). Phylogenomic fingerprinting of tempo and functions of horizontal gene transfer within ochrophytes. Proc Natl Acad Sci USA, 118(4). doi:10.1073/pnas.2009974118

30. Dorrell, R. G., Lovejoy, C., Zarevski, N., Bowler, C., Kuo, A., Grigoriev, I., … Dacks, J. (2021b). Arctic algal genomes supporting data. https://osf.io/3pmxb/.

31. Eegeesiak, O., Aariak, E., & Kleist, K. (2017). People of the ice bridge: the future of the Pikialasorsuaq. Report of the Pikialasorsuaq Commission, Inuit Circumpolar Council Canada, Ottawa.

32. Eme, L., Gentekaki, E., Curtis, B., Archibald, J. M., & Roger, A. J. (2017). Lateral gene transfer in the adaptation of the anaerobic parasite *Blastocystis* to the gut. Curr Biol, 27(6), 807–820. doi:10.1016/j.cub.2017.02.003

33. Emms, D. M., & Kelly, S. (2019). OrthoFinder: phylogenetic orthology inference for comparative genomics. Genome Biol, 20(1), 238. doi:10.1186/s13059-019-1832-y

34. Furnholm, T. R., & Tisa, L. S. (2014). The ins and outs of metal homeostasis by the root nodule actinobacterium Frankia. BMC Genomics, 15, 1092. doi:10.1186/1471-2164-15-1092

35. Gachon, C. M. M., Heesch, S., Kuepper, F. C., Achilles-Day, U. E. M., Brennan, D., Campbell, C. N., … Day, J. G. (2013). The CCAP KnowledgeBase: linking protistan and cyanobacterial biological resources with taxonomic and molecular data. Systematics and Biodiversity, 11(4), 407–413. doi:10.1080/14772000.2013.859641

36. Gast, R. J., Moran, D. M., Dennett, M. R., & Caron, D. A. (2007). Kleptoplasty in an Antarctic dinoflagellate: caught in evolutionary transition? Environ Microbiol, 9(1), 39–45. doi:10.1111/j.1462-2920.2006.01109.x

37. Gnerre, S., Maccallum, I., Przybylski, D., Ribeiro, F. J., Burton, J. N., Walker, B. J., … Jaffe, D. B. (2011). High-quality draft assemblies of mammalian genomes from massively parallel sequence data. Proc Natl Acad Sci U S A, 108(4), 1513–1518. doi:10.1073/pnas.1017351108

38. Gobler, C. J., Berry, D. L., Dyhrman, S. T., Wilhelm, S. W., Salamov, A., Lobanov, A. V., … Grigoriev, I. V. (2011). Niche of harmful alga *Aureococcus anophagefferens* revealed through ecogenomics. Proc Natl Acad Sci USA 108(11), 4352–4357. doi:10.1073/pnas.1016106108

39. Grigoriev, I. V., Hayes, R. D., Calhoun, S., Kamel, B., Wang, A., Ahrendt, S., … Kuo, A. (2021). PhycoCosm, a comparative algal genomics resource. Nucleic Acids Res, 49(D1), D1004–D1011. doi:10.1093/nar/gkaa898

40. Grossmann, L., Bock, C., Schweikert, M., & Boenigk, J. (2016). Small but manifold - hidden diversity in “*Spumella*-like flagellates”. J Eukaryot Microbiol, 63(4), 419–439. doi:10.1111/jeu.12287

41. Gruber, A., Rocap, G., Kroth, P. G., Armbrust, E. V., & Mock, T. (2015). Plastid proteome prediction for diatoms and other algae with secondary plastids of the red lineage. Plant J, 81(3), 519–528. doi:10.1111/tpj.12734

42. Gschloessl, B., Guermeur, Y., & Cock, J. M. (2008). HECTAR: a method to predict subcellular targeting in heterokonts. BMC Bioinformatics, 9, 393. doi:10.1186/1471-2105-9-393

43. Guiry, M. D., Guiry, G. M., Morrison, L., Rindi, F., Valenzuala Miranda, S., Mathieson, A. C., … Garbary, D. J. (2014). AlgaeBase: an on-line resource for algae. *Cryptogamie*, Algologie, 35(2), 11.

44. Hamilton, A. K., Lovejoy, C., Galand, P. E., & Ingram, R. G. (2008). Water masses and biogeography of picoeukaryote assemblages in a cold hydrographically complex system. Limnology and Oceanography, 53(3), 922–935. doi:10.4319/lo.2008.53.3.0922

45. Han, K. Y., Graf, L., Reyes, C. P., Melkonian, B., Andersen, R. A., Yoon, H. S., & Melkonian, M. (2018). A re-investigation of *Sarcinochrysis marina* (Sarcinochrysidales, Pelagophyceae) from its type locality and the descriptions of *Arachnochrysis, Pelagospilus, Sargassococcus* and *Sungminbooa genera* nov. Protist, 169(1), 79–106. doi:10.1016/j.protis.2017.12.004

46. Horn, H., Slaby, B. M., Jahn, M. T., Bayer, K., Moitinho-Silva, L., Förster, F., … Hentschel, U. (2016). An enrichment of CRISPR and other defense-related features in marine sponge-associated microbial metagenomes. Front Microbiol, 7, 1751. doi:10.3389/fmicb.2016.01751

47. Horton, P., Park, K. J., Obayashi, T., Fujita, N., Harada, H., Adams-Collier, C. J., & Nakai, K. (2007). WoLF PSORT: protein localization predictor. Nucleic Acids Research, 35, W585–W587. doi:10.1093/nar/gkm259

48. Hovde, B. T., Deodato, C. R., Hunsperger, H. M., Ryken, S. A., Yost, W., Jha, R. K., … Cattolico, R. A. (2015). Genome sequence and transcriptome Analyses of *Chrysochromulina tobin*: metabolic tools for enhanced algal fitness in the prominent order Prymnesiales (Haptophyceae). PLoS Genetics, 11(9). doi:10.1371/journal.pgen.1005469

49. Ibarbalz, F. M., Henry, N., Brandão, M. C., Martini, S., Busseni, G., Byrne, H., … Coordinators, T. O. (2019). Global trends in marine plankton diversity across kingdoms of life. Cell, 179(5), 1084–1097.e1021. doi:10.1016/j.cell.2019.10.008

50. Initiative, O. T. P. T. (2019). One thousand plant transcriptomes and the phylogenomics of green plants. Nature, 574(7780), 679–685. doi:10.1038/s41586-019-1693-2

51. Irwin, N., Pittis, A., Richards, T., & Keeling, P. (2021). Viral-eukaryotic gene exchange drives infection mode and cellular evolution. ResearchSquare, preprint, 380297. doi:10.21203/rs.3.rs-380297/v1

52. Jeong, H., Kang, H., Lim, A., Jang, S., Lee, K., Lee, S., … Kim, K. (2021). Feeding diverse prey as an excellent strategy of mixotrophic dinoflagellates for global dominance. Sci Adv., 7(2), eabe4214.

53. Joli, N., Monier, A., Logares, R., & Lovejoy, C. (2017). Seasonal patterns in Arctic prasinophytes and inferred ecology of *Bathycoccus* unveiled in an Arctic winter metagenome. ISME J, 11(6), 1372–1385. doi:10.1038/ismej.2017.7

54. Jones, P., Binns, D., Chang, H. Y., Fraser, M., Li, W., McAnulla, C., … Hunter, S. (2014). InterProScan 5: genome-scale protein function classification. Bioinformatics, 30(9), 1236–1240. doi:10.1093/bioinformatics/btu031

55. Kanehisa, M., & Sato, Y. (2020). KEGG Mapper for inferring cellular functions from protein sequences. Protein Sci, 29(1), 28–35. doi:10.1002/pro.3711

56. Katoh, K., Rozewicki, J., & Yamada, K. D. (2017). MAFFT online service: multiple sequence alignment, interactive sequence choice and visualization. Brief Bioinform. doi:10.1093/bib/bbx108

57. Keeling, P. J., Burki, F., Wilcox, H. M., Allam, B., Allen, E. E., Amaral-Zettler, L. A., … Worden, A. Z. (2014). The Marine Microbial Eukaryote Transcriptome Sequencing Project (MMETSP): illuminating the functional diversity of eukaryotic life in the oceans through transcriptome sequencing. PLoS Biol, 12(6), e1001889. doi:10.1371/journal.pbio.1001889

58. Kiełbasa, S. M., Wan, R., Sato, K., Horton, P., & Frith, M. C. (2011). Adaptive seeds tame genomic sequence comparison. Genome Res, 21(3), 487–493. doi:10.1101/gr.113985.110

59. Kulakova, A. N., Kulakov, L. A., Akulenko, N. V., Ksenzenko, V. N., Hamilton, J. T., & Quinn, J. P. (2001). Structural and functional analysis of the phosphonoacetate hydrolase (phnA) gene region in *Pseudomonas fluorescens* 23F. J Bacteriol, 183(11), 3268–3275. doi:10.1128/JB.183.11.3268-3275.2001

60. Kumar, S., Stecher, G., Li, M., Knyaz, C., & Tamura, K. (2018). MEGA X: Molecular Evolutionary Genetics Analysis across computing platforms. Mol Biol Evol, 35(6), 1547–1549. doi:10.1093/molbev/msy096

61. Kuo, A., Bushnell, B., & Grigoriev, I. V. (2014). Chapter One - Fungal Genomics: sequencing and annotation. In F. M. Martin (Ed.), Advances in Botanical Research (Vol. 70, pp. 1–52): Academic Press.

62. Leu, E., Mundy, C. J., Assmy, P., Campbell, K., Gabrielsen, T. M., Gosselin, M., … Gradinger, R. (2015). Arctic spring awakening - steering principles behind the phenology of vernal ice algal blooms. Progress in Oceanography, 139, 151–170. doi:10.1016/j.pocean.2015.07.012

63. Li, W. K., McLaughlin, F. A., Lovejoy, C., & Carmack, E. C. (2009). Smallest algae thrive as the Arctic Ocean freshens. Science, 326(5952), 539. doi:10.1126/science.1179798

64. Liang, Y., Koester, J. A., Liefer, J. D., Irwin, A. J., & Finkel, Z. V. (2019). Molecular mechanisms of temperature acclimation and adaptation in marine diatoms. ISME J, 13(10), 2415–2425. doi:10.1038/s41396-019-0441-9

65. Lie, A. A. Y., Liu, Z., Terrado, R., Tatters, A. O., Heidelberg, K. B., & Caron, D. A. (2018). A tale of two mixotrophic chrysophytes: Insights into the metabolisms of two *Ochromonas* species (Chrysophyceae) through a comparison of gene expression. PLoS One, 13(2), e0192439. doi:10.1371/journal.pone.0192439

66. Lin, S., Cheng, S., Song, B., Zhong, X., Lin, X., Li, W., … Morse, D. (2015). The *Symbiodinium kawagutii* genome illuminates dinoflagellate gene expression and coral symbiosis. Science, 350(6261), 691–694. doi:10.1126/science.aad0408

67. Lommer, M., Specht, M., Roy, A. S., Kraemer, L., Andreson, R., Gutowska, M. A., … LaRoche, J. (2012). Genome and low-iron response of an oceanic diatom adapted to chronic iron limitation. Genome Biology, 13(7). doi:10.1186/gb-2012-13-7-r66

68. Longhurst, A. (2006). *Ecological Geography of the Sea* (2nd ed.): Academic Press.

69. Lovejoy, C., Vincent, W. F., Bonilla, S., Roy, S., Martineau, M. J., Terrado, R., … Pedros-Alio, C. (2007). Distribution, phylogeny, and growth of cold-adapted picoprasinophytes in arctic seas. Journal of Phycology, 43(1), 78–89. doi:10.1111/j.1529-8817.2006.00310.x

70. Marron, A. O., Ratcliffe, S., Wheeler, G. L., Goldstein, R. E., King, N., Not, F., … Richter, D. J. (2016). The evolution of silicon transport in eukaryotes. Molecular Biology and Evolution, 33(12), 3226–3248. doi:10.1093/molbev/msw209

71. Martin, J. A., & Wang, Z. (2011). Next-generation transcriptome assembly. Nature Reviews Genetics, 12(10), 671–682. doi:10.1038/nrg3068

72. McKie-Krisberg, Z. M., & Sanders, R. W. (2014). Phagotrophy by the picoeukaryotic green alga *Micromonas*: implications for Arctic Oceans. ISME J, 8(10), 1953–1961. doi:10.1038/ismej.2014.16

73. Metpally, R. P., & Reddy, B. V. (2009). Comparative proteome analysis of psychrophilic versus mesophilic bacterial species: insights into the molecular basis of cold adaptation of proteins. BMC Genomics, 10, 11. doi:10.1186/1471-2164-10-11

74. Miller, M. A., Schwartz, T., Pickett, B. E., He, S., Klem, E. B., Scheuermann, R. H., … O’Leary, M. A. (2015). A RESTful API for access to phylogenetic tools via the CIPRES Science Gateway. Evol Bioinform Online, 11, 43–48. doi:10.4137/EBO.S21501

75. Mistry, J., Chuguransky, S., Williams, L., Qureshi, M., Salazar, G. A., Sonnhammer, E. L. L., … Bateman, A. (2020). PFAM: the protein families database in 2021. Nucleic Acids Res. doi:10.1093/nar/gkaa913

76. Mock, T., Otillar, R. P., Strauss, J., McMullan, M., Paajanen, P., Schmutz, J., … Grigoriev, I. V. (2017). Evolutionary genomics of the cold-adapted diatom *Fragilariopsis cylindrus*. Nature, 541(7638), 536–540. doi:10.1038/nature20803

77. Muñoz-Villagrán, C. M., Mendez, K. N., Cornejo, F., Figueroa, M., Undabarrena, A., Morales, E. H., … Vásquez, C. C. (2018). Comparative genomic analysis of a new tellurite-resistant *Psychrobacter* strain isolated from the Antarctic Peninsula. PeerJ, 6, e4402. doi:10.7717/peerj.4402

78. Nelson, D. R., Hazzouri, K. M., Lauersen, K. J., Jaiswal, A., Chaiboonchoe, A., Mystikoi, A., … Salehi-Ashtiani, K. (2021). Large-scale genome sequencing reveals the driving forces of viruses in microalgal evolution. Cell Host and Microbe,29, 250–266. doi:10.1016/j.chom.2020.12.005

79. Nguyen, L. T., Schmidt, H. A., von Haeseler, A., & Minh, B. Q. (2015). IQ-TREE: a fast and effective stochastic algorithm for estimating maximum-likelihood phylogenies. Mol Biol Evol, 32(1), 268–274. doi:10.1093/molbev/msu300

80. Notz, D., & Stroeve, J. (2016). Observed Arctic sea-ice loss directly follows anthropogenic CO2 emission. Science, 354(6313), 747–750. doi:10.1126/science.aag2345

81. Pesant, S., Not, F., Picheral, M., Kandels-Lewis, S., Le Bescot, N., Gorsky, G., … Tara Oceans, C. (2015). Open science resources for the discovery and analysis of Tara Oceans data. Scientific Data, 2. doi:10.1038/sdata.2015.23

82. Potter, S. C., Luciani, A., Eddy, S. R., Park, Y., Lopez, R., & Finn, R. D. (2018). HMMER web server: 2018 update. Nucleic Acids Res, 46(W1), 200–204. doi:10.1093/nar/gky448

83. Radakovits, R., Jinkerson, R. E., Fuerstenberg, S. I., Tae, H., Settlage, R. E., Boore, J. L., & Posewitz, M. C. (2013). Draft genome sequence and genetic transformation of the oleaginous alga *Nannochloropsis gaditana*. Nature Communications, 4, 686. doi:10.1038/ncomms3356

84. Rastogi, A., Maheswari, U., Dorrell, R. G., Vieira, F. R. J., Maumus, F., Kustka, A., … Tirichine, L. (2018). Integrative analysis of large scale transcriptome data draws a comprehensive landscape of *Phaeodactylum tricornutum* genome and evolutionary origin of diatoms. Sci Rep, 8(1), 4834. doi:10.1038/s41598-018-23106-x

85. Raymond, J. A. (2011). Algal ice-binding proteins change the structure of sea ice. Proc Natl Acad Sci USA, 108(24), E198. doi:10.1073/pnas.1106288108

86. Raymond, J. A., & Kim, H. J. (2012). Possible role of horizontal gene transfer in the colonization of sea ice by algae. PLoS One, 7(5), e35968. doi:10.1371/journal.pone.0035968

87. Read, B. A., Kegel, J., Klute, M. J., Kuo, A., Lefebvre, S. C., Maumus, F., … Consortium, E. h. A. (2013). Pan genome of the phytoplankton *Emiliania* underpins its global distribution. Nature, 499(7457), 209–213. doi:10.1038/nature12221

88. Revell, L. J., & Graham Reynolds, R. (2012). A new Bayesian method for fitting evolutionary models to comparative data with intraspecific variation. Evolution, 66(9), 2697–2707. doi:10.1111/j.1558-5646.2012.01645.x

89. Rio, T. G. d., Harmon-Smith, M., Lucas, S. M., Copeland, A., Barry, K., Richardson, P., … Pangilinan, J. (2006). JGI Sequencing Projects-the process of ensuring efficiency and quality from initiation to completion.

90. Royo-Llonch, M., Sánchez, P., Ruiz-González, C., Salazar, G., Pedrós-Alió, C., Labadie, K., … Acinas, S. G. (2020). Ecogenomics of key prokaryotes in the arctic ocean. bioRxiv, 2020.2006.2019.156794. doi:10.1101/2020.06.19.156794

91. Sato, S., Nakamura, Y., Kaneko, T., Asamizu, E., & Tabata, S. (1999). Complete structure of the chloroplast genome of *Arabidopsis thaliana*. DNA Res, 6(5), 283–290. doi: 10.1093/dnares/6.5.283

92. Simão, F. A., Waterhouse, R. M., Ioannidis, P., Kriventseva, E. V., & Zdobnov, E. M. (2015). BUSCO: assessing genome assembly and annotation completeness with single-copy orthologs. Bioinformatics, 31(19), 3210–3212. doi:10.1093/bioinformatics/btv351

93. Song, Y., Liu, L., & Ma, X. (2019). CbAdh1 improves plant cold tolerance. Plant Signal Behav, 14(7), 1612680. doi:10.1080/15592324.2019.1612680

94. Stamatakis, A. (2014). RAxML version 8: a tool for phylogenetic analysis and post-analysis of large phylogenies. Bioinformatics, 30(9), 1312–1313. doi:10.1093/bioinformatics/btu033

95. Stephens, T. G., González-Pech, R. A., Cheng, Y., Mohamed, A. R., Burt, D. W., Bhattacharya, D., … Chan, C. X. (2020). Genomes of the dinoflagellate *Polarella glacialis* encode tandemly repeated single-exon genes with adaptive functions. BMC Biol, 18(1), 56. doi:10.1186/s12915-020-00782-8

96. Stephens, T. G., Ragan, M. A., Bhattacharya, D., & Chan, C. X. (2018). Core genes in diverse dinoflagellate lineages include a wealth of conserved dark genes with unknown functions. Sci Rep, 8(1), 17175. doi:10.1038/s41598-018-35620-z

97. Stewart, A., Rioux, D., Boyer, F., Gielly, L., Pompanon, F., Saillard, A., … Coissac, E. (2021). Altitudinal zonation of green algae biodiversity in the French Alps. Frontiers in Plant Science, 12(1066). doi:10.3389/fpls.2021.679428

98. Stiller, J. W., Schreiber, J., Yue, J., Guo, H., Ding, Q., & Huang, J. (2014). The evolution of photosynthesis in chromist algae through serial endosymbioses. Nat Commun, 5, 5764. doi:10.1038/ncomms6764

99. Strassert, J. F. H., Irisarri, I., Williams, T. A., & Burki, F. (2021). A molecular timescale for eukaryote evolution with implications for the origin of red algal-derived plastids. Nat Commun, 12(1), 1879. doi:10.1038/s41467-021-22044-z

100. Sunagawa, S., Coelho, L. P., Chaffron, S., Kultima, J. R., Labadie, K., Salazar, G., … coordinators, T. O. (2015). Ocean plankton. Structure and function of the global ocean microbiome. Science, 348(6237), 1261359. doi:10.1126/science.1261359

101. Suzek, B. E., Huang, H. Z., McGarvey, P., Mazumder, R., & Wu, C. H. (2007). UniRef: comprehensive and non-redundant UniProt reference clusters. Bioinformatics, 23(10), 1282–1288. doi:10.1093/bioinformatics/btm098

102. Søgaard, D. H., Sorrell, B. K., Sejr, M. K., Andersen, P., Rysgaard, S., Hansen, P. J., … Lund-Hansen, L. C. (2021). An under-ice bloom of mixotrophic haptophytes in low nutrient and freshwater-influenced Arctic waters Sci Rep,11, 2915. doi: 0.1038/s41598-021-82413-y

103. Terrado, R., Monier, A., Edgar, R., & Lovejoy, C. (2015). Diversity of nitrogen assimilation pathways among microbial photosynthetic eukaryotes. Journal of Phycology, 51(3), 490–506. doi:10.1111/jpy.12292

104. Turchetti, B., Thomas Hall, S. R., Connell, L. B., Branda, E., Buzzini, P., Theelen, B., … Boekhout, T. (2011). Psychrophilic yeasts from Antarctica and European glaciers: description of *Glaciozyma* gen. nov., Glaciozyma martinii sp. nov. and Glaciozyma watsonii sp. nov. Extremophiles, 15(5), 573–586. doi:10.1007/s00792-011-0388-x

105. Vancaester, E., Depuydt, T., Osuna-Cruz, C. M., & Vandepoele, K. (2020). Comprehensive and functional analysis of horizontal gene transfer events in diatoms. Mol Biol Evol, 37(11), 3243–3257. doi:10.1093/molbev/msaa182

106. Vance, T. D. R., Bayer-Giraldi, M., Davies, P. L., & Mangiagalli, M. (2019). Ice-binding proteins and the ‘domain of unknown function’ 3494 family. The FEBS Journal, 286(5), 855–873. doi:10.1111/febs.14764

107. Vaulot, D., Le Gall, F., Marie, D., Guillou, L., & Partensky, F. (2004). The Roscoff Culture Collection (RCC): a collection dedicated to marine picoplankton. Nova Hedwigia, 79(1-2), 32.

108. Vorobev, A., Dupouy, M., Carradec, Q., Delmont, T. O., Annamalé, A., Wincker, P., & Pelletier, E. (2020). Transcriptome reconstruction and functional analysis of eukaryotic marine plankton communities via high-throughput metagenomics and metatranscriptomics. Genome Res, 30(4), 647–659. doi:10.1101/gr.253070.119

109. Wang, D. M., Ning, K., Li, J., Hu, J. Q., Han, D. X., Wang, H., … Xu, J. (2014). *Nannochloropsis* genomes reveal evolution of microalgal oleaginous traits. Plos Genetics, 10(1). doi:10.1371/journal.pgen.1004094

110. Wang, H. C., Susko, E., & Roger, A. J. (2019). The relative importance of modeling site pattern heterogeneity versus partition-wise heterotachy in phylogenomic inference. Syst Biol, 68(6), 1003–1019. doi:10.1093/sysbio/syz021

111. Wang, Y., Pang, C., Li, X., Hu, Z., Lv, Z., Zheng, B., & Chen, P. (2017). Identification of tRNA nucleoside modification genes critical for stress response and development in rice and *Arabidopsis*. BMC Plant Biol, 17(1), 261. doi:10.1186/s12870-017-1206-0

112. Wheeler, D. L., Barrett, T., Benson, D. A., Bryant, S. H., Canese, K., Chetvernin, V., … Yaschenko, E. (2006). Database resources of the National Center for Biotechnology Information. Nucleic Acids Res, 34(Database issue), D173-180. doi:10.1093/nar/gkj158

113. Worden, A. Z., Lee, J. H., Mock, T., Rouze, P., Simmons, M. P., Aerts, A. L., … Grigoriev, I. V. (2009). Green evolution and dynamic adaptations revealed by genomes of the marine picoeukaryotes *Micromonas*. Science, 324(5924), 268–272. doi:10.1126/science.1167222

114. Zhang, X., Cvetkovska, M., Morgan-Kiss, R., Hüner, N. P. A., & Smith, D. R. (2021). Draft genome sequence of the Antarctic green alga *Chlamydomonas* sp. UWO241. iScience, 24(2), 102084. doi:10.1016/j.isci.2021.102084

115. Zhang, Z., Qu, C., Zhang, K., He, Y., Zhao, X., Yang, L., … Miao, J. (2020). Adaptation to extreme Antarctic environments revealed by the genome of a sea ice green alga. Curr Biol, 30(17), 3330–3341.e3337. doi:10.1016/j.cub.2020.06.029

